# An shRNA screen in primary human beta cells identifies the serotonin 1F receptor as a negative regulator of survival during transplant

**DOI:** 10.1101/2024.05.01.591950

**Authors:** Rebecca A. Lee, Deeksha G. Chopra, Vinh Nguyen, Xi-Ping Huang, Yaohuan Zhang, Kaavian Shariati, Nicholas Yiv, Rebecca Schugar, Justin Annes, Bryan Roth, Gregory M. Ku

## Abstract

Islet transplantation can cure type 1 diabetes, but peri-transplant beta cell death limits this procedure to those with low insulin requirements. Improving human beta cell survival or proliferation may make islet transplantation a possibility for more type 1 patients. To identify novel regulators of beta cell survival and proliferation, we conducted a pooled small hairpin RNA (shRNA) screen in primary human beta cells transplanted into immunocompromised mice. shRNAs targeting several cyclin dependent kinase inhibitors were enriched after transplant. Here, we focused on the Gi/o-coupled GPCR, serotonin 1F receptor (*HTR1F,* 5-HT_1F_) which our screen identified as a negative regulator of beta cell numbers after transplant. *In vitro*, 5-HT_1F_ knockdown induced human beta cell proliferation but only when combined with harmine and exendin-4. *In vivo*, knockdown of 5-HT_1F_ reduced beta cell death during transplant. To demonstrate the feasibility of targeting 5-HT_1F_ in islet transplant, we identified and validated a small molecule 5-HT_1F_ antagonist. This antagonist increased glucose stimulated insulin secretion from primary human islets and cAMP accumulation in primary human beta cells. Finally, the 5-HT_1F_ antagonist improved glycemia in marginal mass, human islet transplants into immunocompromised mice. We identify 5-HT_1F_ as a novel druggable target to improve human beta cell survival in the setting of islet transplantation.

**One Sentence Summary:** Serotonin 1F receptor (5-HT_1F_) negatively regulates insulin secretion and beta cell survival during transplant.

## INTRODUCTION

Type 1 diabetes is caused by an autoimmune attack on the pancreatic beta cells. The cure for type 1 diabetes must involve replacing or regenerating these critical cells (likely combined with immunosuppression). Indeed, type 1 diabetes can be cured by islet transplantation but many beta cells die soon after they are infused into the portal vein, meaning that only recipients with low insulin requirements can be cured with islets from a single donor(*1, 2*). Therefore, new therapeutics that improve human beta cell survival may make islet transplantation a possibility for more type 1 patients. While ES-cell derived beta cells may soon allow a virtually unlimited supply of beta-like cells, these cells also die in significant numbers upon transplant(*3*). Outside of the transplant setting, beta cell death is an important part of the pathophysiology of both type 1 and type 2 diabetes(*4*) and understanding human beta cell death may reveal new targets to treat both major forms of diabetes.

Increasing human beta cell replication would be a complementary approach to reducing beta cell death. Many studies of beta cell replication have identified candidates in rodent beta cells and then asked if these candidates can drive primary human beta cell replication. Unfortunately, while many new mouse beta cell mitogens have been found, most have proven ineffective in adult human beta cells (recently reviewed by (*5*)). Starting in rodent cells is not ideal since this strategy cannot identify any biology of primary human beta cells that is not conserved in rodents. Indeed, the biology of the human beta cell can be quite different than that of the mouse or rat, particularly with regard to proliferation(reviewed by(*6*)). A bright spot in the area of human beta cell replication has come from chemical screens(*7–10*). Interestingly, though these studies started with diverse libraries and/or screening strategies, most have converged on the inhibition of a single target, DYRK1A(*8, 9, 11, 12*). While the DYRK1A inhibitors have certainly been a game changer, a major gap in the field remains a relative paucity of other pathways that can trigger primary human beta cell replication.

To identify new regulators of human beta cell survival and proliferation, we performed a pooled shRNA screen in primary human beta cells. Our screen identified the serotonin G protein-coupled receptor (GPCR) 5-HT_1F_ as a novel regulator of human beta cell survival. We found that 5-HT_1F_ knockdown prevented human beta cell death, improved proliferation, and characterized a novel 5-HT_1F_ antagonist that could improve function and proliferation of human beta cells. Our data suggest that 5-HT_1F_ antagonists could be a novel therapeutic target to improve the efficiency of human islet transplant.

## RESULTS

### An *in vivo* shRNA screen in primary human beta cells identifies *HTR1F* (5-HT_1F_) as a negative regulator of human beta cell numbers after transplant

Since a tissue culture dish is not the native environment of the beta cell, we (and others) have reasoned that *in vivo* islet transplantation of human beta cells would allow for longer term and more physiologic studies of beta cell proliferation and survival(*13*). Therefore, we performed a pooled shRNA screen in primary human islets transplanted under the kidney capsule of immunocompromised mice. We infected primary human beta cells with a custom, pooled library of 12,472 independent shRNAs under the control of the insulin promoter (Figure 1A). A puromycin resistance gene was also expressed under the control of the same promoter to allow for selection of infected beta cells. Each of the 479 target genes had 25 independent shRNAs targeting it. Five hundred non-targeting shRNAs were included as negative controls. The targeted genes were chosen based on expression in primary human beta cells and annotation for either druggability or cell surface expression(*14, 15*). Genomic DNA was extracted from approximately half of the cells prior to transplant and the remaining cells were transplanted under the kidney capsule of an immunocompromised mouse. Four weeks after transplant, the graft was harvested, and genomic DNA was isolated. To address the likely small signal to noise ratio and to increase biological reproducibility, we performed our screen in biological triplicate, using islets from 3 independent donors.

**Fig. 1.**
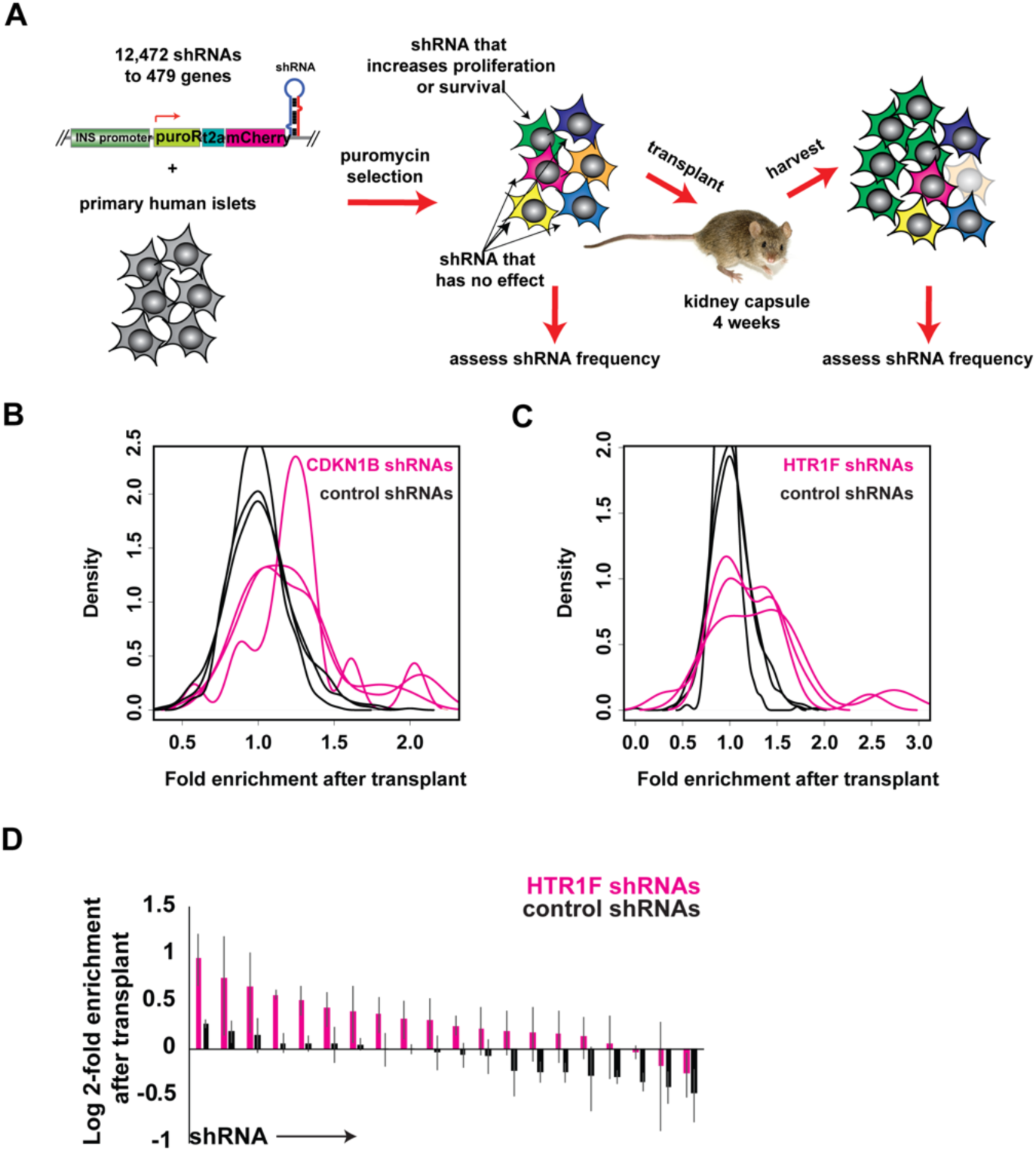
Pooled shRNA dropout screen in primary human beta cells. **(A)** Schema of screen design. See text for details. In green is a cell with an shRNA targeting a negative regulator of proliferation or survival. Note that there is a higher frequency of green cells after transplant. **(B)** Fold-enrichment after transplant of 25 shRNAs targeting *CDKN1B* (red) or 500 non-targeting shRNAs (black). Each line represents a different donor. A total of 3 donors are shown. **(C)** As in A, but for shRNAs targeting 5-HT_1F_ (pink). **(D)** Enrichments for each of the 5-HT_1F_ targeting shRNAs averaged over the 3 donors (red) and 20 randomly selected negative controls shRNAs (black). Note that these enrichments are represented as the log2 fold enrichment. Standard error is shown over the 3 independent donors.

Digital droplet PCR was used to estimate lentiviral copy number from the genomic DNA samples before and after the transplant. Sixty to eighty percent of lentiviral copies were lost after the transplant period, consistent with reported levels of beta cell loss after kidney capsule transplant(*3*). After transplant, the cell coverage (i.e. the number of lentiviral inserts recovered per unique shRNA in the original library) ranged between 50-500 (Data File S1, line 30). The frequency of each shRNA was then measured from the pre-transplant and post-transplant genomic DNA samples by next generation sequencing of the integrated lentiviral inserts. We elected to focus on shRNAs that became enriched after transplant as these are likely more specific (i.e. not simply causing increased cell death) and would potentially identify more feasible drug targets. A p-value for each gene was calculated based on the enrichments of all shRNAs targeting that gene compared to those of the 500 non-targeting shRNAs using a multiple testing corrected Mann-Whitney U test (Data File S2). We note that the screen that identified the lowest numbers of hits (transplant C) was also the lowest in terms of cell coverage.

shRNAs targeting two genes were statistically significantly enriched after transplant in all 3 donors (Data File S2). One gene was *CDKN1B* or p27Kip1, a cyclin dependent kinase (CDK) inhibitor. The enrichment ratios for non-targeting shRNAs (Figure 1B, black lines) were centered around 1 (no enrichment after transplant) while the *CDKN1B* shRNAs (Figure 1B, pink lines) had a tail to the right with 1.2-2-fold enrichment after transplant. Notably, there were some shRNAs to *CDKN1B* that did not become enriched. This was expected since many computationally predicted shRNAs do not knockdown the intended target and thus should not show enrichment. Knockdown or knockout of *CDKN1B* is known to increase human beta cell proliferation(*16*) and mouse beta cell proliferation(*17*), consistent with screen data. shRNAs to *CDKN2B*, another CDK inhibitor, were enriched after transplant in 2 of the 3 donors. Finding these two CDK inhibitors in our screen suggests that our screen method could detect negative regulators of human beta cell proliferation.

The second gene with statistically significant enriched shRNAs after transplant in all 3 donors was *HTR1F* (Figure 1C). The behavior of 20 of the 25 different *HTR1F* targeting shRNAs are shown in the pink lines -- the 5 other *HTR1F* shRNAs did not have enough reads to reliably quantitate. The same 500 non-targeting shRNAs shown in Figure 1B are again shown in black. As with *CDKN1B*, there were a subset of shRNAs to *HTR1F* that became enriched after transplant in all three donors (Figure 1C). Importantly, the same *HTR1F* shRNAs were enriched after transplant in all 3 donors (Figure 1D).

*HTR1F* (5-HT_1F_) is a G_i/o_-coupled, class A GPCR(*18, 19*) whose ligand, serotonin, is released from pancreatic beta cells in a glucose-dependent fashion, suggesting a possible autocrine or paracrine loop(*20*). In mice, serotonin plays an important role in beta cell proliferation during pregnancy, lactation, and the perinatal period through the Gaq-coupled 5-HT_2B_ receptor(*21–23*). Serotonin also increases glucose stimulated insulin secretion (GSIS) through *HTR2B* in adult mice and human islets(*24*). In pregnancy, serotonin increases GSIS through *Htr3a*(*25*). 5-HT_1F_ in the alpha cell negatively regulates glucagon secretion in a paracrine response to serotonin release from beta cells(*26*), but there is no known role for 5-HT_1F_ in the beta cell. Previously published mRNA-seq data shows that 5-HT_1F_ is also expressed in sorted primary human beta cells at 5.3 transcripts per million (TPM) (*14, 27*), making it the most highly expressed serotonin GPCR by 10-fold in the primary human beta cell (Fig. S1A). Previously published single cell RNA-seq also confirms that 5-HT_1F_ is expressed in the human beta cell, alpha cell, delta cell, and gamma cell but not in the acinar cell(*28, 29*) (Fig. S1B). Notably, while 5-HT_1F_ is among the most highly expressed GPCRs in human islets, it is not expressed in the mouse or rat islet (*30, 31*) – highlighting the importance of primary screening in human beta cells.

### 5-HT1F knockdown increases human beta cell proliferation in combination with harmine and exendin-4

One possibility is that 5-HT_1F_ silencing increases human beta cell proliferation. To test this, we infected dissociated human islets with lentivirus expressing a validated 5-HT_1F_ shRNA or control shRNA (expressing nuclear GFP under the control of the insulin promoter) (knockdown validated in Fig. S2A). The cells were then treated with DMSO, 5 nM GLP1R agonist exendin-4, 10 μM harmine, or exendin-4 and harmine (*32*). Cells were then stained for GFP, insulin, and KI67 to assess proliferation. Treatment with exendin-4 and harmine increased the proliferation of human beta cells and this was further increased by 5-HT_1F_ knockdown (Fig. 2A). In contrast, GFP negative insulin positive cells had similar proliferation rates regardless of the shRNA (Fig. 2B). This was expected since GFP negative cells do not express any shRNA. However, 5-HT_1F_ knockdown alone did not increase proliferation, suggesting that increased proliferation was not likely to explain why 5-HT_1F_ shRNAs were enriched after transplant.

**Fig. 2.**
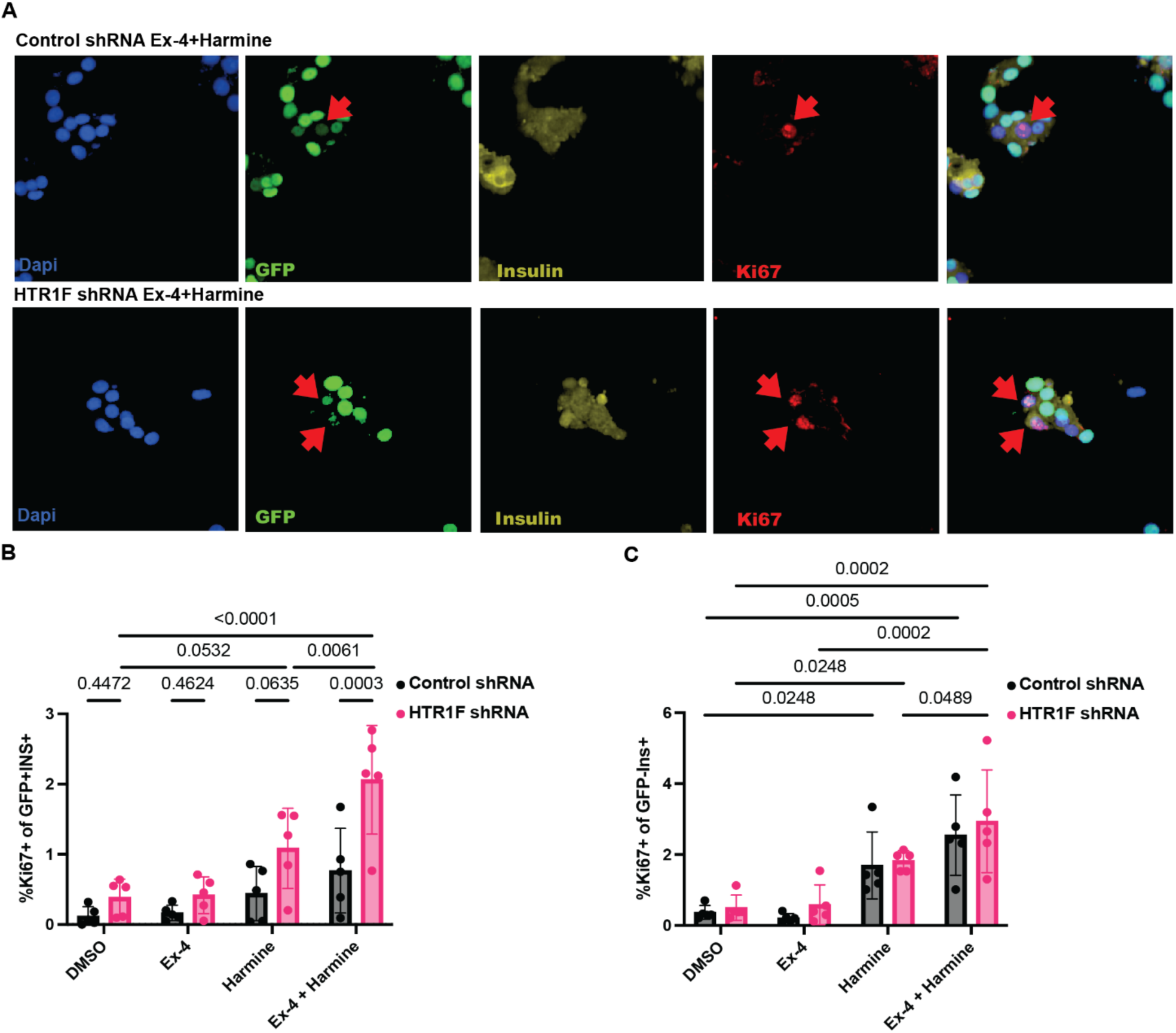
5-HT_1F_ knockdown increases human beta cell proliferation but only in the setting of harmine and exedin-4. **(A)** Human islets infected with control or 5-HT_1F_ shRNA and treated with harmine and exendin-4 and then stained for DNA, GFP, insulin, and KI67. Red arrows indicate a GFP+Insulin+KI67+ cell. **(B)** The percent of the GFP+INS+ cells that were also KI67 is shown. n=5 donors. 2-way ANOVA: p=0.0001 for shRNA effect, p<0.0001 for drug effect, p=0.049 for interaction. **(C)** As in A, but the percent of GFP negative (uninfected), INS+ cells that are also KI67+ is shown. 2-way ANOVA: p=0.16 for shRNA effect, p<0.0001 for drug effect. The interaction was not significant. Error bars show standard error. Post-hoc testing with Benjamini-Hochberg correction. Only p-values <0.05 are shown for clarity (out of 28 tests).

### 5-HT1F knockdown prevents human beta cell death during transplant

We next asked if 5-HT_1F_ knockdown could reduce human beta cell death after transplant. To reduce variability between transplants and between donors, we infected intact primary human islets with either control shRNA or 5-HT_1F_ shRNA lentiviruses (Fig. 3A), pooled the infected islets, and transplanted the mixture into a single mouse recipient. This approach allows comparison of two conditions (5-HT_1F_ knockdown or control knockdown) in the same animal, allowing us to control for variability between different transplants. To distinguish between the control shRNA expressing beta cells and 5-HT_1F_ shRNA expressing beta cells, the control shRNA was co-expressed with nuclear GFP and the 5-HT_1F_ shRNA was co-expressed with nuclear mCherry. The grafts were harvested 4 days after transplant as we reasoned that most death occurred in the immediate post-transplant period. We measured the frequency of cleaved caspase-3+ cells in the GFP+ population (control knockdown) and compared it to the frequency of cleaved caspase-3+ cells in the mCherry+ population (5-HT_1F_ knockdown). Over 4 independent donors (4 independent transplants), we found that the frequency of cell death in the 5-HT_1F_ knockdown cells was reduced by 30% as compared to control knockdown (Fig. 3B and C). To rule out an effect of the fluorescent protein, we paired the control shRNA with mCherry and the 5-HT_1F_ knockdown with GFP and reproduced a reduction in cell death in 5-HT_1F_ knockdown beta cells in a 5^th^ donor (Fig. 3D). A second, independent shRNA targeting 5-HT_1F_ (validated in Fig. S2A) also reduced beta cell death over the non-targeting shRNA (Fig. 3E).

**Fig. 3.**
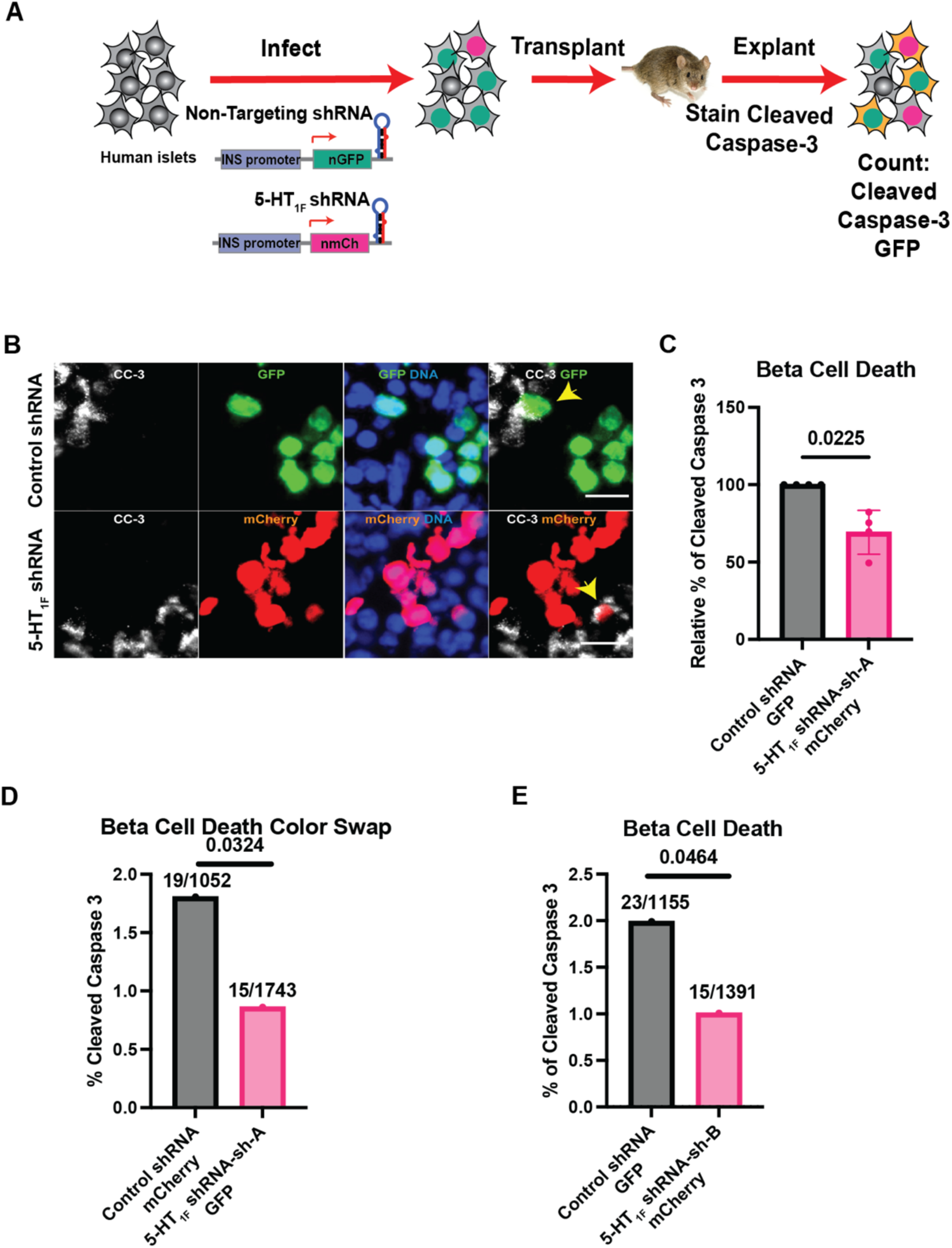
5-HT_1F_ knockdown reduces beta cell death after transplant. **(A)** Strategy to measure beta cell death in the setting of transplant (see text for details). **(B)** Human islets infected with control shRNA lentivirus (GFP) or 5-HT_1F_ shRNA (sh-A) lentivirus (mCherry), transplanted, and stained for cleaved caspase-3, mCherry, and GFP as described in A. **(C)** The % reduction in cleaved caspase-3 in mCherry positive cells (5-HT_1F_ knockdown for sh-A shRNA) as compared to the GFP positive cells (control knockdown), n=4 donors. ** p<0.01 by one sample, two-tailed Student’s t-test. Error bars show standard error. **(D)** As in (B), but the 5-HT_1F_ knockdown, sh-A is paired with mCherry and the non-targeting shRNA is paired with GFP. The % of cleaved caspase-3 in GFP positive cells (5-HT_1F_ knockdown, sh-A) and mCherry positive cells (control knockdown) is plotted, **(E)** The % of cleaved caspase-3 in mCherry positive cells (5-HT_1F_ knockdown for shRNA B, sh-B) and GFP positive cells (control knockdown). For both D and E, n=1 donor, p-value from Fisher’s exact test. Numbers above each bar are the number of CC-3+fluorescent marker+ cells over the total number of fluorescent marker+ cells counted.

### Identification and validation of a small molecule antagonist of 5-HT_1F_

Although there are no published, specific 5-HT_1F_ antagonists, we did find a series of small molecule antagonists for 5-HT_1F_ described in a patent as potential anti-anxiety agents (*33*). To validate one of these small molecule 5-HT_1F_ antagonists, we turned to a luciferase-based biosensor for cAMP (GloSensor, Promega) to measure G_i/o_ antagonist activity in HEK293T cells. Briefly, HEK293T cells were transfected with the cAMP GloSensor with or without 5-HT_1F_ cDNA and treated with serotonin (5-HT) and the potential 5-HT_1F_ antagonist. Without 5-HT_1F_ transfection, serotonin had no G_i/o_ agonist activity and the putative 5-HT_1F_ antagonist, 1-(2-hydroxy-3-(naphthalen-2-yloxy)propyl)-4-(quinolin-3-yl)piperidin-4-ol (Fig, 4A) had non-specific inhibitory effects on luminescence at concentrations ≥3 uM (Fig. S3A). Expression of 5-HT_1F_ conferred an inhibitory response to serotonin with an EC_50_ of 0.1 nM (Fig. S3B), consistent with the reported Gi/o coupling of 5-HT_1F_ and similar to its reported Ki for serotonin (*19, 34*). Similarly, 5-HT showed potent (2.3 nM EC_50_) G_i/o_-agonist activity at 5-HT_1A_ (Fig. S3B). The putative 5-HT_1F_ antagonist, 1-(2-hydroxy-3-(naphthalen-2-yloxy)propyl)-4-(quinolin-3-yl)piperidin-4-ol, showed no G_i/o_-agonist activity, and again non-specifically reduced luminescence signals at concentrations of ≥3 uM (Fig. S3B) at 5-HT_1F_ and 5-HT_1A_ cells as in control cells. Furthermore, 1-(2-hydroxy-3-(naphthalen-2-yloxy)propyl)-4-(quinolin-3-yl)piperidin-4-ol had no G_s_-agonist activity to increase cAMP production, while isoproterenol stimulation led to a >40-fold increase in cAMP production (Fig. S3C). In contrast, 1-(2-hydroxy-3-(naphthalen-2-yloxy)propyl)-4-(quinolin-3-yl)piperidin-4-ol potently inhibited 5-HT_1F_ with a Ki of 47 nM in the presence of 10 nM 5-HT, but did not inhibit 5-HT_1A_ (Fig. 4B). We used a radioligand binding assay with 3H-LSD to determine compound binding affinity to 5-HT_1F_ and found that 1-(2-hydroxy-3-(naphthalen-2-yloxy)propyl)-4-(quinolin-3-yl)piperidin-4-ol had a Ki of 11 nM, similar to other known ligands of 5-HT_1F_ (Fig. 4C). Among all other serotonin GPCRs, the next highest binding membrane binding affinity was for 5-HT_2B_ with a Ki of 343 nM (Fig. 4D). These data show that 1-(2-hydroxy-3-(naphthalen-2-yloxy)propyl)-4-(quinolin-3-yl)piperidin-4-ol is a specific 5-HT_1F_ antagonist.

**Fig. 4.**
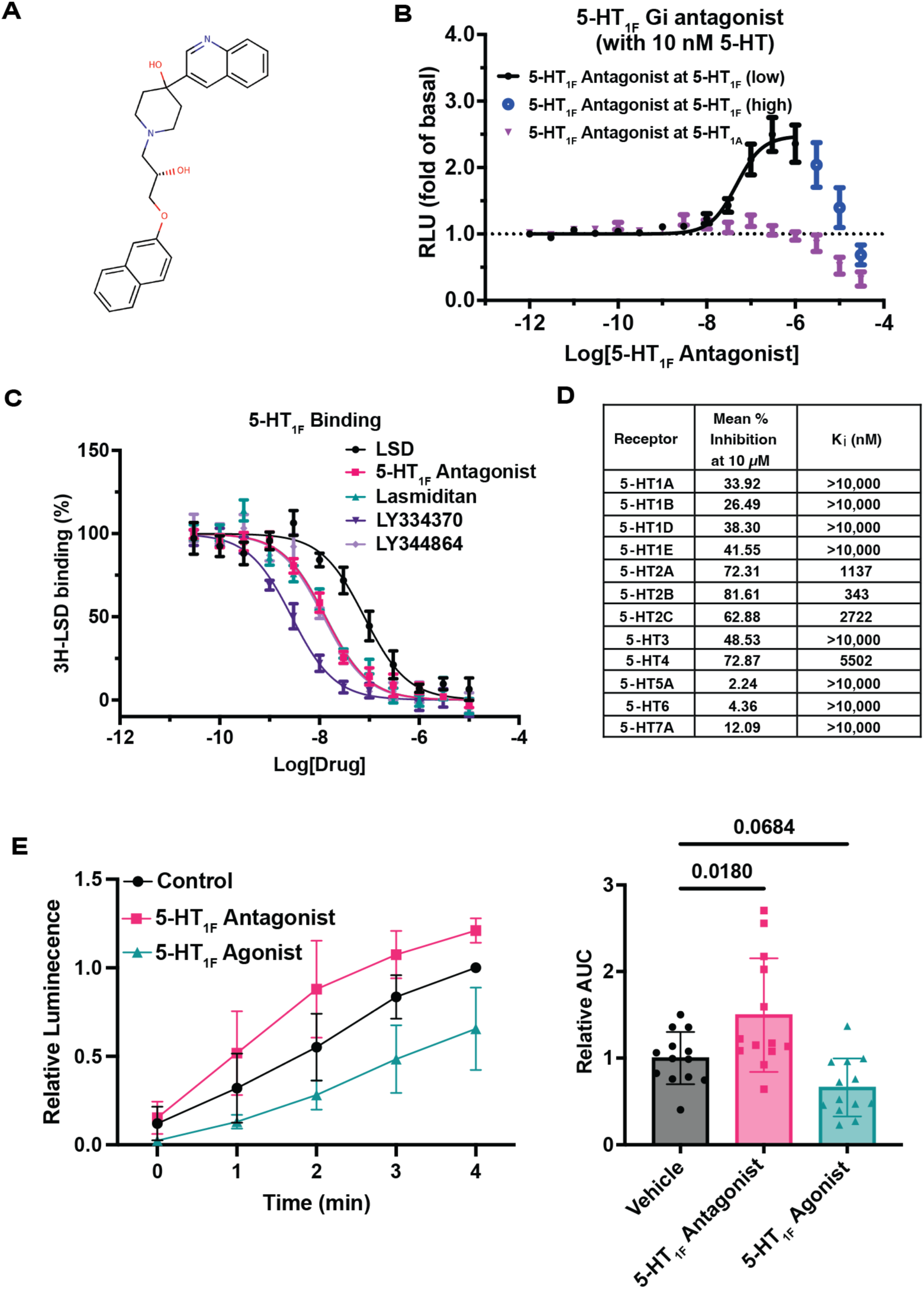
Validation of a small molecule 5-HT_1F_ antagonist. **(A)** Structure of 5-HT_1F_ antagonist 1-(2-hydroxy-3-(naphthalen-2-yloxy)propyl)-4-(quinolin-3-yl)piperidin-4-ol. **(B)** 293T cells were transiently transfected with the GloSensor plasmid (Promega) and 5-HT_1F_ or 5-HT_1A_ cDNA. Cell were incubated with indicated concentrations of 1-(2-hydroxy-3-(naphthalen-2-yloxy)propyl)-4-(quinolin-3-yl)piperidin-4-ol (x-axis), followed by 10 nM 5-HT and isoproterenol. Fold change from of baseline luminescence is plotted. Error bars show standard error from n=3 independent replicates, each in quadruplicate. Because of a non-specific inhibitory effect on luminescence, antagonist results above 3 uM were separated as Antagonist at 5-HT1F (high). **(C)** 5-HT_1F_ radioligand binding assay using ^3^H-LSD to determine compound binding affinity to 5-HT_1F_, Error bars indicate standard error from n=3 independent replicates, each in triplicate. **(D)** Binding affinity (Ki) determined by competitive binding to 5-HT receptors, LSD (64 nM), 1-(2-hydroxy-3-(naphthalen-2-yloxy)propyl)-4-(quinolin-3-yl)piperidin-4-ol (11 nM), Lasmiditan (11 nM), LY334370 (2.2 nM), LY334864 (9.6 nM), from 3 independent assays. (**E)** Human islets were infected with an adenovirus expressing the GloSensor cAMP responsive luciferase under the control of the insulin promoter. Forty-eight hours later, they were incubated in HBSS with 1% GloSensor reagent. 1-(2-hydroxy-3-(naphthalen-2-yloxy)propyl)-4-(quinolin-3-yl)piperidin-4-ol 300nM or agonist 100nM was added at time −5 min. 1μM of forskolin was added at time 0 and luminescence was measured each minute following. Values were normalized to vehicle luminescence at 4 minutes (left panel) or the area under the curve of the vehicle treatment (right panel). n=3 donors with 4-5 replicates per donor. One-way ANOVA with Benjamini-Hochberg correction. Error bars show standard error.

### 5-HT1F negatively regulates cAMP in human beta cells

To validate these findings in human beta cells, we generated an adenovirus expressing a cAMP activated luciferase (GloSensor, Promega) under the control of the insulin promoter. We infected intact human islets with this adenovirus and measured luminescence after treatment with the 5-HT_1F_ agonist LY344864 or 1-(2-hydroxy-3-(naphthalen-2-yloxy)propyl)-4-(quinolin-3-yl)piperidin-4-ol or vehicle. 1-(2-hydroxy-3-(naphthalen-2-yloxy)propyl)-4-(quinolin-3-yl)piperidin-4-ol increased the integrated cAMP signal over the vehicle control (Fig. 4E). In contrast, treatment with the 5-HT_1F_ agonist, LY344865, reduced integrated cAMP signal though this did not meet our threshold for significance (Fig. 4E). These data suggest that 5-HT_1F_ also reduces cAMP in human beta cells, consistent with G_i/o_ coupling.

### 5-HT1F negatively regulates glucose stimulated insulin secretion in human beta cells in vitro

Since 5-HT_1F_ appears to be G_i/o_-coupled in human beta cells, we predicted that it would negatively regulate insulin secretion. Indeed, we found that the 5-HT_1F_ agonist, LY344864, reduced glucose stimulated insulin secretion in islets from 8 human donors (Fig. S4). In contrast, 1-(2-hydroxy-3-(naphthalen-2-yloxy)propyl)-4-(quinolin-3-yl)piperidin-4-ol increased insulin secretion at 11 mM glucose and was trending to increase insulin secretion at the basal 2.8 mM glucose (Fig. 5A) over 9 human donors. 5-HT_1F_ was previously reported to negatively regulate glucagon secretion from alpha cells (*26*). Confirming this study, we found that the 5-HT_1F_ antagonist increased glucagon secretion from human islets (Fig. 5B).

**Fig. 5.**
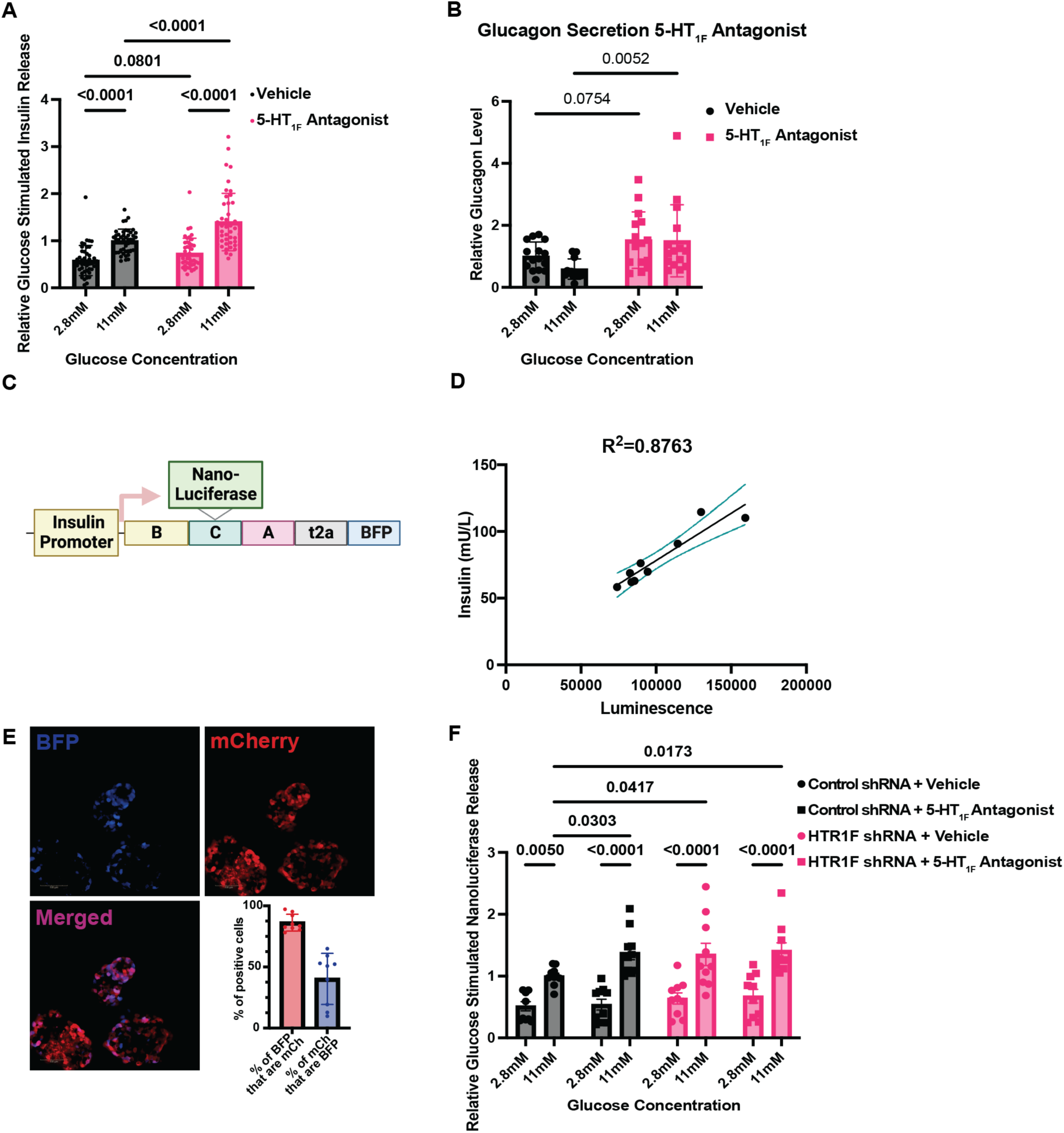
5-HT_1F_ negatively regulates insulin secretion. **(A)** Static glucose stimulated insulin secretion in presence of vehicle or 150nM of 1-(2-hydroxy-3-(naphthalen-2-yloxy)propyl)-4-(quinolin-3-yl)piperidin-4-ol. n=9 donors, 5 replicates for each donor. 2-way ANOVA: drug effect p<0.0001, glucose p<0.0001, interaction p=0.038. Post-hoc test p-values are Benjamini-Hochberg corrected. **(B)** Glucagon levels measured from a subset of (A), n=3 donors, 5 replicates for each donor. 2-way ANOVA: drug effect p<0.001, glucose p=0.2925, interaction p=0.3461. Post-hoc test p-values are Benjamini-Hochberg corrected. **(C)** Schematic of an insulin nanoluciferase BFP adenovirus to monitor insulin secretion from human beta cells. B = B chain of insulin, A = A chain of insulin, C = C-peptide. T2A = ribosomal skip sequence. BFP = blue fluorescent protein. **(D)** Correlation between insulin determined by ELISA and nanoluciferase luminescence from secretion samples of human islets infected with the insulin nanoluciferase reporter adenovirus shown in part C. n=9 independent samples from one human donor. **(E)** Image of human islets infected with adenovirus expressing control or 5-HT_1F_ shRNA mCherry co-infected with the insulin nanoluciferase BFP adenovirus and percent infection of co-infection, n=2 donors, 3-6 replicates for each donor **(F)** Human islets infected with the insulin nanoluciferase BFP adenovirus (MOI 50) and the 5-HT_1F_ shRNA or control shRNA adenovirus (MOI 200). 72 hours later, GSIS was performed. n=2 donors with 5 replicates for each donor. 3 way ANOVA 1-(2-hydroxy-3-(naphthalen-2-yloxy)propyl)-4-(quinolin-3-yl)piperidin-4-ol effect p=0.1077, glucose p<0.0001, and 5-HT_1F_ shRNA p=0.0392. Post-hoc test p-values were Benjamini-Hochberg FDR corrected. Error bars show standard error.

To rule out off targets effects of 1-(2-hydroxy-3-(naphthalen-2-yloxy)propyl)-4-(quinolin-3-yl)piperidin-4-ol, we assessed its ability to increase insulin secretion in 5-HT_1F_ knockdown cells. We knocked down 5-HT_1F_ in human islets using an adenovirus expressing an shRNA targeting 5-HT_1F_ and mCherry driven by the CMV promoter (validation shown in Fig. S2B). However, we were unable to attain infection of the core of intact human islets, most likely due to poor access by the adenovirus. Therefore, we co-infected with a adenovirus that expressed a copy of the insulin cDNA along with nanoluciferase inserted into C-peptide as previously reported (*35*) with BFP, both driven by the rat insulin promoter (Fig. 5C). To validate this construct, we infected human islets and 4 days later measured luciferase activity and insulin levels from the same secretion samples. We found that nanoluciferase activity was well correlated with insulin measured by ELISA (Fig. 5D). We then infected human islets with this insulin nanoluciferase BFP adenovirus (MOI 50) along with an excess (MOI 200) of the adenovirus expressing control or 5-HT_1F_ targeting shRNA to ensure that most cells with knockdown of 5-HT_1F_ expressed the nanoluciferase reporter. Indeed, 86% of BFP infected cells also expressed mCherry (Fig. 5E), showing that under these infection conditions, luciferase secretion predominantly comes from cells that express the shRNA. We found that 1-(2-hydroxy-3-(naphthalen-2-yloxy)propyl)-4-(quinolin-3-yl)piperidin-4-ol increased luciferase secretion at high glucose (Fig. 5F, bar 4) as we previously observed in Figure 5A as did 5-HT_1F_ knockdown (Fig. 5F, bar 6). However, combining 5-HT_1F_ knockdown with 1-(2-hydroxy-3-(naphthalen-2-yloxy)propyl)-4-(quinolin-3-yl)piperidin-4-ol (Fig. 5F, bar 8) did not further increase GSIS above either alone, showing that 1-(2-hydroxy-3-(naphthalen-2-yloxy)propyl)-4-(quinolin-3-yl)piperidin-4-ol’s effect on insulin secretion is dependent on 5-HT_1F_ expression.

### 5-HT1F antagonism modestly increases human beta cell replication in combination with harmine and exendin-4

Since knockdown of 5-HT_1F_ increased human beta cell replication in combination with harmine and exendin-4, we asked if 5-HT_1F_ antagonism would do the same. We treated dissociated islets with 1-(2-hydroxy-3-(naphthalen-2-yloxy)propyl)-4-(quinolin-3-yl)piperidin-4-ol or vehicle in combination with harmine and exendin-4. We found that 1-(2-hydroxy-3-(naphthalen-2-yloxy)propyl)-4-(quinolin-3-yl)piperidin-4-ol increased human beta cell replication in combination with harmine and exendin-4 (Fig. S5) but did not do so on its own.

### 5-HT1F antagonist improves glycemia after human islet transplant in diabetic mice

Our studies suggest 5-HT_1F_ antagonism could be used to improve human islet transplant. We found that a single intraperitoneal (IP) injection of 20 mg/kg in NOD.Cg-Prkdc^scid^Il2rg^tm1Wjl^/Sz (NSG) mice resulted in plasma levels >100 nM for approximately 8 hours (Fig. S6A). To generate diabetic recipients that would not reject a human islet transplant, diabetes was induced in NSG mice with STZ injections. Then, human islets were pre-treated with 200 nM of 5-HT_1F_ antagonist overnight prior to transplantation. Two thousand IEq were transplanted under the kidney capsule of these diabetic mice followed by twice daily IP injections of 20 mg/kg of 1-(2-hydroxy-3-(naphthalen-2-yloxy)propyl)-4-(quinolin-3-yl)piperidin-4-ol or a vehicle control (Fig. 6A). Over 3 separate human donors, we found that 1-(2-hydroxy-3-(naphthalen-2-yloxy)propyl)-4-(quinolin-3-yl)piperidin-4-ol did not change non-fasting blood glucose levels or body weight following transplant (Fig. S6B-C). However, the more sensitive intraperitoneal glucose tolerance test was improved in 1-(2-hydroxy-3-(naphthalen-2-yloxy)propyl)-4-(quinolin-3-yl)piperidin-4-ol treated mice (Fig. 6B). Since glucose levels were lower in the antagonist treated animals, we normalized fasting plasma insulin to fasting glucose. The antagonist treated mice had higher normalized insulin levels compared to vehicle treated mice (Fig. 6C). Following the glucose tolerance test, grafts were removed, and all mice reverted to blood glucose levels >350 mg/dl (Fig. 6D).

**Fig. 6.**
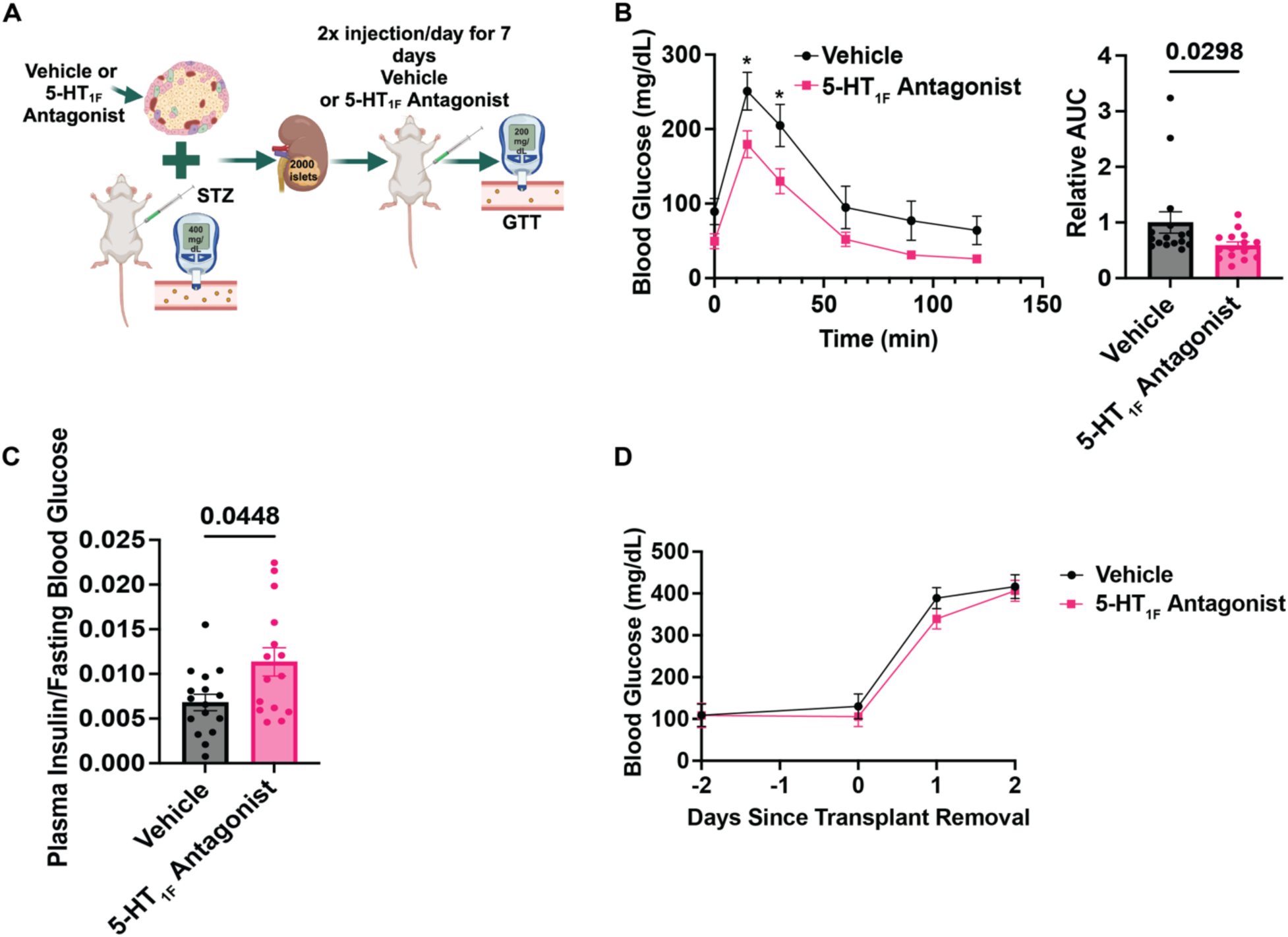
Chemical inhibition of 5-HT_1F_ improves glycemia after human islet transplant. **(A)** Schema of marginal mass islet transplant model. Human islets from were pretreated with 200 nM of 1-(2-hydroxy-3-(naphthalen-2-yloxy)propyl)-4-(quinolin-3-yl)piperidin-4-ol or vehicle and then transplanted under the kidney capsule of diabetic male NSG mice. Mice received 20 mg/kg of 1-(2-hydroxy-3-(naphthalen-2-yloxy)propyl)-4-(quinolin-3-yl)piperidin-4-ol or vehicle by IP injection every 12 hours. **(B)** Intraperitoneal glucose tolerance test on mice at day 7 after transplant. Two-way ANOVA: drug effect p=0.0432, time p<0.0001. Interaction p=0.25. * denote timepoints where p<0.05 post-hoc test p-values are Benjamini-Hochberg FDR corrected. Relative area under the curve was calculated, p=0.0298 Mann-Whitney test. **(C)** Plasma insulin normalized to fasting blood glucose, p=0.0448 Mann-Whitney test, from a total of 3 different donors, vehicle n=16 mice, 5-HT_1F_ antagonist n=15 mice. **(D)** Non-fasting plasma glucose levels prior to and following transplant removal n=12 vehicle mice, n=10 1-(2-hydroxy-3-(naphthalen-2-yloxy)propyl)-4-(quinolin-3-yl)piperidin-4-ol treated mice from 2 independent donors. Two-way ANOVA: drug effect p=0.5068, time p<0.0001. Interaction p=0.5490. Error bars show standard error.

Although *Htr1f* is not expressed in the mouse islet, it is expressed in the brain and peripheral blood. To rule out a non-islet effect of 1-(2-hydroxy-3-(naphthalen-2-yloxy)propyl)-4-(quinolin-3-yl)piperidin-4-ol on glucose homeostasis, we treated non-diabetic, non-transplanted NSG mice with 1 week of 1-(2-hydroxy-3-(naphthalen-2-yloxy)propyl)-4-(quinolin-3-yl)piperidin-4-ol given IP twice per day. The 5-HT_1F_ antagonist did not change non-fasting blood glucose, bodyweight, glucose tolerance, or insulin tolerance (Fig. S7A-D). Finally, since 1-(2-hydroxy-3-(naphthalen-2-yloxy)propyl)-4-(quinolin-3-yl)piperidin-4-ol can increase insulin secretion acutely in human islets *in vitro*, the effect of 5-HT_1F_ antagonism on glucose tolerance after transplant could be due to an effect on insulin secretion and not an effect on survival. Therefore, we asked if 1-(2-hydroxy-3-(naphthalen-2-yloxy)propyl)-4-(quinolin-3-yl)piperidin-4-ol could increase insulin secretion acutely during glucose tolerance testing. We found no change in glucose tolerance in mice with stably engrafted human islets after 1-(2-hydroxy-3-(naphthalen-2-yloxy)propyl)-4-(quinolin-3-yl)piperidin-4-ol treatment (Fig. S8), suggesting the improvement in glucose tolerance in our acute transplant model is mediated by beta cell survival, not by increasing insulin secretion per beta cell.

## DISCUSSION

Using a pooled shRNA screen in primary human beta cells, we uncovered a role for 5-HT_1F_ during beta cell transplant. While our pooled shRNA screen was successfully able to identify negative regulators of beta cell survival and proliferation and identified 5-HT_1F_, the enrichment of shRNAs after transplant was relatively mild. This was expected. If there was beta cell proliferation, enrichments were expected to be modest because not all cells with knockdown of a gene would likely proliferate. In the case of DYRK1A inhibition, arguably the gold standard to trigger human beta cell replication, only a small fraction of treated cells actually enter the cell cycle(*7*). If 50% of cells with knockdown of a gene divide twice, the expected enrichment of that shRNA would be 2-fold. If there was prevention of beta cell death, the maximum enrichment is expected to be inversely proportional to the fraction of cells that die after transplant. In our case, we estimated enrichment being capped at approximately 5-fold assuming that knockdown of the gene of interest protects 100% of the cells from death and given 60-80% cell death after transplant. In contrast, dropout screens in cell lines are often capable of generating thousands of fold enrichments or depletions due to ongoing death and proliferation(*36*). Future dropout screens in human beta cells might be optimized to improve the signal from proliferation or survival.

While 5-HT_1F_ negatively regulates glucagon secretion from alpha cells(*26*), it had no known role in beta cells. In our shRNA screen, 5-HT_1F_ knockdown cells were more abundant after transplant. We believe that this is primarily due to reduced cell death and not proliferation since 5-HT_1F_ knockdown or antagonisms was only able to increase proliferation in the setting of harmine and exendin-4 treatment.

The fact that antagonism or knockdown of 5-HT_1F_ can increase insulin secretion, cell survival, and proliferation in vitro is consistent with serotonin production within human islets. Indeed, serotonin is secreted from human beta cells themselves (*26*). This suggests that 5-HT_1F_ not only participates in a paracrine inhibition of alpha cell glucagon release, but also an autocrine feedback inhibition on beta cells. We note that endogenous serotonin does not appear to be saturating in isolated human islets in vitro as an 5-HT_1F_ agonist was still able to reduce insulin secretion and cAMP levels in human beta cells.

In our human islet transplant model, inhibition of 5-HT_1F_ improved glucose tolerance. We took advantage of the fact that 5-HT_1F_ is not expressed in the mouse islet to demonstrate that the effect of the 5-HT_1F_ antagonist was not due to effects of the antagonist on the donor mouse. Indeed, the 5-HT_1F_ antagonist had no effect on mice that did not receive an islet transplant in terms of body weight, glucose tolerance or insulin, showing that the 5-HT_1F_ antagonist requires human islets to improve blood glucose. These data strongly suggest that the effects of the 5-HT_1F_ antagonist are not due to changes in insulin sensitivity of the recipient mouse.

We believe improved glucose tolerance in 5-HT_1F_ antagonist treated mice during the peri-transplant period is most likely due to an effect on beta cell survival because 5-HT_1F_ antagonism in mice with stably engrafted human islets did not show improved glucose tolerance with 5-HT_1F_ antagonist treatment. While this appears to contradict the increased insulin secretion from human islets treated with 5-HT_1F_ antagonist *in vitro*, the lack of an *in vivo* glucose lowering effect might come from a concomitant increase in glucagon release, which could offset the glucose lowering effect of increased insulin secretion. Nonetheless, our data suggests that 5-HT_1F_ antagonists might be useful for human islet transplants to reduce cell loss immediately after transplant.

We found that 5-HT_1F_ antagonism increased glucagon secretion from human islets, in agreement with previous work and providing another piece of evidence that the 5-HT_1F_ antagonist is specific (*26*). However, with respect to its effect on insulin secretion from beta cells, we cannot exclude an effect from intra-islet communication. Since glucagon can increase insulin secretion from beta cells (*37*), it is possible that some of the increase in insulin secretion during 5-HT_1F_ antagonism comes from increased glucagon release from alpha cells. Furthermore, since delta cell also express 5-HT_1F_, it is possible that there is altered somatostatin secretion after 5-HT_1F_ antagonism. Cell type specific targeting of the antagonist or cell type specific knockdown of 5-HT_1F_ could help resolve these questions.

Together, we identify 5-HT_1F_ as a novel negative regulator of human beta cell survival and function and provide pre-clinical data supporting the use 5-HT_1F_ antagonists to improve human islet transplant.

## MATERIALS AND METHODS

### Study Design

Sample sizes were determined via power analysis and previous experiments using human islet donors. For mouse experiments, data collection was stopped, and mice were euthanized in consultation with UCSF veterinarian staff if body condition scoring worsened and if blood glucose levels following human islet transplant were >350 mg/dL for two consecutive days. All data acquired for these experiments was included in the manuscript. Mouse experiment endpoints were determined by preliminary studies as shown in the supplementary figures where we determined that healthy mice could be dosed for up to 7 days (2x daily) without any observed changes in body weight, random blood glucose levels, nor glucose/insulin tolerance. Each experiment was performed the number of times stated in the figure legend and replicates were determined by past experiments and availability of donors. The research objectives of this work were to identify and interrogate new targets involved in human beta cell survival and/or proliferation. We hypothesized that we could identify new targets using a pooled shRNA screen and then look for molecular tools to target hits identified in the screen. Anonymized human islet donor samples and mice were used as described below. Mice were evenly distributed into experimental groups by age, weight, and blood glucose level before transplant and were housed either individually or in treatment groups. Data was collected and processed both randomly and grouped by treatment. The surgeon performing the transplants and graft collection was blinded to the treatment groups. Images and data were analyzed in groups blinded to the intervention when possible, such as during image analysis and data analysis.

### Pooled shRNA library construction

A pSicoR-based lentiviral vector expressing puromycin resistance T2A mCherry under the control of the insulin promoter was a generous gift from Sergio Covarubius and Michael T. McManus (pSC2). A custom pool of DNA oligos encoding 12,472 unique shRNAs was synthesized (CustomArray), amplified by PCR (15 cycles, Phusion, NEB) and cloned into the XhoI and EcoRI sites of pSC2 as previously described(*38*) with the modification that the original EcoRI site was restored back to the native mir-30 sequence to improve shRNA processing (*39*).

### Human islet transplant screen

Human islets (20-30K IEQ) were dissociated the day after isolation with collagenase P (Roche) digestion for 15 minutes at 37 ℃, 0.05% trypsin (Thermo)for 10 minutes, and finally gentle trituration to dissociate the islets into single cells. Trypsin was inactivated with fetal bovine serum and the cells were plated on 804G coated plates in CMRL1066 (MediaTech) + B-27 supplement (Thermo), Glutamax (Thermo), non-essential amino acids (Thermo), penicillin and streptomycin. The cells were infected with lentivirus containing the shRNA pool at MOI 3-5. Puromycin (3 ug/mL final) was added 3 days after infection and 60% of the surviving cells were transplanted under the kidney capsule of scd-beige (Charles River) mice at day 7-9 after infection while 40% were frozen for genomic DNA isolation and lentiviral insert sequencing. Four weeks after transplant, the graft was removed and genomic DNA from both samples were isolated. Lentiviral copy number was measured using digital droplet PCR (QX100, Biorad) using a probe recognizing the lentiviral backbone RRE sequence. RRE-F= GGCAAAGAGAAGAGTGGTGC; RRE-R= GACGGTACAGGCCAGACAAT; RRE-probe = CCATAGTGCTTCCTGCTGCTCCC. The lentiviral inserts from the pre-transplant and post-transplant samples were barcoded, pooled and sequenced on a HiSeq 4000 (transplants A and B) or MiniSeq (transplant C) (Illumina). Reads were mapped to the original oligo sequences using Hisat2(*40*). shRNAs with less than 50 reads were discarded. Reads counts were normalized to total read number using DESeq and each shRNA post-transplant to pre-transplant ratio was calculated (enrichment score). Gene level p-values were calculated by Benjamini-Hochberg corrected Mann-Whitney U tests comparing the distribution of enrichment scores of all shRNAs to the tested gene to the distribution of the non-targeting shRNAs. Given the anticipated high level of heterogeneity in these different human islets, we set a significance threshold of 0.1.

### Human islet donor characteristics

Human islets were obtained from the UCSF Human Islet Production Core or the IsletCore at the University of Alberta. See Data File S1 for donor characteristics.

### Human beta cell death

5000-6000 IEQs donor islets were infected at MOI 10 with lentiviruses containing either a non-targeting shRNA expressed in the 3’UTR of nuclear GFP or 5-HT_1F_ shRNA in the 3’ UTR of nuclear mCherry for 5 days. Both were driven by the proximal 360 base pairs of the human insulin promoter. Separate infections were performed to prevent double infection of the beta cells with the two different shRNAs. Islets were pooled and 1000-1500 IEq were transplanted under the kidney capsule of a scd-beige mouse. The graft was removed after 5 days. Grafts were frozen in O.C.T. (Tissue-Tek) and cryosections were prepared at 5 µm thickness. The primary antibodies were rabbit anti-cleaved caspase-3 (Cell Signaling; 9661; 1:100), chicken anti-GFP (Aves Labs, 1:100) and mouse anti-mCherry (Clonetech, 1:200) overnight at 4 degrees.

Secondary antibodies were anti-rabbit Alexa 647 at 1:300, anti-chicken Alexa 488 at 1:500 and anti-mouse Alexa 555 at 1:300. Imaging was performed using a Leica SP5 confocal laser microscope.

### 1-(2-hydroxy-3-(naphthalen-2-yloxy)propyl)-4-(quinolin-3-yl)piperidin-4-ol binding assay

1-(2-hydroxy-3-(naphthalen-2-yloxy)propyl)-4-(quinolin-3-yl)piperidin-4-ol was assayed for binding to serotonin receptors at the NIMH-PSPD using methods previously described (*41*). In primary binding assays, 10 µM of 1-(2-hydroxy-3-(naphthalen-2-yloxy)propyl)-4-(quinolin-3-yl)piperidin-4-ol was tested in quadruplicate. Targets with a minimum of 50% inhibition were subjected to secondary concentration-response assays in triplicate to determine equilibrium binding affinity (K_i_). Both primary and secondary radioligand binding assays were carried out in a final of volume of 125 µl per well in 96-well plates in appropriate binding buffers. Total binding and nonspecific binding were determined in the absence and presence of 10 µM of the appropriate reference compound, respectively. In brief, plates were usually incubated at room temperature and in the dark for 90 min. Reactions were stopped by vacuum filtration onto 0.3% polyethyleneimine (PEI) soaked 96-well filter mats using a 96-well Filtermate harvester, followed by three washes with cold wash buffers. Scintillation cocktail was then melted onto the microwave-dried filters on a hot plate and radioactivity is counted in a MicroBeta counter. Results were analyzed with the built-in one-site binding competition binding function in GraphPad Prism V10.

### Glosensor cAMP in 293T cells

1-(2-hydroxy-3-(naphthalen-2-yloxy)propyl)-4-(quinolin-3-yl)piperidin-4-ol was assayed at the NIMH-PDSP using techniques previously described (*42*) with modifications. Briefly, HEK293T cells were transiently transfected with the Glosensor-cAMP plasmid (Promega) with or without 5-HT_1F_ or 5-HT_1A_ cDNA. One day post transfection, cells were plated in 384-well white plates coated with 0.01% poly-L-lysine (Sigma Aldrich) at a density of 10K cells in 40 µl / well in DMEM supplemented with 1% dialyzed FBS, and used for assays after recovery for 5-6 hours. Media was removed and cells received 20 µl/well drug dilutions at 1.5x of final in assay buffer (1x HBSS, 20 mM HEPES, pH 7.40, supplemented with 0.1% BSA and GloSensor reagent). The plate was counted on a luminescence counter for G_s_ agonist activity at 20 min after drug addition, followed by addition of 10 µl isoproterenol (final at 100 nM) for G_i_ agonist activity at 20 min later. For antagonist activity assays, cells received 10 µl of 3x antagonist solutions for 10 min, 10 µl of 5-HT for another 10 min, followed by 10 µl of isoproterenol (final of 100 nM). The plate was read at 20 min after isoproterenol. Luminescence counts were normalized and analyzed using the built-in 4-parameter logistic function in the GraphPad Prism V10.

### Human beta cell proliferation

Human islets were dissociated and plated as above in 96 well plates (Perkin Elmer, 6055302) and were infected with MOI 1 of lentivirus expressing shRNA for 48 hours before drug treatment. For the 1-(2-hydroxy-3-(naphthalen-2-yloxy)propyl)-4-(quinolin-3-yl)piperidin-4-ol experiments, islets were immediately treated with DMSO, Exendin-4 5 nM, harmine 10 μM, or exendin-4 and harmine for 48-72 hours. Cells were then fixed with 4% paraformaldehyde for 15 minutes at 37 ℃. Cells were stained with anti-KI67(BD Pharmigen, 1:200), chicken anti-GFP (Aves Labs, 1:500), and insulin (Agilent, 1:500) overnight at 4 ℃. The secondary antibodies were, anti-chicken Alexa 488, anti-mouse Alexa 555 at 1:300, and anti-guinea pig 647 1:500 (Invitrogen). Plates were imaged and analyzed using the Opera Phenix Plus High Throughput Confocal and Harmony High Content Imaging and Analysis software.

### GSIS in Human Islets

Human islets were rested overnight in islet media as above Forty to sixty islets were hand-picked and equilibrated for two hours at 37 ℃ in 1 mL Kreb-Ringer Bicarbonate HEPES Buffer pH 7.4 (KRBH; 137 mM NaCl, 4.7 mM KCl, 1.2 mM KH2PO4, 1.2 mM MgSO4, 2.5 mM CaCl2, 25 mM NaHCO3, 20 mM HEPES, 2.8 mM glucose, and 0.25% BSA). Buffer was then removed and islets were incubated at 37 ℃ with 1mL KRBH with 2.8 mM glucose and and DMSO or 150 nM 1-(2-hydroxy-3-(naphthalen-2-yloxy)propyl)-4-(quinolin-3-yl)piperidin-4-ol or water and 100 nM LY344864. After 1 hour, islets were changed to 1mL KRBH 11 mM glucose and 0.25% BSA with or without drug at 37 ℃ for another hour. The islets were then lysed in 1 mL RIPA buffer to determine the total islet insulin content. Insulin and glucagon were measured by ELISA (Mercodia, 10-1113-01, Mercodia, 10-1271-01).

### Human Islet Adenovirus Infection

An insulin nanoluciferase fusion (*35*) and was placed in a shuttle vector encoding the rat insulin promoter and rabbit beta-globin intron (a gift from Matt Merrins) (*43*) between EcoRI and XhoI with a T2A and BFP cloned by HiFi (NEB). Sequence was validated by Sanger sequencing. LR recombination was then performed with the Gateway LR clonase Enzyme mix (Invitrogen, 11791019). Adenoviral vectors expressing control non-targeting shRNA and shRNA targeting 5-HT_1F_ were produced by cloning the U6 shRNA (A) into pAdtrack-CMV-mCherry (*44*). The loop was changed to TTCAAGAGA. Adenovirus production was performed as previously described (*45*). Human islets were infected with MOI 50 of the nanoluciferase t2a-bfp and MOI 200 of control non-targeting shRNA and shRNA targeting 5-HT_1F_ adenovirus. 72 hours later islets were picked and GSIS was performed as above.

### Human Islet Adenovirus Imaging

Human islets were infected with adenovirus as described above and were imaged in 96 well imaging plates (Perkin Elmer, 6055302) using Opera Phenix Plus High Throughput Confocal and images were analyzed using the Harmony High Content Imaging and Analysis software.

### Nanoluciferase

Nanoluciferase activity was measured from 100 μl of supernatants and lysates (Promega, N1110) using a Perkin Elmer Envision.

### GloSensor cAMP measurement in human beta cells

GloSensor, cAMP-22p was cloned into pEN-RIP1-BGH-pA vector with T2a-h2b-mCherry between the EcoRI and XhoI by HiFi assembly (NEB, E2621). The correct sequence was validated by Sanger sequencing and virus was produced as above. Human islets were infected with MOI 200. 48 hours later, islets were picked and incubated for 2 hours in HBSS+10% FBS+2.8 mM glucose+2% of the GloSensor reagent in the machine taking readings every 15 min. 1-(2-hydroxy-3-(naphthalen-2-yloxy)propyl)-4-(quinolin-3-yl)piperidin-4-ol at 300 nM or LY344864 at 100 nM were added at time −5 min. 1 μM of forskolin was added at time 0 and luminescence was measured at 1 minute intervals using the Perkin Elmer Envision.

### Animals

All mice were group or individually housed with a 12 hr light/dark cycle at 70 °F and 45% humidity. Mice had free access to food and water (PicoLab Mouse Diet 20). Experiments were conducted according to the Guide for the Care and Use of Laboratory Animals and with UCSF’s Institutional Animal Care Use Committee (IACUC) approval. Male mice aged 8-15 weeks were used for all experiments.

### Drug Treatment in Untransplanted Mice

NOD.Cg-Prkdc^scid^Il2rg^tm1Wjl^/Sz (NSG) mice were IP injected with 20 mg/kg of 1-(2-hydroxy-3-(naphthalen-2-yloxy)propyl)-4-(quinolin-3-yl)piperidin-4-ol dissolved in 5% DMSO + 1.7% lactic acid pH 4. For testing potential effects on bodyweight and glucose or insulin tolerance in unengrafted mice, mice were injected every 12 hours (2x daily) with vehicle or 20 mg/kg of 1-(2-hydroxy-3-(naphthalen-2-yloxy)propyl)-4-(quinolin-3-yl)piperidin-4-ol for 7 days before tolerance tests were performed.

### Blood collection

Distal tail vein blood was collected into heparin coated tubes (Sarstedt, 16.443.100) and centrifuged at 2,000 x g 4 °C for 10 min. Plasma was stored at −80°C until downstream use.

### Pharmacokinetics

10 μL of plasma was extracted in 20 μL of 1:1 acetonitrile:0.1% formic acid. Samples were vortexed and centrifuged to pellet protein and supernatants were collected. Targeted measurements of plasma 5-HT_1F_ antagonist levels were quantified using an Agilent 6470 Triple Quadrupole (QQQ) LC/MS. Separation of metabolites was conducted on a Kinetix 2.6 μm Polar C18 100 Å LC column (Phenomenex 00B-4759-AN) using reverse phase chromatography. Mobile phases were 0.1% formic acid in water (A) and acetonitrile (B). The LC gradient started with 5% mobile phase B with a flow rate of 0.25 ml/min from 0–0.4 min. The gradient was then increased linearly to 50% A/50% B at a flow rate of 0.25 ml/min from 0.4–1.7 minutes. From 1.7 to 3.2 minutes the gradient was maintained at 50% A/50% B. Mass spectrometry analysis was performed using electrospray ionization (ESI) in positive mode. Additional parameters were set as follows: gas temperature, 250 °C with a gas flow of 12 l/min; nebulizer pressure, 25 psi; sheath gas temperature, 300 °C with the sheath gas flow of 12 l/min; capillary voltage, 3500 V. Multiple reaction monitoring (MRM) was performed for 5-HT_1F_ antagonist detection using the following parameters: precursor ion, 429.2; product ion, 70.1; dwell, 50; fragmentor, 167; collision energy, 38; cell accelerator voltage, 4; polarity, positive. Plasma levels were quantified against an 5-HT_1F_ antagonist standard curve using Agilent MassHunter Quantitative Analysis software.

### Glucose Tolerance and Insulin Tolerance Tests

Mice were individually housed and fasted with free access to water. For glucose tolerance tests: 6 hours fasted with 2 g/kg glucose. For insulin tolerance tests: 5 hours fasted with 0.1 mU/kg of insulin. Glucose levels were taken from distal tail samples using the Freestyle Lite glucometer (Abbott). Plasma samples were collected as described above and 10 μl of plasma were used for the Insulin Ultrasensitive HTRF kit (62IN2PEG).

### Human Islet transplant into NSG mice

NSG mice were made diabetic with 170 mg/kg streptocozin (Sigma, S0130) in lactate ringer solution. Mice with blood glucose levels >350 mg/dL for over 48 hours were used for the transplant studies. Human islets were cultured at a concentration of 1,000 Islet Equivalents (IEQs) per 5 ml of media (CMRL 1066 Supplemented, CIT modification (Corning, #98-304-CV) with 10 ml of Human Serum Albumin 25%, 5000 U Heparin (stock at 5,000 U/ml, Fresenius Kabi, #4710), 10 mg Ciprofloxacin (stock at 10 mg/ml, Claris), 1 mg DNase I (Roche, 11284932001), and Antibiotic-antimycotic (Gibco, 15240062). Islets were rested 16-48 hours after isolation prior to transplant. 6 hours before transplant, islets were treated with vehicle or 200 nM 1-(2-hydroxy-3-(naphthalen-2-yloxy)propyl)-4-(quinolin-3-yl)piperidin-4-ol. Approximately 2,000 IEQs were transplanted under the mouse kidney capsule. For the transplant experiments, mice were injected intraperitoneally with vehicle or 20 mg/kg of 1-(2-hydroxy-3-(naphthalen-2-yloxy)propyl)-4-(quinolin-3-yl)piperidin-4-ol ∼6 hours before transplant and then every 12 hours for 7 days before tolerance tests were performed. For the stable engrafted mouse experiments (4-7 weeks post-transplant), the mice were IP injected as described above a single time 2 hours before the tolerance tests.

### Human islets

Human islets for research were provided by the Alberta Diabetes Institute Islet Core at the University of Alberta in Edmonton (http://www.bcell.org/adi-isletcore.html) with the assistance of the Human Organ Procurement and Exchange (HOPE) program, Trillium Gift of Life Network (TGLN), and other Canadian organ procurement organizations. Islet isolation was approved by the Human Research Ethics Board at the University of Alberta (Pro00013094). All donors’ families gave informed consent for the use of pancreatic tissue in research. Human islets for research were also provided by the UCSF Islet Production Core.

### Chemicals

Chemicals used were as follows: 1-(2-hydroxy-3-(naphthalen-2-yloxy)propyl)-4-(quinolin-3-yl)piperidin-4-ol synthesized by Enamine CAS# 256372-99-3, LY344864 (Tocris, 2451), streptozotocin (Sigma, S0130), exendin-4 (Medchemexpress, 50-202-8996), harmine (Sigma, 286044), isoproterenol (Sigma, 16504).

### shRNAs

These sequences were cloned into the mir-30 context of pSC2. Non targeting shRNA sequence (including loop): TAAGACTCGAATTGTAGTGTCATAGTGAAGCCACAGATGTATGACACTACAATTCGA GTCTTT 5-HT_1F_ targeting shRNA sequence (including loop) (sh-A): TTAGAAGATATACGAAATAATATAGTGAAGCCACAGATGTATATTATTTCGTATATCTTCTAT 5-HT_1F_ targeting shRNA sequence (including loop) (sh-B): TTTGGTAGAAATGAACAGGAAATAGTGAAGCCACAGATGTATTTCCTGTTCATTTCTACCAAT

### Droplet PCR

RNA was isolated from human islets using Trizol reagent (Thermo), treated with DNase I, and reverse transcribed with Superscript IV (Thermo). Quantitation was performed using the BioRad QX100 system. TBP was used for control expression (Applied Biosystems, 4333769T). 5-HT_1F_ F=AGAGTTCCCATTTTATACAGGGCA, 5-HT_1F_ -R=CCTCTGCACCATCATAACTGT, 5-HT_1F_ probe=AGCAAACTAAAATCTGGC

### Statistical analysis

Tests are listed in figure legends and were calculated with GraphPad (Prism), R, or custom python script. For Fig. 6B AUC and Fig. 6C, a Mann-Whitney test was used because the data were not normally distributed.

### Study Approval

These studies were approved by the UCSF IACUC (AN170193). Human islet donors were anonymized and therefore not considered to be human subjects research by the UCSF IRB (18–26481).

## Supporting information

S1 Donor Information

S2 p-values

## List of Supplementary Materials

Fig. S1

Fig. S2

Fig. S3

Fig. S4

Fig. S5

Fig. S6

Fig. S7

Fig. S8

Data file S1: Donor information

Data file S2: p-values for screen References (1–45)

## Acknowledgments

We would like to thank Michael S. German, Michael T. McManus, Gerold M. Grodsky and Gregory Szot for helpful discussions and Vi Dang, Vinh Nguyen, and Gregory Szot from the UCSF Islet Production Core. Ki determinations, receptor binding profiles, and functional data was generously provided by the National Institute of Mental Health’s Psychoactive Drug Screening Program, Contract # HHSN-271-2018-00023-C (NIMH PDSP). The NIMH PDSP is Directed by Bryan L. Roth MD, PhD at the University of North Carolina at Chapel Hill and Project Officer Jamie Driscoll at NIMH, Bethesda MD, USA. We would also like to thank the Parnassus Center For Advanced Technologies which supplied the Opera Phenix Plus High Throughput Confocal and ddPCR equipment used. Thank you to the UCSF Islet Production Core and the University of Alberta Diabetes Institute Islet Core for supplying the islets used in this study. Finally, we thank the generous individuals who donated their pancreatic islets so that future patients might benefit.

## Funding

JDRF (5-CDA-2014-199-A-N) (GMK)

JDRF (2-SRA-2021-1036-M-N) (GMK)

Nora Treadwell Foundation Grant (GMK)

R01 DK107650 (GMK)

R01 DK118337 (GMK)

K08 DK087945 (GMK)

P30 DK063720 (a Pilot and Feasibility Grant to GMK and core laboratories)

The UCSF Islet Production Core is supported by Diabetes Research Center (DRC) grant NIH P30 DK063720

Kraft Post-doctoral Fellowship (RAL)

## Author contributions

Conceptualization: GMK

Methodology: RAL, DGC, VN, YZ, KS, NY, RS, JA, XPH, BR, and GMK

Investigation: RAL, DGC, VN, YZ, KS, NY, RS, XPH, and GMK

Visualization: RAL, DGC, GMK

Funding acquisition: GMK

Writing: RAL, DGC, GMK

## Data and materials availability

All data are available in the main text or supplementary materials.

## Supplementary Materials

**Fig. S1.**
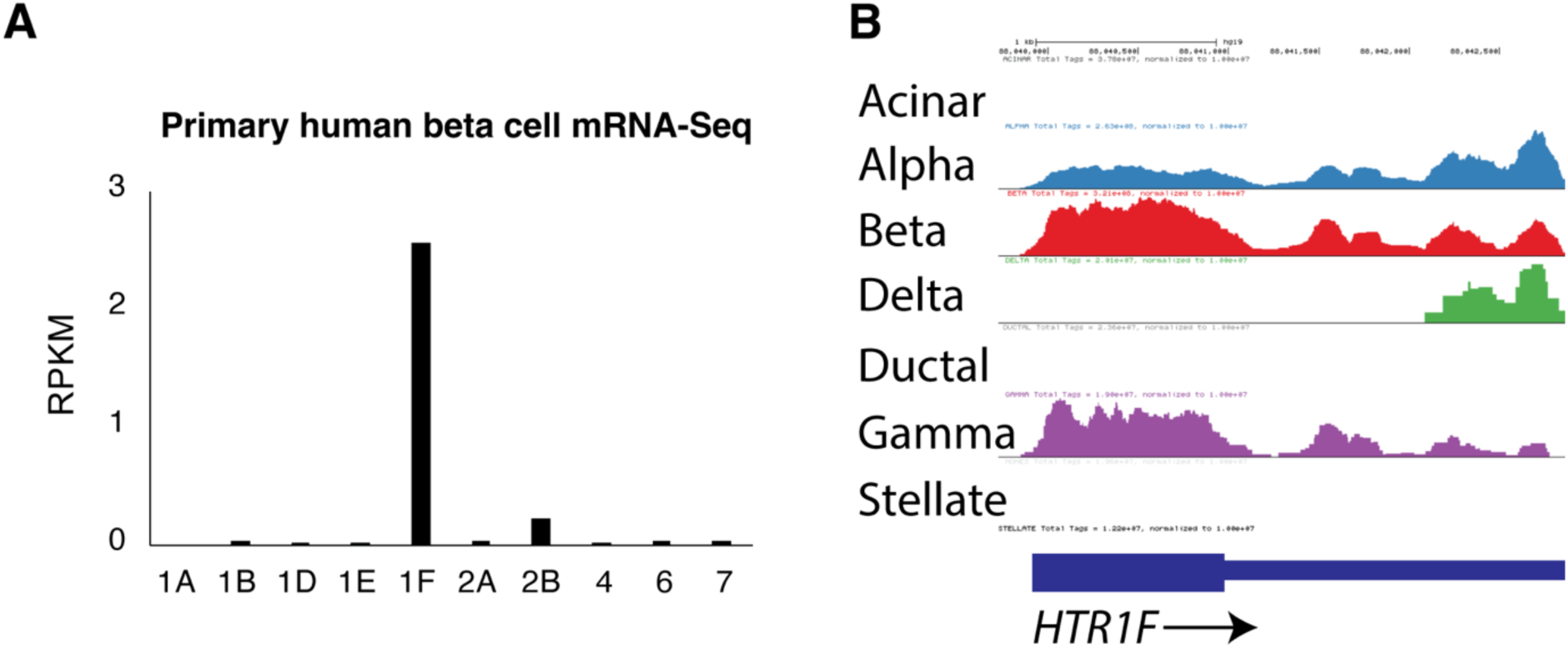
*5-HT_1F_* is the most highly expressed serotonin receptor in adult primary human beta cells. **(A)** Expression of the indicated serotonin receptor gene from conventional mRNA-seq data from sorted primary human beta cells. Expression is reported in reads per kilobase per million (RPKM). Data replotted from(*14*). **(B)** Expression of 5-HT_1F_ in alpha, in human beta, delta, and gamma cells by aggregated single cell mRNA-seq. Replotted from (*28*).

**Fig. S2.**
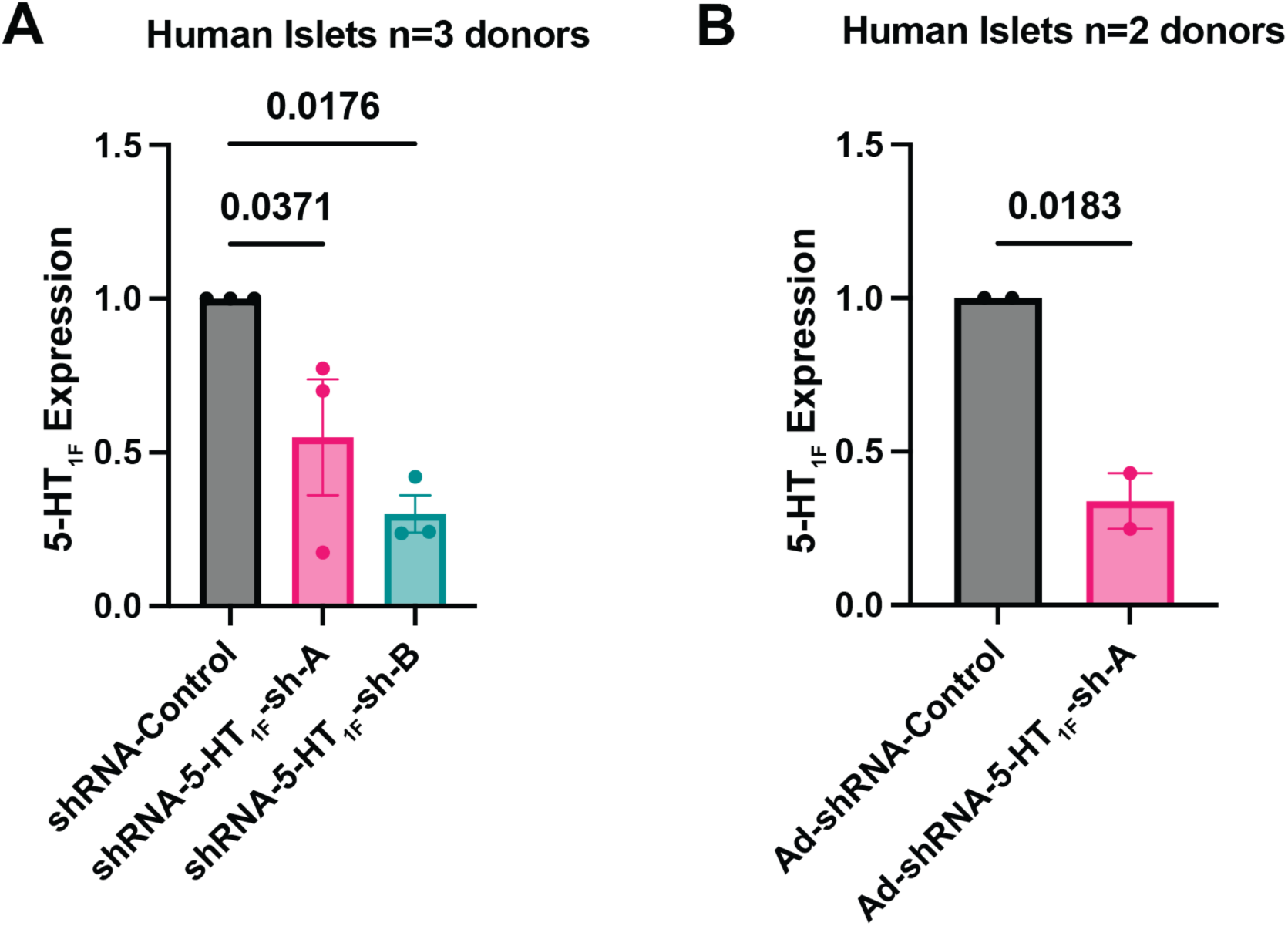
*5-HT_1F_* knockdown in human islets. **(A)** 5-HT_1F_ mRNA normalized to TBP in human islets infected with lentivirus expressing shRNA a control, or two separate shRNAs targeting 5-HT_1F_: 5-HT_1F_ -sh-A or 5-HT_1F_ -sh-B from 3 independent donors with 3-4 replicates from each donor. P-values from a One-way ANOVA with Benjamini-Hochberg correction. (**B)** 5-HT_1F_ mRNA normalized to TBP in human islets infected with adenovirus expressing shRNA for control or 5-HT_1F_-sh-A from two independent donors. P-values from non-paired Student’s t-test. Error bars show standard error.

**Fig. S3.**
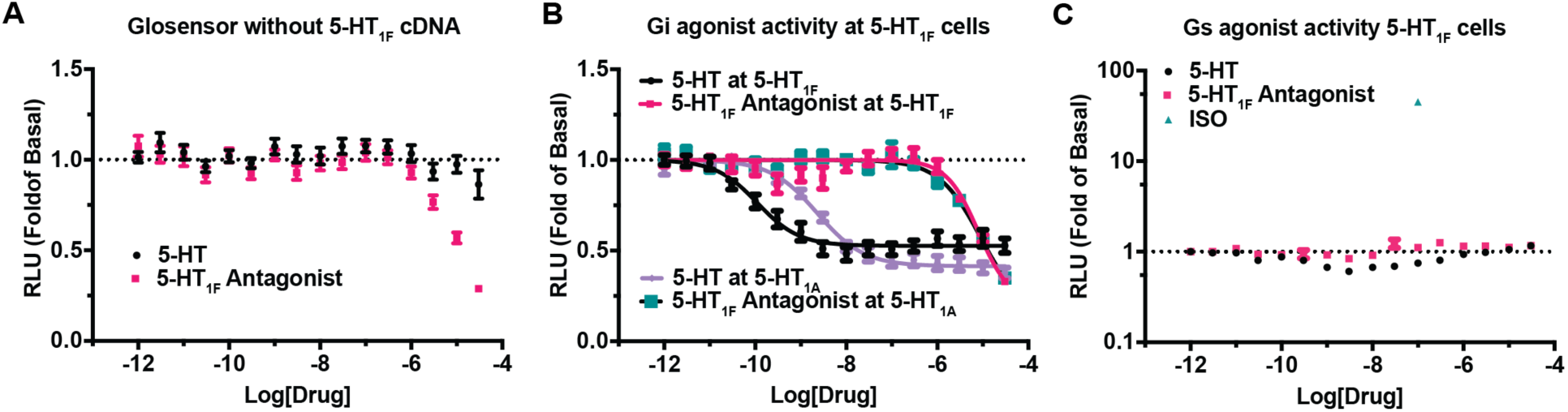
1-(2-hydroxy-3-(naphthalen-2-yloxy)propyl)-4-(quinolin-3-yl)piperidin-4-ol is an 5-HT_1F_ antagonist. **(A)** 293T cells transfected with the cAMP GloSensor plasmid (Promega) alone and treated with the indicated drug at the indicated concentration and then with 0.1μM isoproterenol prior to luminescence measurement, Error bars show standard error from n=4 independent assays, each in quadruplicate. **(B)** 293T cells were transiently transfected with the GloSensor plasmid (Promega) and the cDNA for 5-HT_1A_ or 5-HT_1F_ and were treated the indicated drugs at the indicated concentrations. Prior to luminescence measurement, the cells were treated with 0.1 μM isoproterenol. Error bars show standard error from n=4 independent assays, each in quadruplicate. **(C)** 293T cells transfected and treated as above without isoproterenol. Error bars show standard error from n=4 independent assays, each in quadruplicate.

**Fig. S4.**
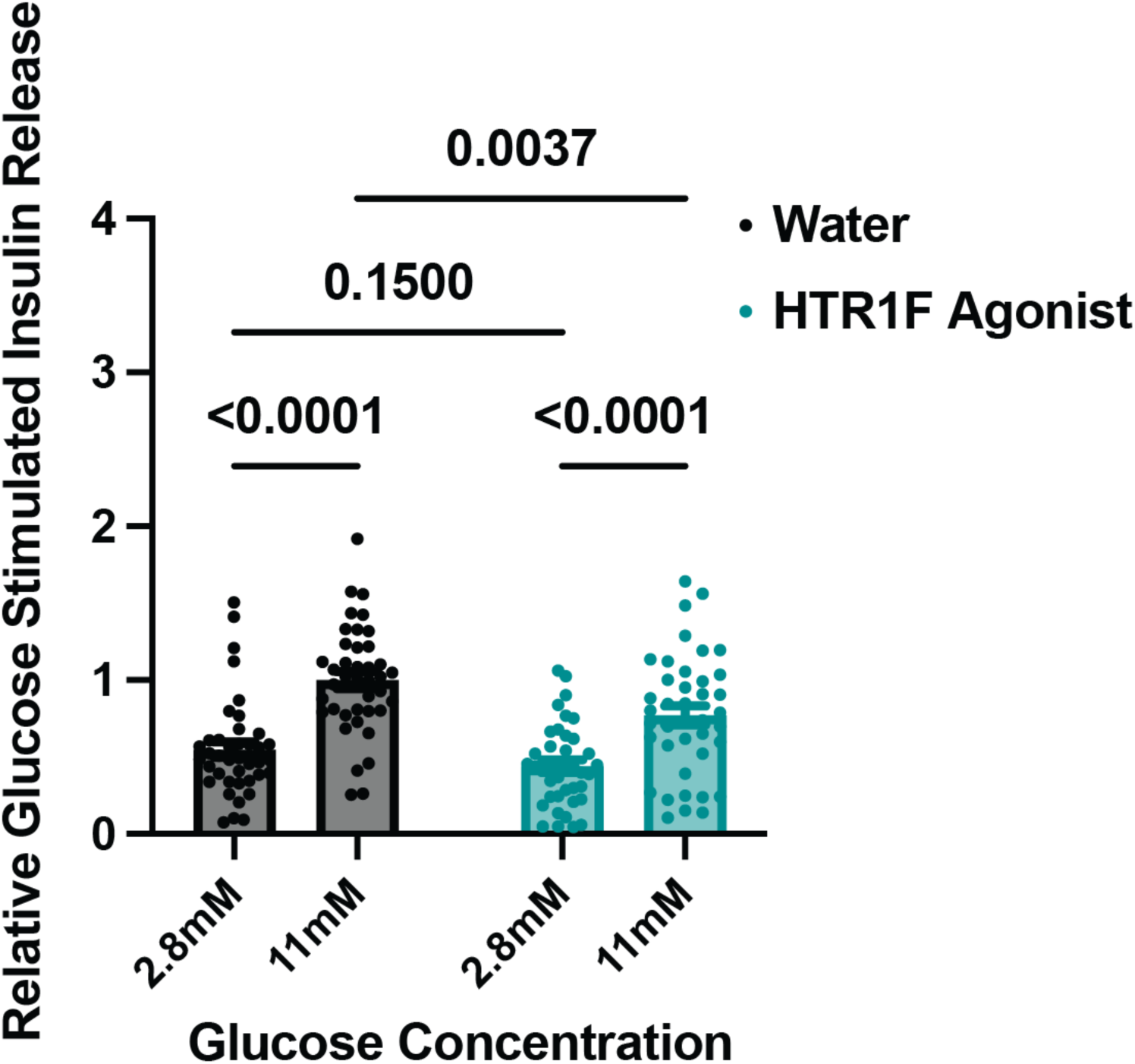
5-HT_1F_ agonist inhibits glucose stimulated insulin secretion from human islets. Human islets were pre-incubated with vehicle (water) or 150 nM 5-HT_1F_ agonist (LY344864) for 1 hour and during each hour of stimulation at 2.8 mM glucose and 11 mM glucose. n=8 independent donors, 5 replicates for each donor. Two-way ANOVA: drug effect p=0.0017, glucose effect p<0.0001. Interaction p=0.25. Post-hoc tests with Benjamini-Hochberg correction. Error bars show standard error.

**Fig. S5.**
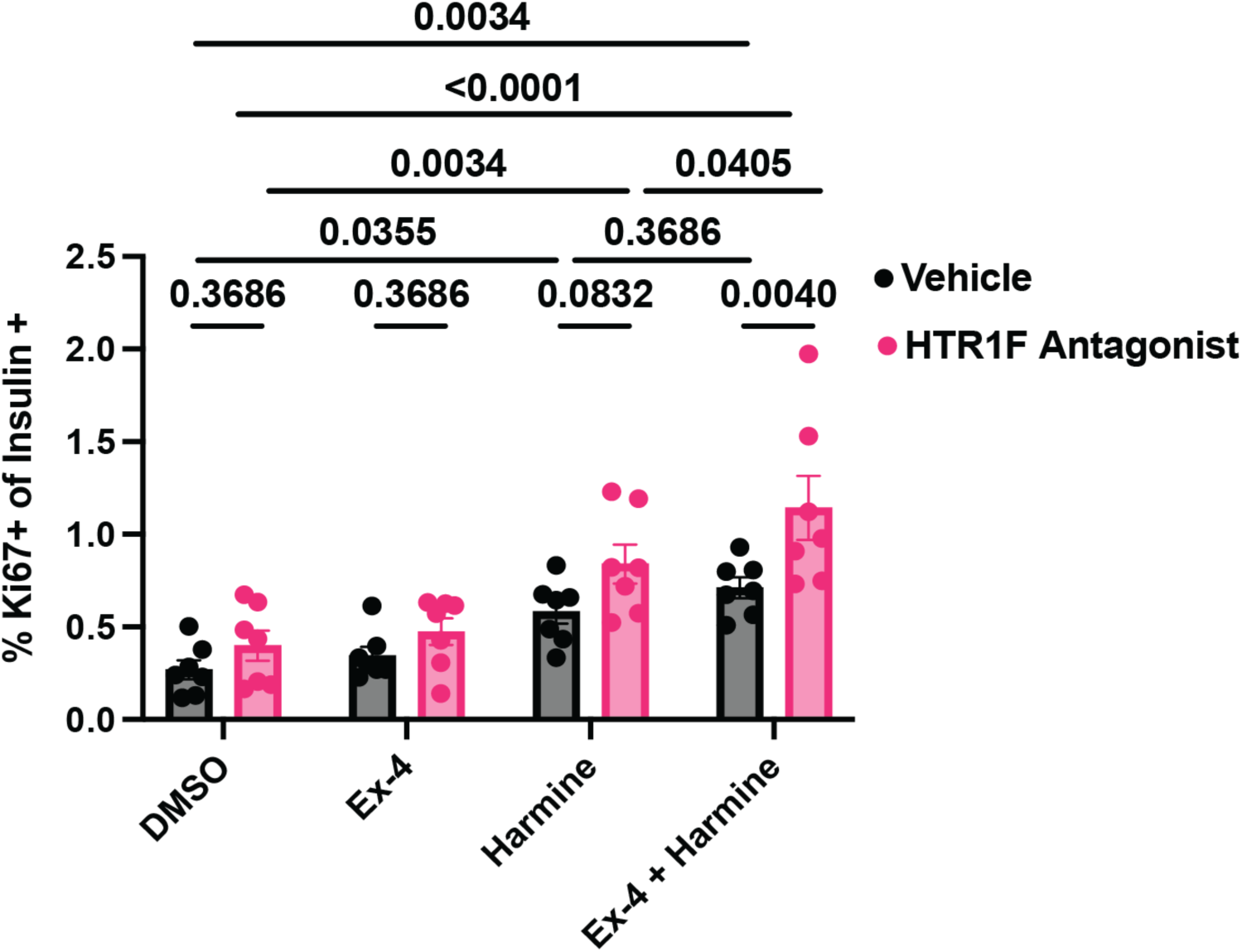
5-HT_1F_ antagonist increases human beta cell proliferation but only in the presence of harmine and exendin-4. Dissociated human islets were pretreated with 200nM 1-(2-hydroxy-3-(naphthalen-2-yloxy)propyl)-4-(quinolin-3-yl)piperidin-4-ol or vehicle for 24 hours and then treated with the indicated drugs for an additional 48-72 hours. n=7 donors. 2-way ANOVA: p=0.0005 for 1-(2-hydroxy-3-(naphthalen-2-yloxy)propyl)-4-(quinolin-3-yl)piperidin-4-ol effect, p<0.0001 for drug effect, p=0.3002 for interaction. Post-hoc testing with Benjamini-Hochberg correction. Only p-values <0.05 are shown for clarity (out of 28 tests). Error bars show standard error.

**Fig. S6.**
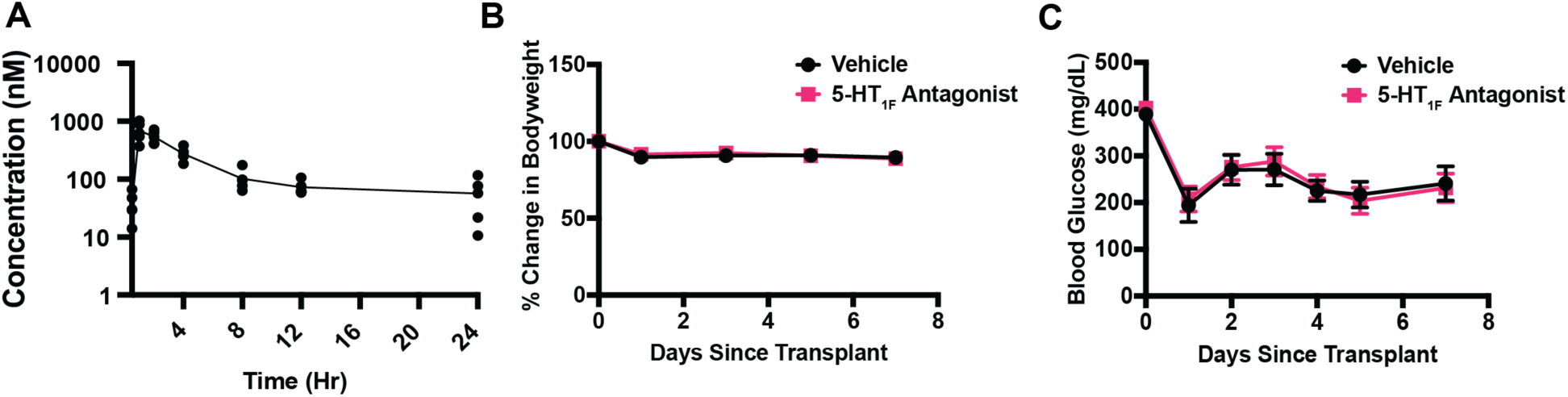
5-HT_1F_ antagonist in NSG mice. **(A)** Plasma levels of 1-(2-hydroxy-3-(naphthalen-2-yloxy)propyl)-4-(quinolin-3-yl)piperidin-4-ol in mice after a single 20 mg/kg IP injection. n=5 mice. **(B)** Bodyweights of diabetic NSG mice who received human islet grafts treated with vehicle n=16 or 5-HT_1F_ antagonist n=15 from 3 independent donors. Two-way ANOVA: drug effect p=0.9631, time p<0.0001, interaction p=0.9414. **(C)** Random blood glucose levels. of diabetic NSG mice who received human islet grafts treated with vehicle n=16 or 5-HT_1F_ antagonist n=15 from 3 independent donors. Two-way ANOVA: drug effect p=0.7722, time p<0.0001, interaction p=0.9252. Error bars show standard error.

**Fig. S7.**
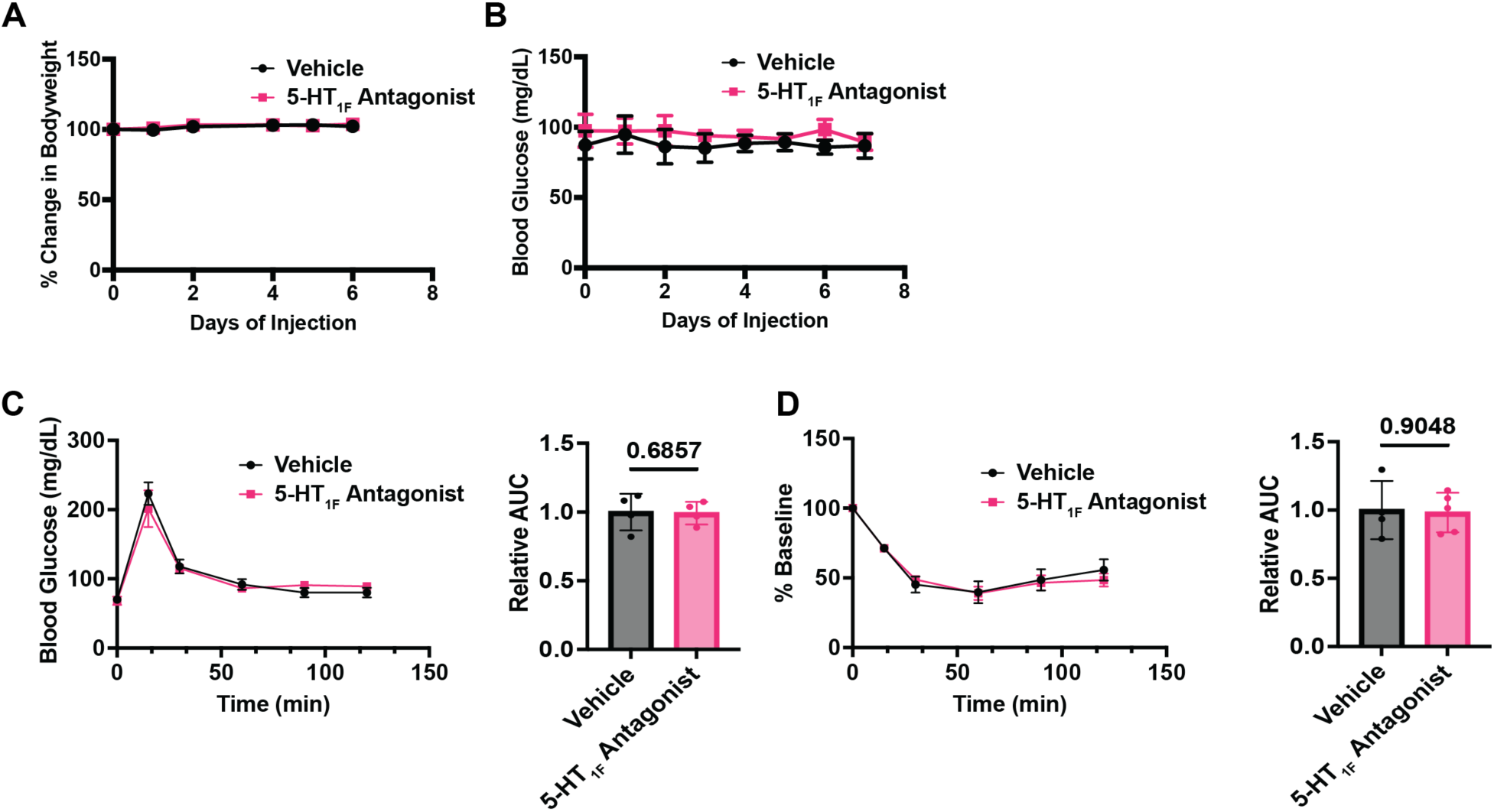
5-HT_1F_ antagonist in NSG mice without human islet transplant. **(A)** Bodyweight of non-transplanted NSG mice during a 7-day course of treatment with vehicle or HT_1F_ antagonist (20 mg/kg injected twice a day). Vehicle n=4, 5-HT_1F_ antagonist n=5. Two-way ANOVA: drug effect p=0.9156, time p=0.0109, interaction p=0.9156. **(B)** As in B, but non-fasting blood glucose is shown. Vehicle n=4, 5-HT_1F_ antagonist n=5. Two-way ANOVA: drug effect p=0.3075, time p=0.9879, interaction p=0.9931. **(C)** IPGTT on the mice from (**A**) at day 7. Vehicle n=4, 5-HT_1F_ antagonist n=4. Two-way ANOVA: drug effect p=0.8224, time p<0.0001, interaction p=0.6099. Relative area under the curve was calculated, p=0.6857 Mann-Whitney test. (**D**) Intraperitoneal insulin tolerance test on the mice from (B) at day 8. Vehicle n=4, 5-HT_1F_ antagonist n=5. Two-way ANOVA: drug effect p=0.8367, time p<0.0001, interaction p=0.7264. Relative area under the curve was calculated, p=0.9048 Mann-Whitney test. Error bars show standard error.

**Fig. S8.**
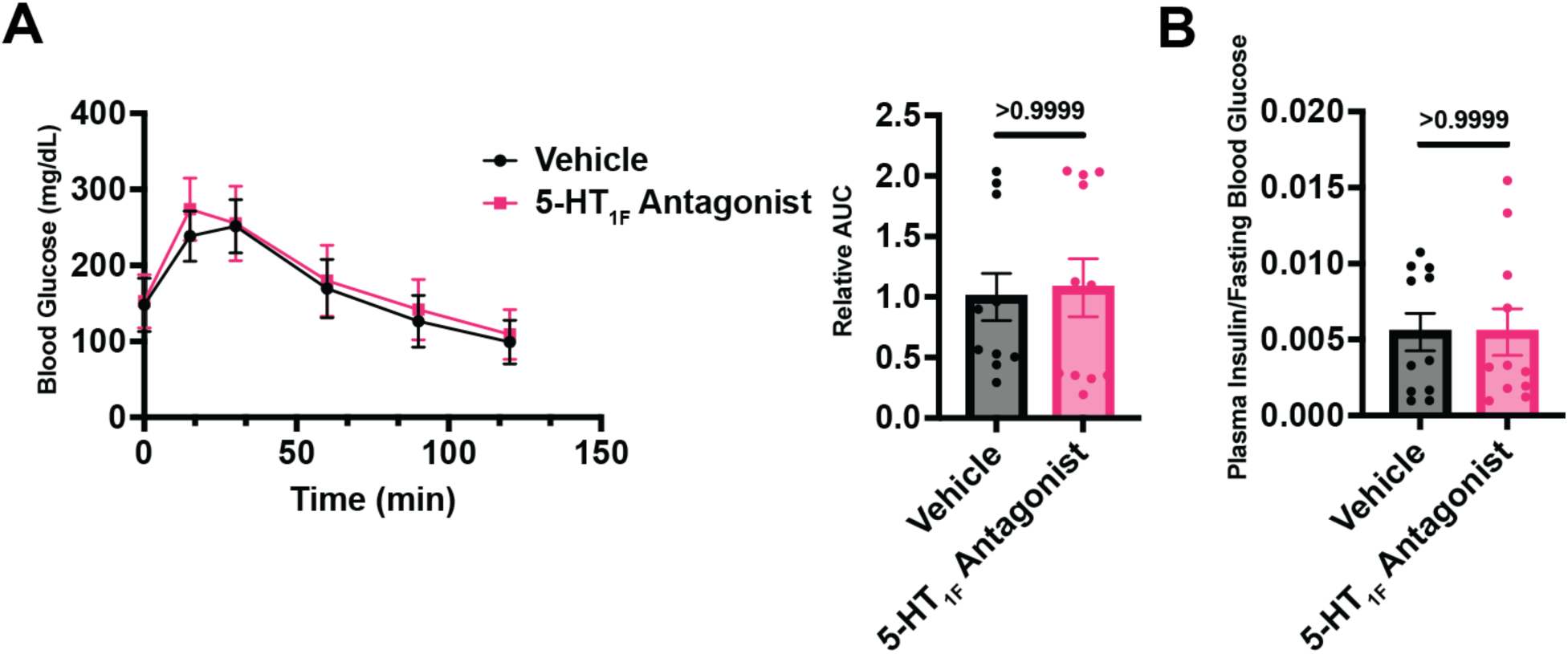
No effect of 5-HT_1F_ antagonist on glycemia or insulin in mice with stable human islet engraftment. **(A)** IPGTT in diabetic NSG mice with human islets stably engrafted IP injected with vehicle or 1-(2-hydroxy-3-(naphthalen-2-yloxy)propyl)-4-(quinolin-3-yl)piperidin-4-ol 2 hours before the GTT. Two-way ANOVA: drug effect p=0.7998, time p<0.0001, interaction p=0.7590. Relative area under the curve was calculated, p>0.9999 Mann-Whitney test. **(B)** Ratio of fasting plasma insulin/fasting blood glucose. Vehicle n=11, 5-HT_1F_ antagonist n=11 from two independent donors, p>0.9999 Mann-Whitney test. Error bars show standard error.

**Data File S1:**
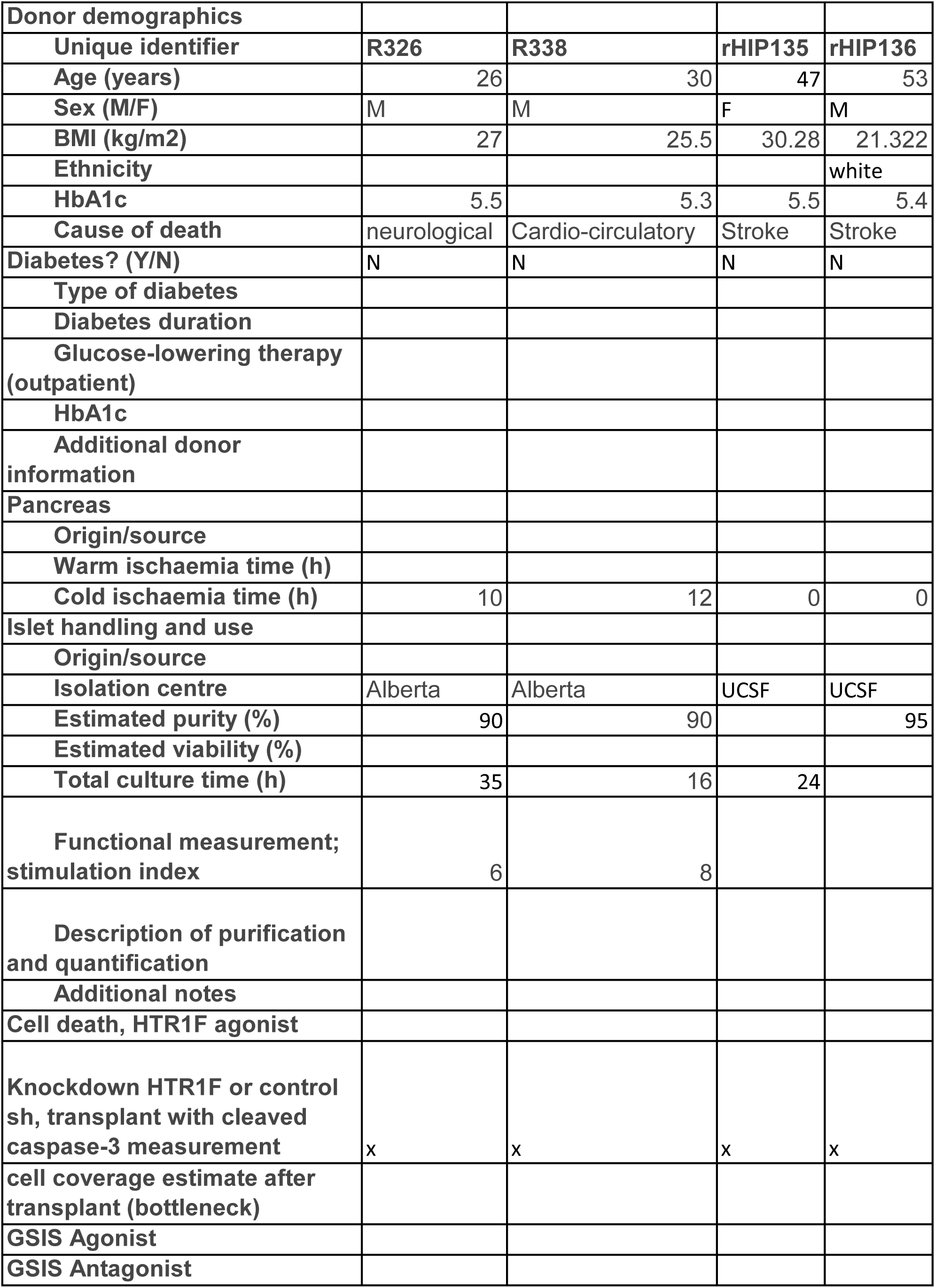
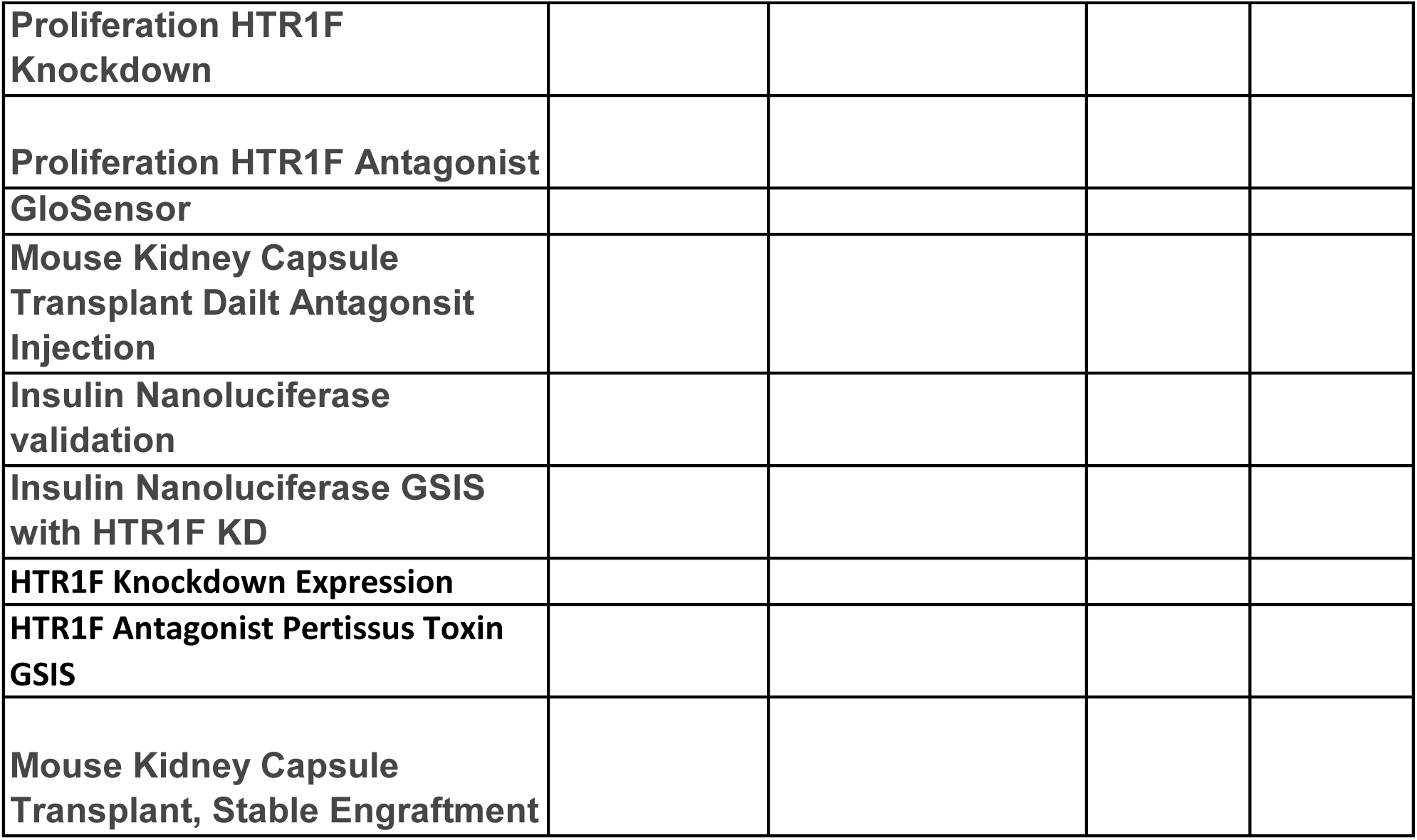
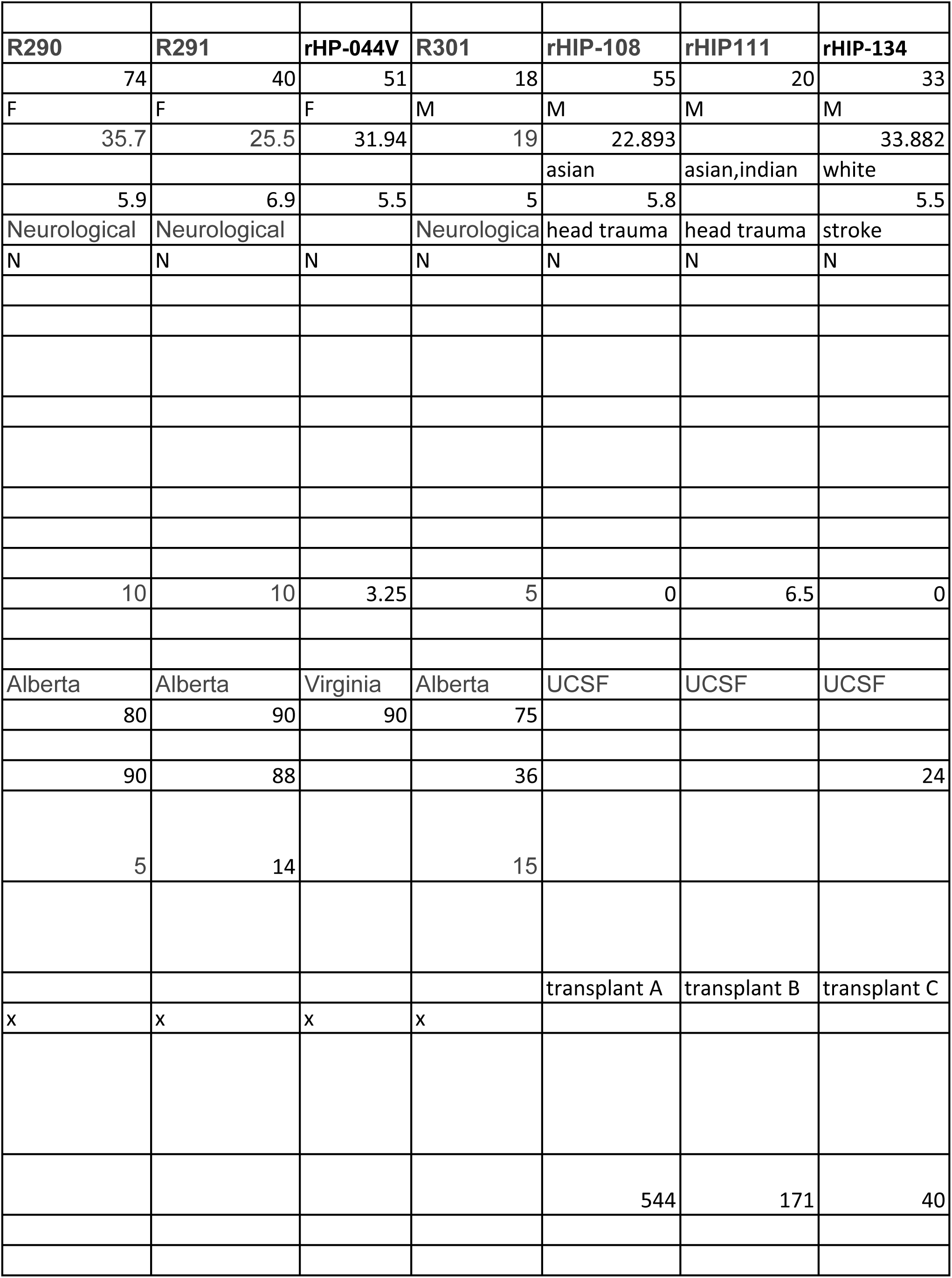
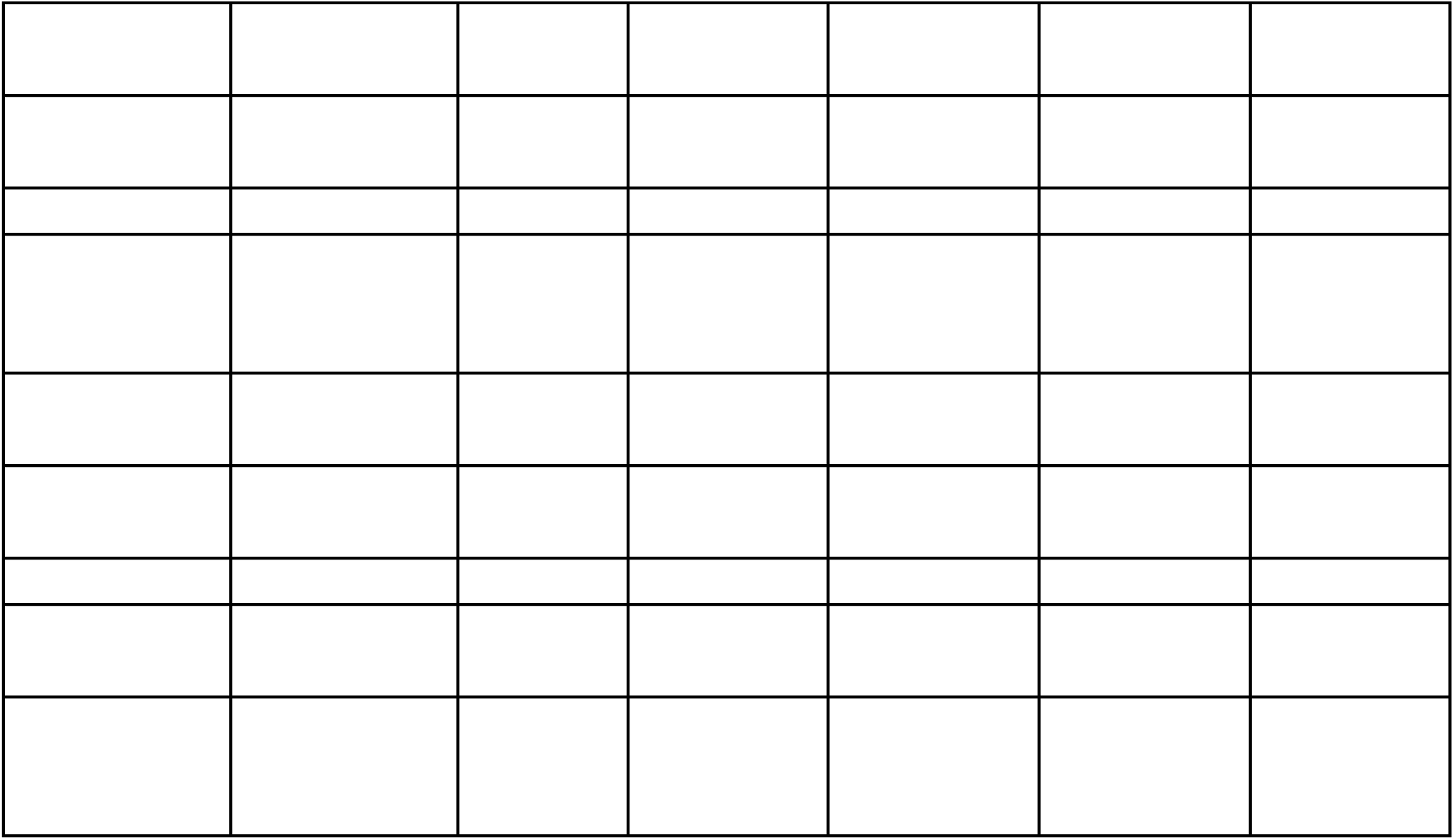
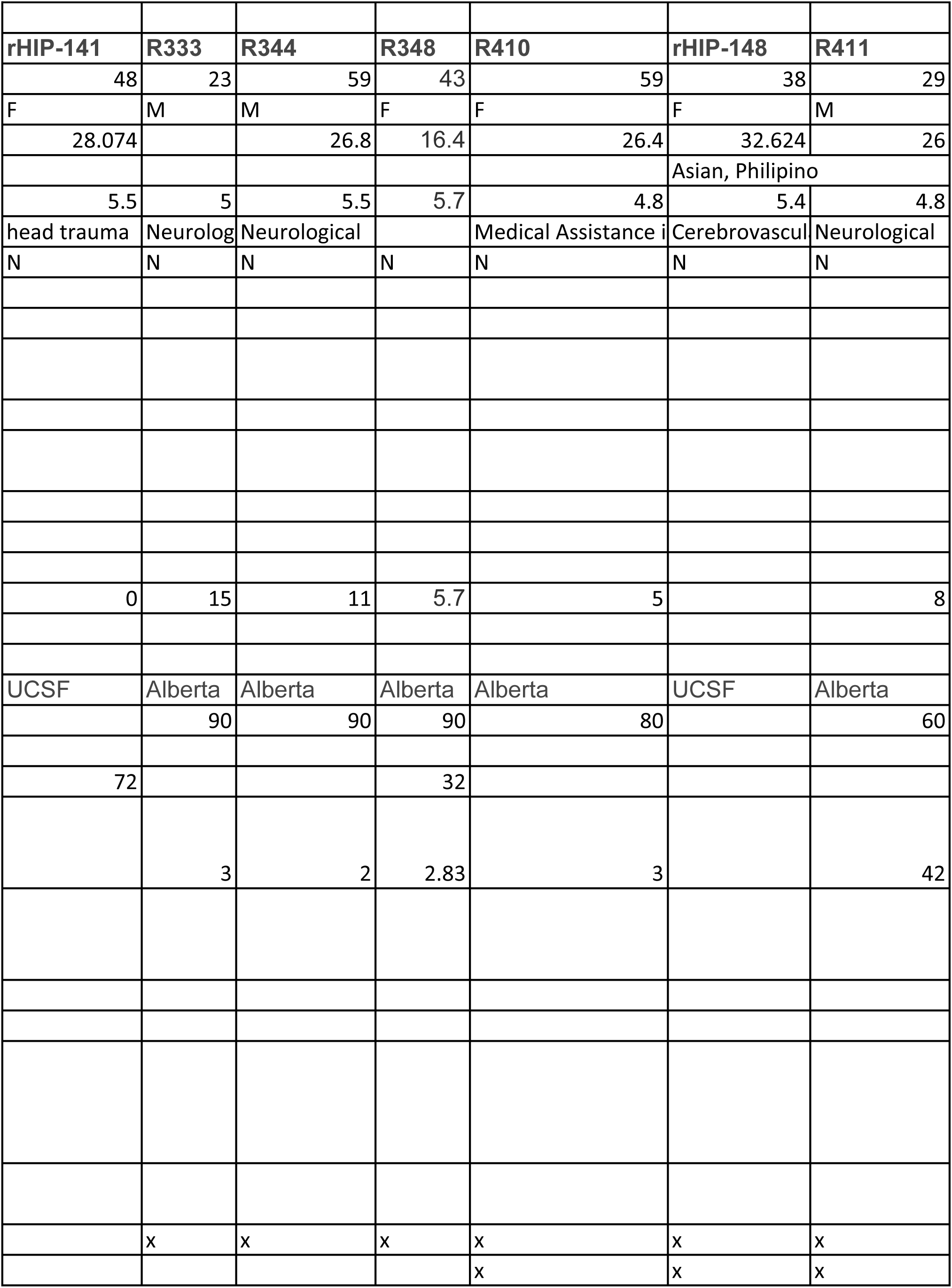
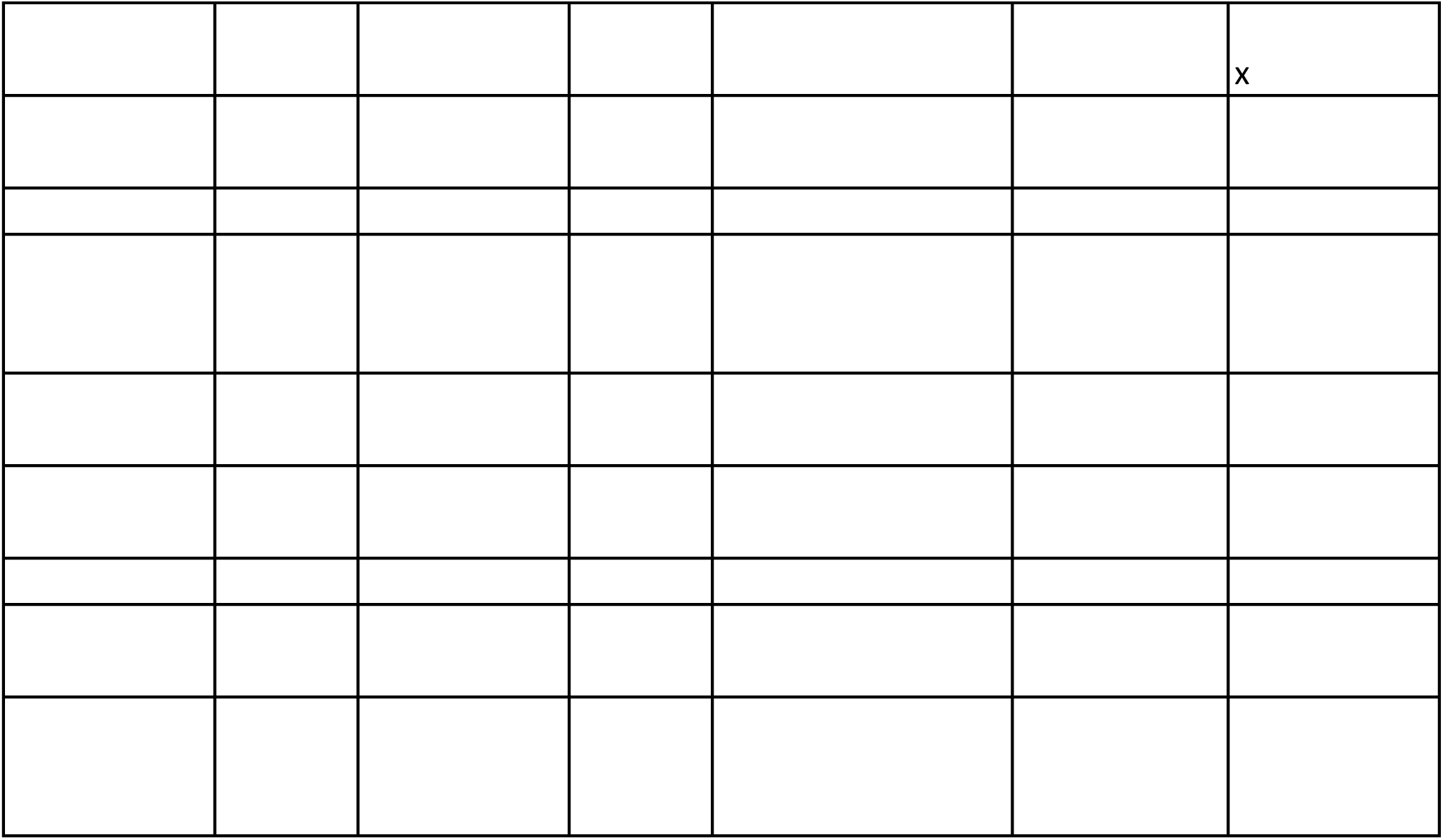
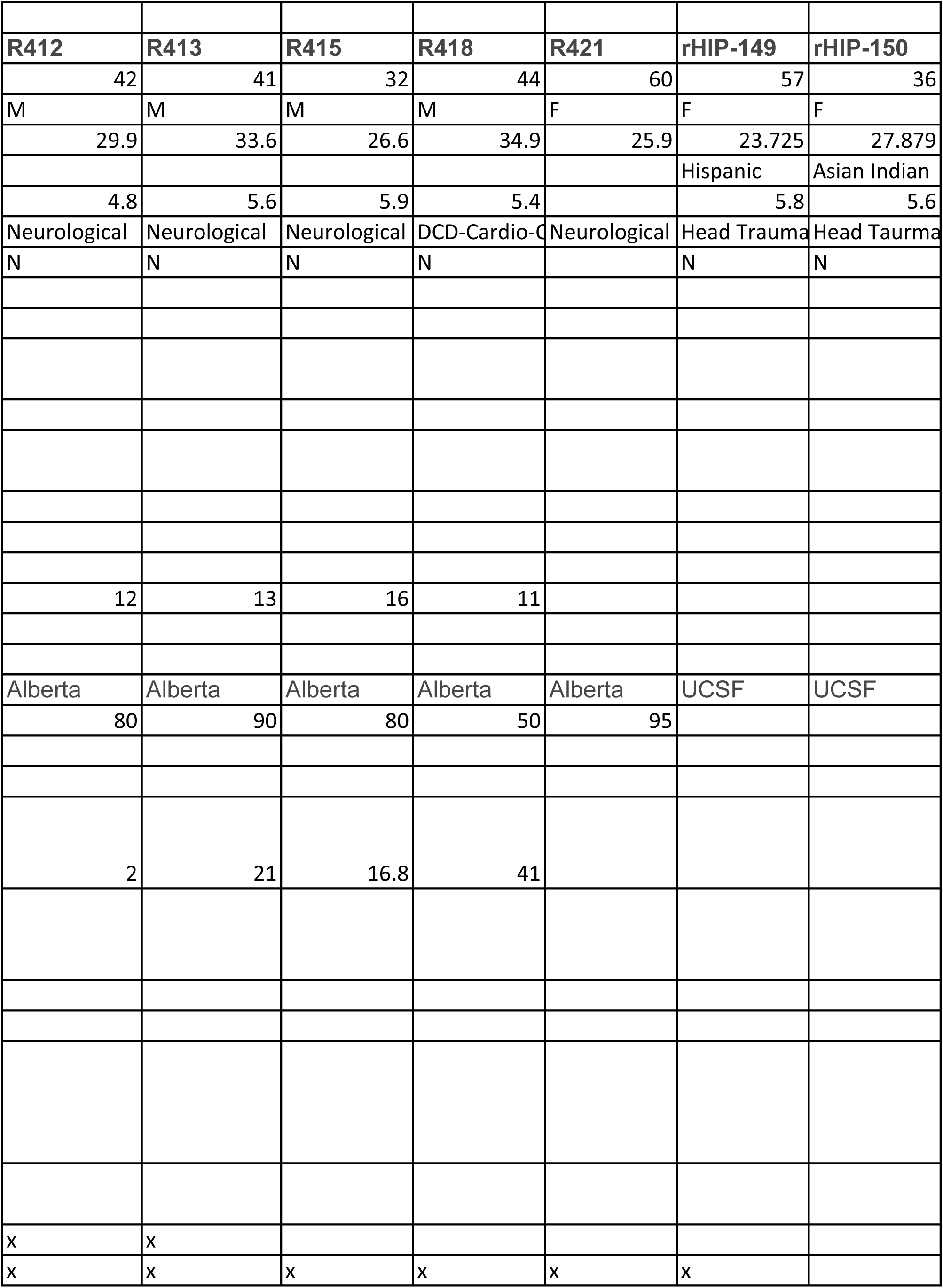
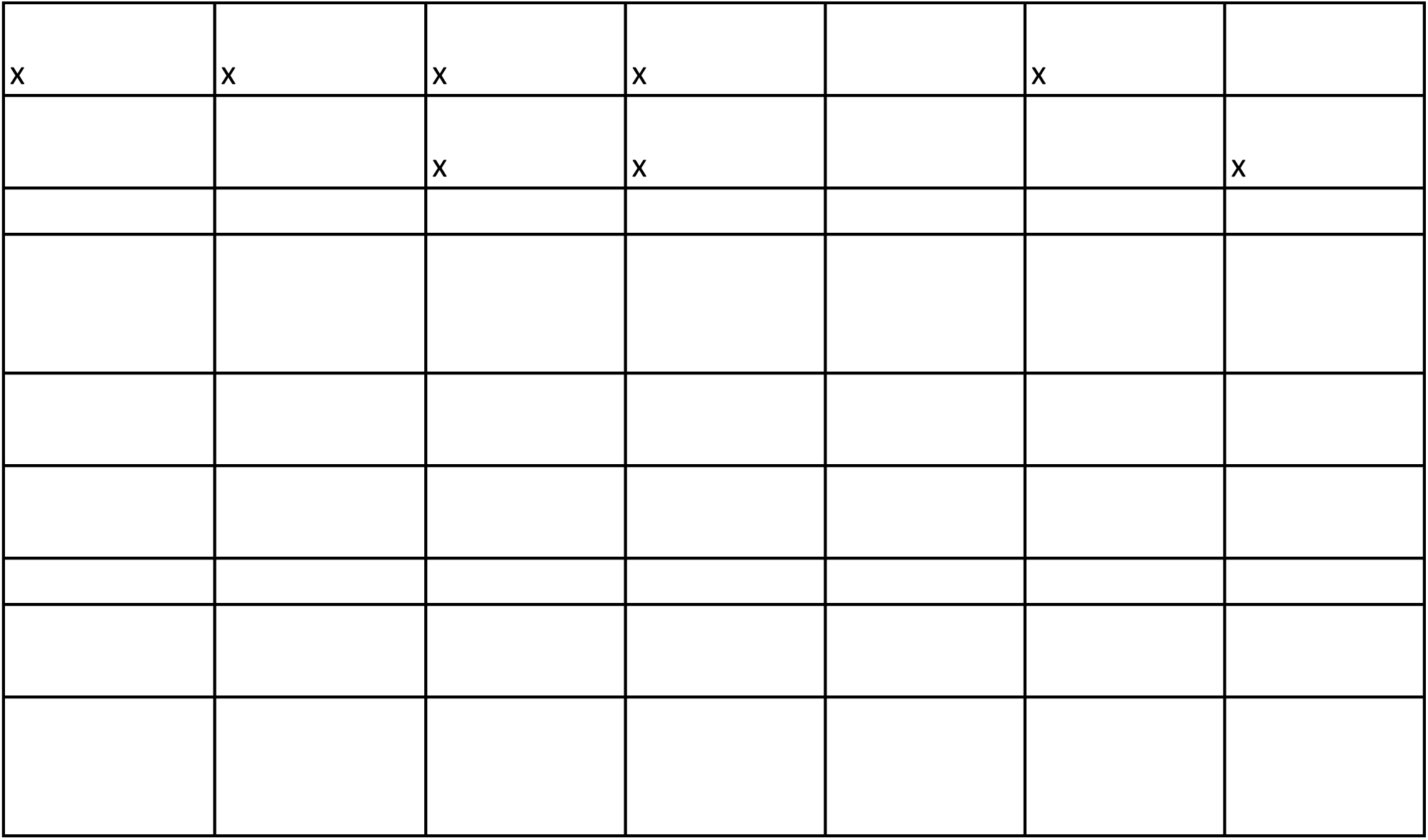
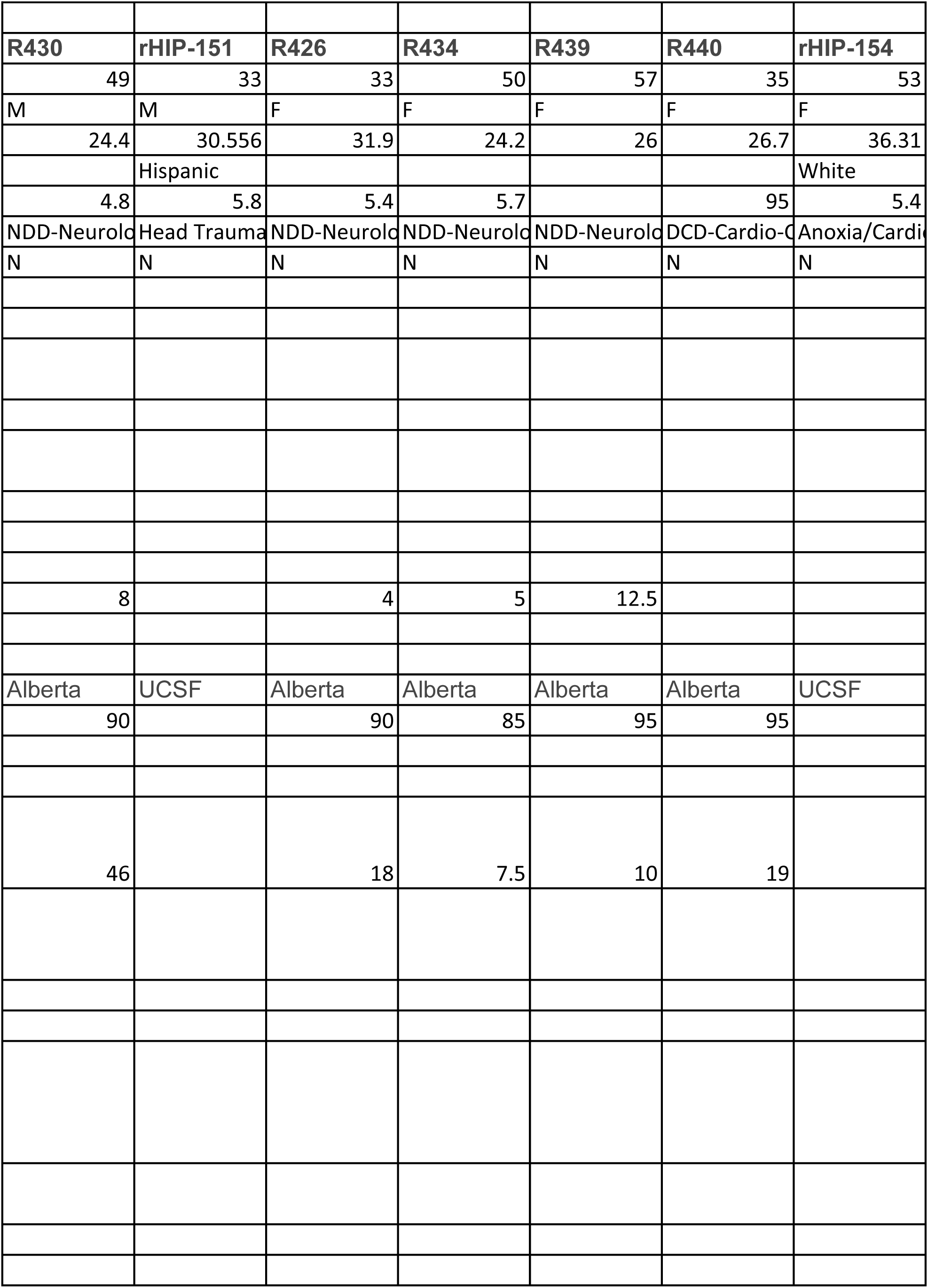
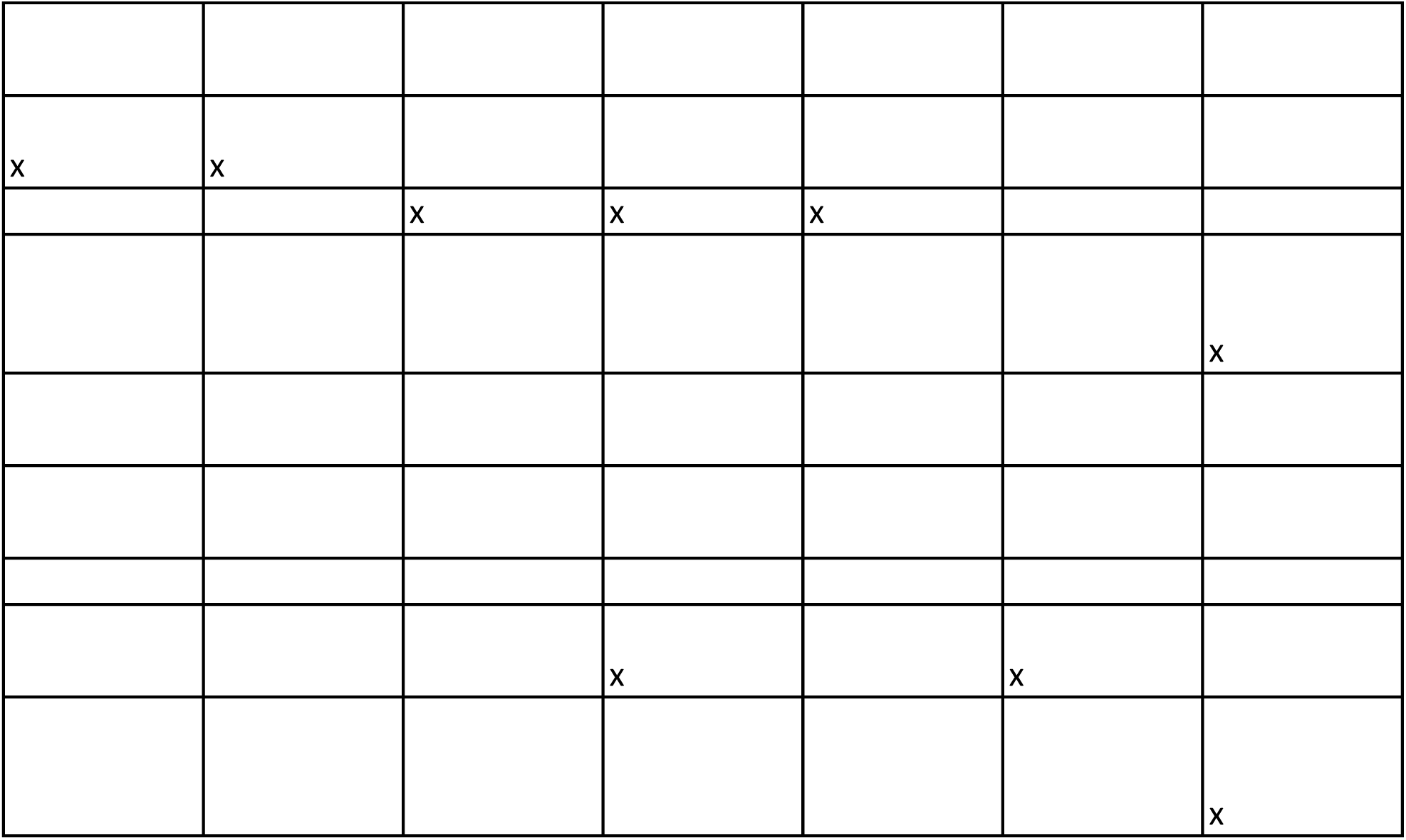
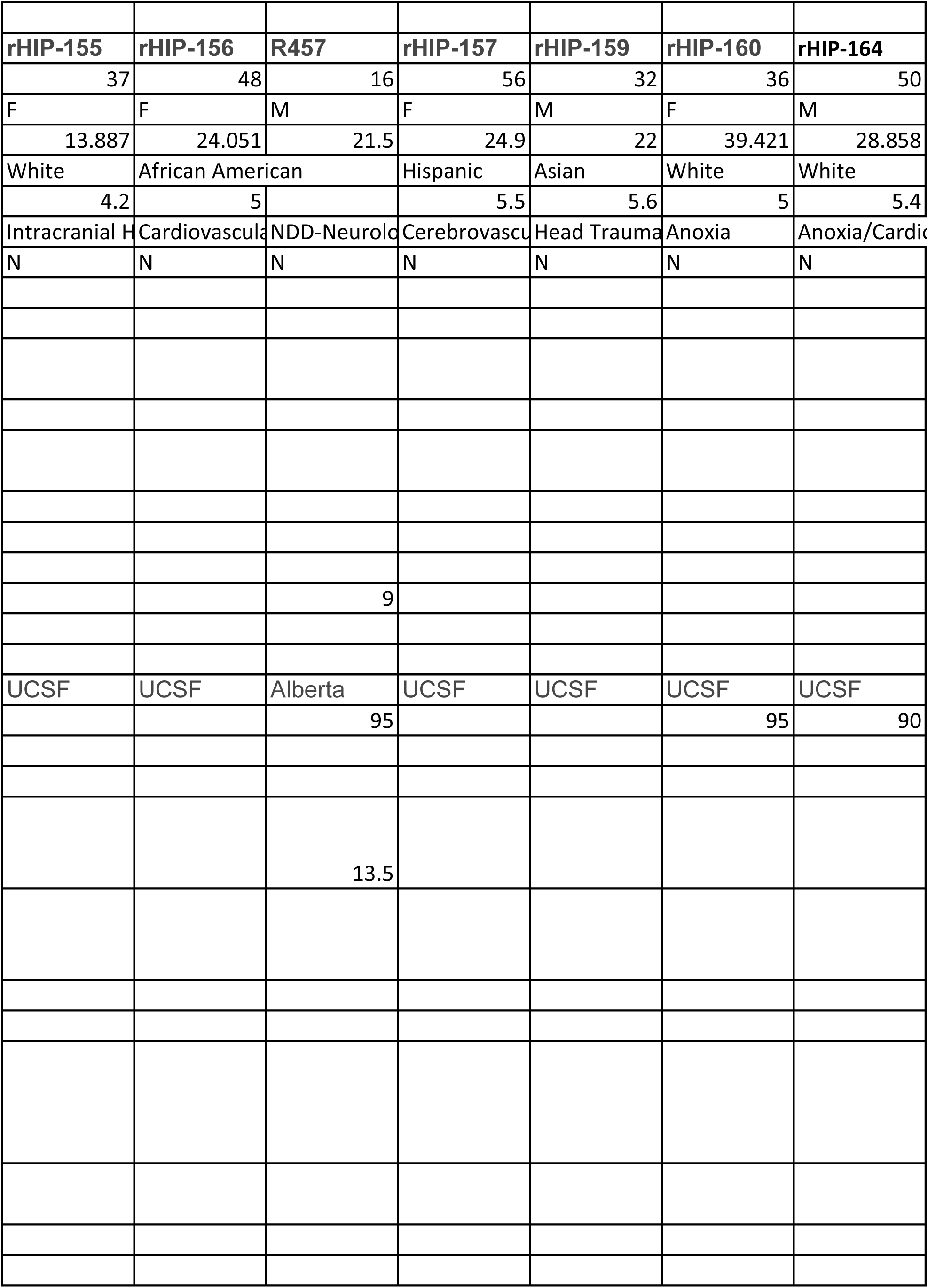
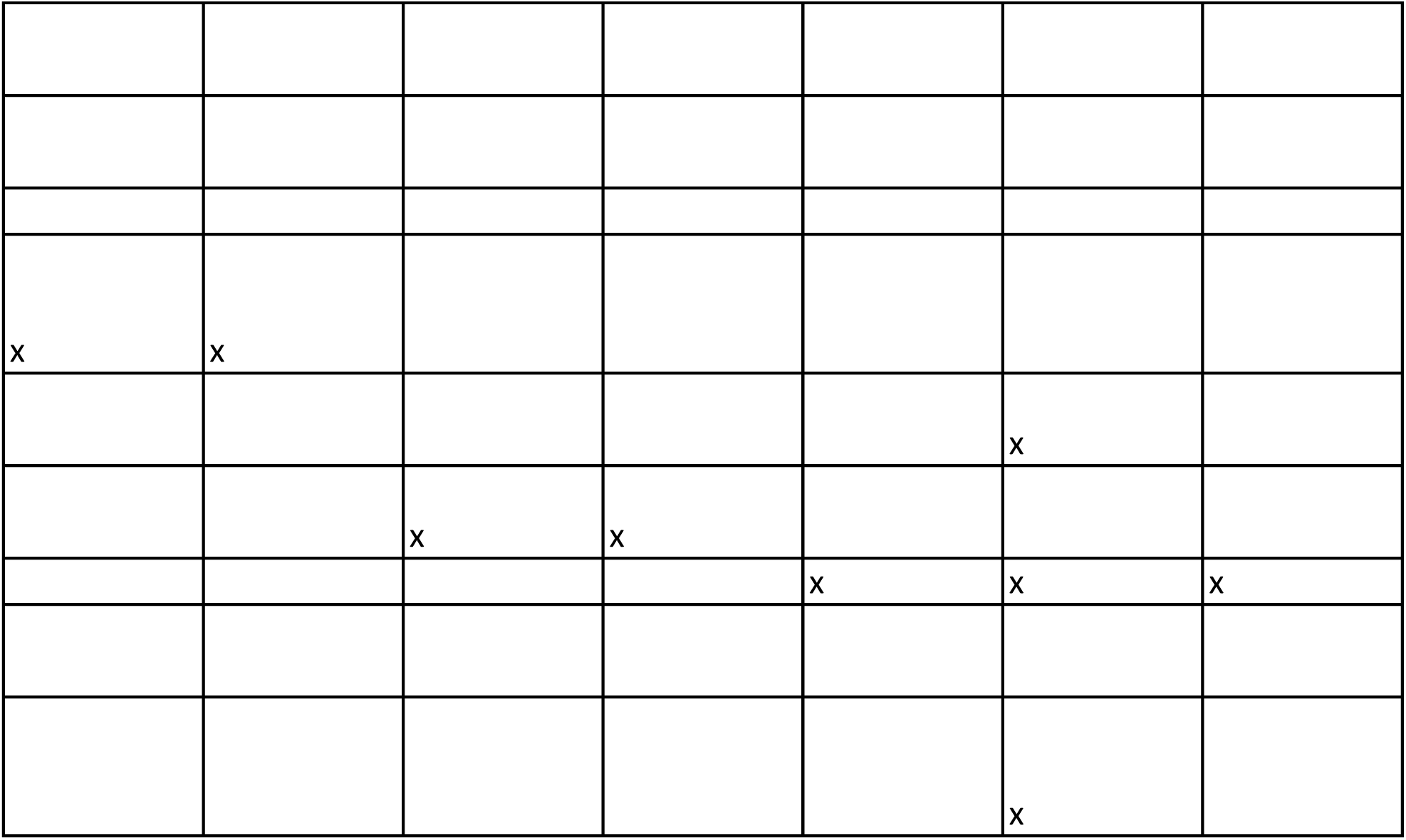
Donor Information.

**Data File S2.**
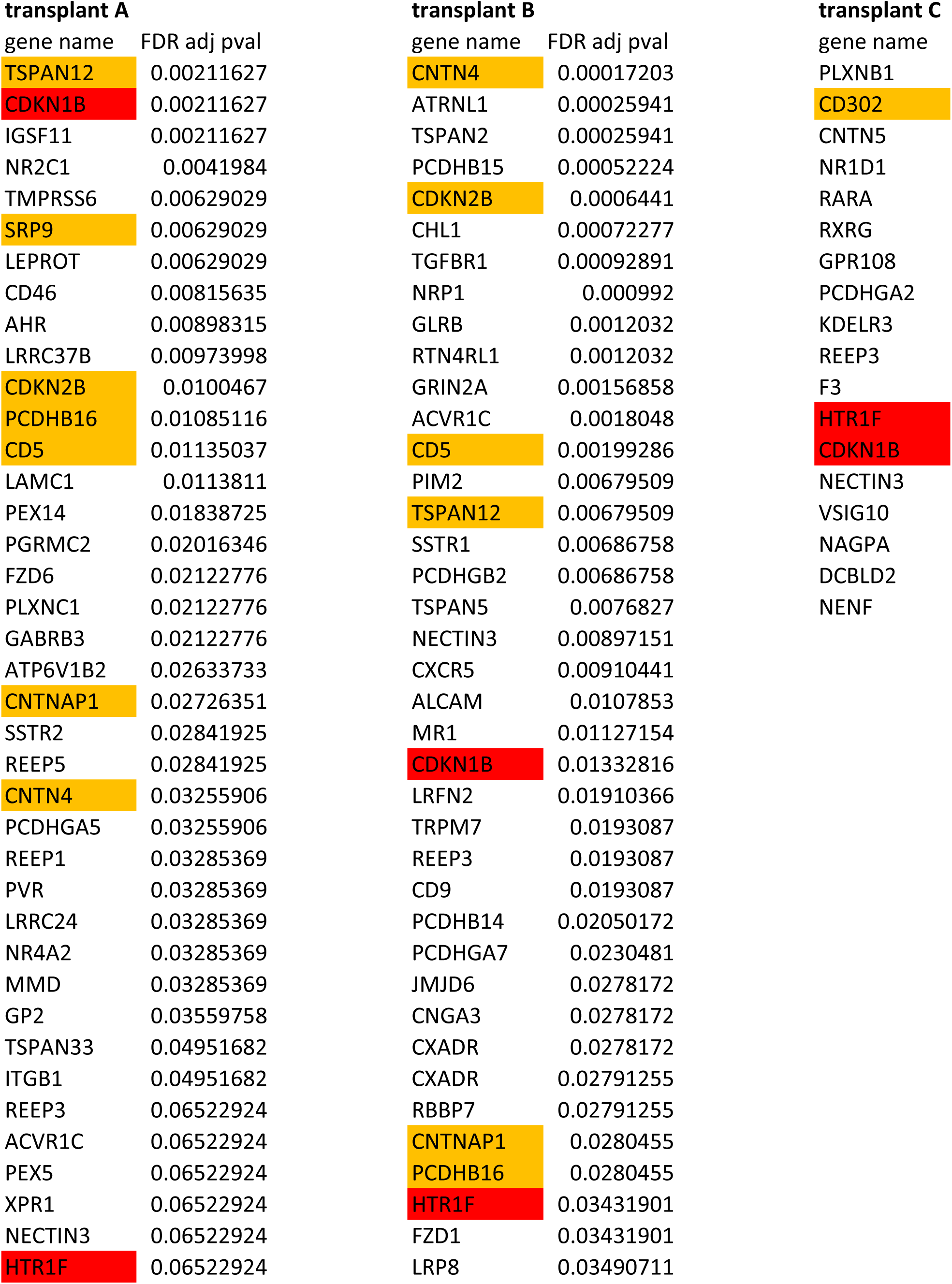
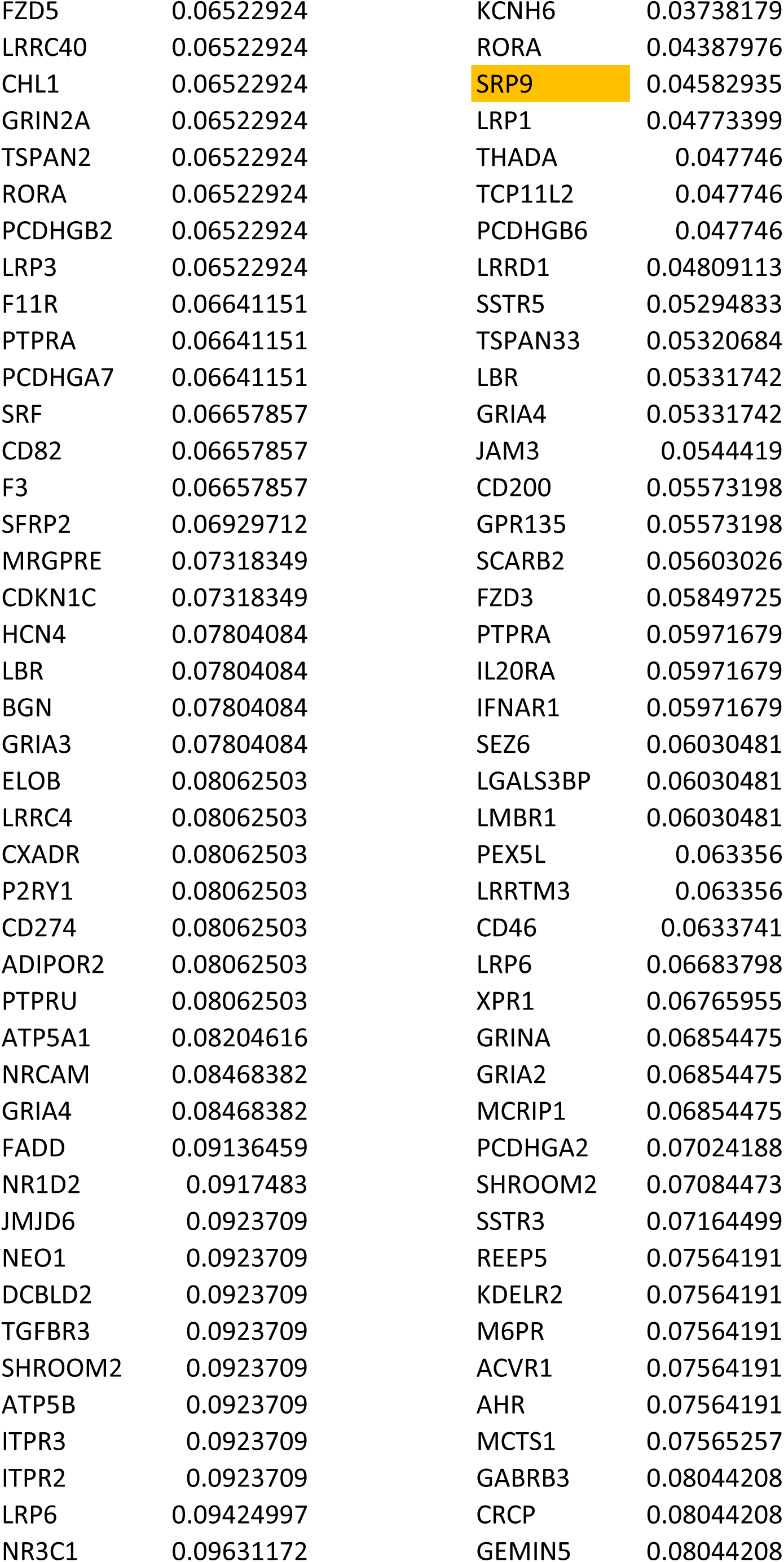
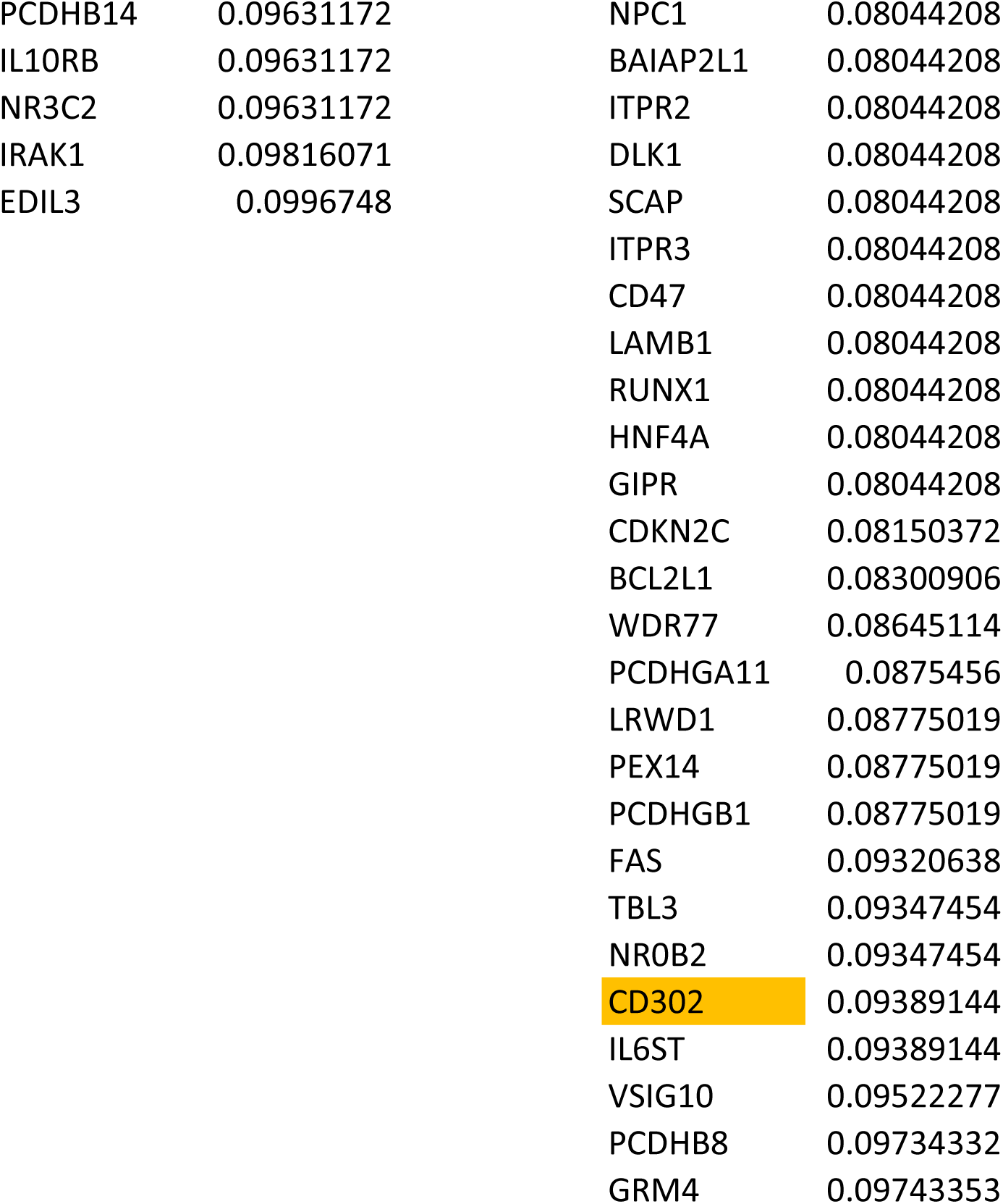
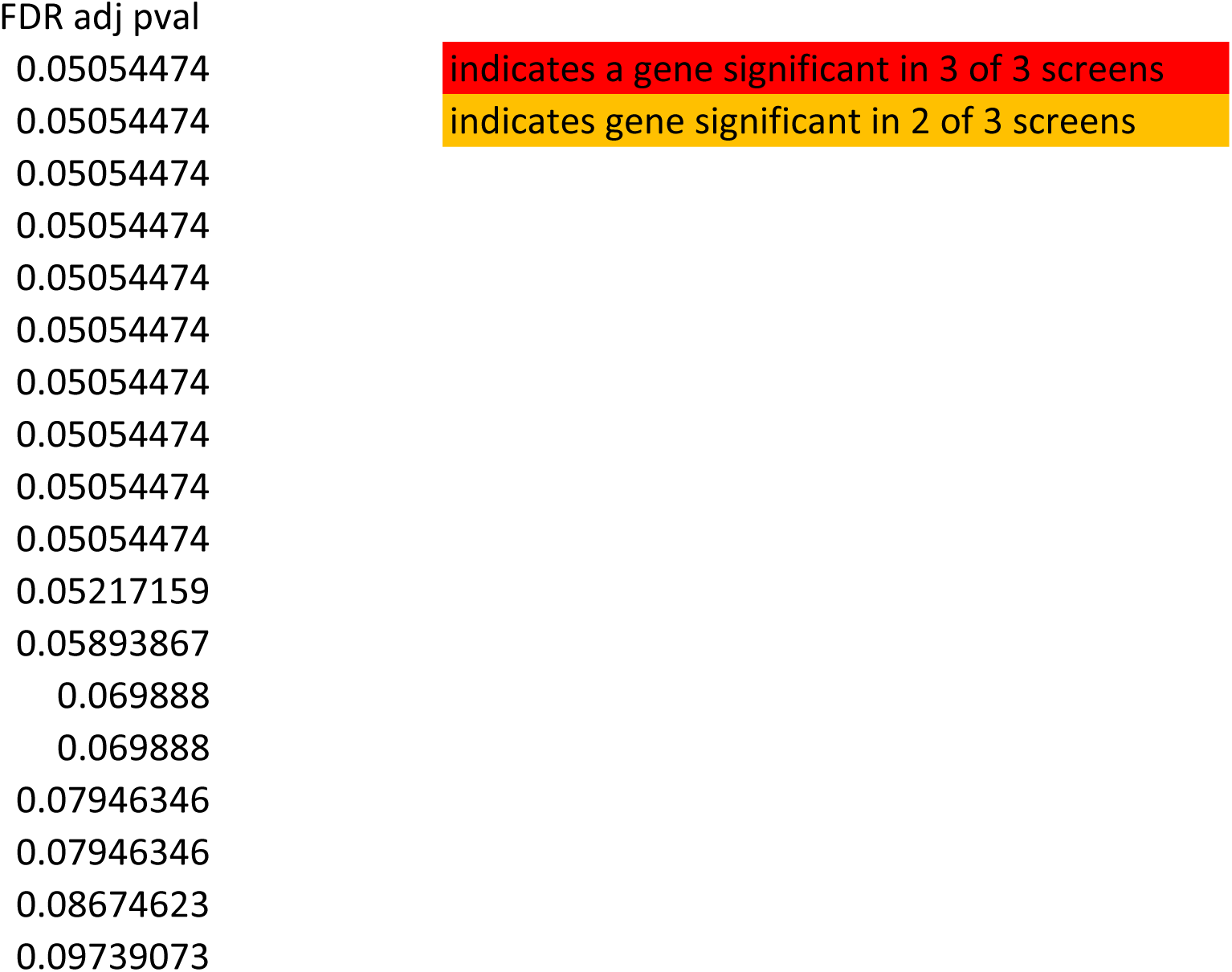
p-values for genes with shRNAs that become enriched after transplantation.

## References and Notes

1. M. C. Vantyghem, E. J. P. de Koning, F. Pattou, M. R. Rickels, Advances in beta-cell replacement therapy for the treatment of type 1 diabetes. Lancet 394, 1274–1285 (2019).

2. E. A. Ryan, B. W. Paty, P. A. Senior, D. Bigam, E. Alfadhli, N. M. Kneteman, J. R. Lakey, A. M. Shapiro, Five-year follow -up after clinical islet transplantation. Diabetes 54, 2060–2069 (2005).

3. G. Faleo, H. A. Russ, S. Wisel, A. V. Parent, V. Nguyen, G. G. Nair, J. E. Freise, K. E. Villanueva, G. L. Szot, M. Hebrok, Q. Tang, Mitigating Ischemic Injury of Stem Cell-Derived Insulin-Producing Cells after Transplant. Stem Cell Reports 9, 807–819 (2017).

4. A. E. Butler, J. Janson, S. Bonner-Weir, R. Ritzel, R. A. Rizza, P. C. Butler, Beta-cell deficit and increased beta-cell apoptosis in humans with type 2 diabetes. Diabetes 52, 102–110 (2003).

5. L. Baeyens, M. Lemper, W. Staels, S. De Groef, N. De Leu, Y. Heremans, M. S. German, H. Heimberg, (Re)generating Human Beta Cells: Status, Pitfalls, and Perspectives. Physiol Rev 98, 1143–1167 (2018).

6. A. F. Stewart, M. A. Hussain, A. Garcia-Ocana, R. C. Vasavada, A. Bhushan, E. Bernal-Mizrachi, R. N. Kulkarni, Human beta-cell proliferation and intracellular signaling: part 3. Diabetes 64, 1872–1885 (2015).

7. P. Wang, J. C. Alvarez-Perez, D. P. Felsenfeld, H. Liu, S. Sivendran, A. Bender, A. Kumar, R. Sanchez, D. K. Scott, A. Garcia-Ocana, A. F. Stewart, A high-throughput chemical screen reveals that harmine-mediated inhibition of DYRK1A increases human pancreatic beta cell replication. Nat Med 21, 383–388 (2015).

8. E. Dirice, D. Walpita, A. Vetere, B. C. Meier, S. Kahraman, J. Hu, V. Dancik, S. M. Burns, T. J. Gilbert, D. E. Olson, P. A. Clemons, R. N. Kulkarni, B. K. Wagner, Inhibition of DYRK1A Stimulates Human beta-Cell Proliferation. Diabetes 65, 1660–1671 (2016).

9. W. Shen, B. Taylor, Q. Jin, V. Nguyen-Tran, S. Meeusen, Y. Q. Zhang, A. Kamireddy, A. Swafford, A. F. Powers, J. Walker, J. Lamb, B. Bursalaya, M. DiDonato, G. Harb, M. Qiu, C. M. Filippi, L. Deaton, C. N. Turk, W. L. Suarez-Pinzon, Y. Liu, X. Hao, T. Mo, S. Yan, J. Li, A. E. Herman, B. J. Hering, T. Wu, H. Martin Seidel, P. McNamara, R. Glynne, B. Laffitte, Inhibition of DYRK1A and GSK3B induces human beta-cell proliferation. Nat Commun 6, 8372 (2015).

10. W. Shen, M. S. Tremblay, V. A. Deshmukh, W. Wang, C. M. Filippi, G. Harb, Y. Q. Zhang, A. Kamireddy, J. E. Baaten, Q. Jin, T. Wu, J. G. Swoboda, C. Y. Cho, J. Li, B. A. Laffitte, P. McNamara, R. Glynne, X. Wu, A. E. Herman, P. G. Schultz, Small-molecule inducer of beta cell proliferation identified by high-throughput screening. J Am Chem Soc 135, 1669–1672 (2013).

11. Y. Abdolazimi, Z. Zhao, S. Lee, H. Xu, P. Allegretti, T. M. Horton, B. Yeh, H. P. Moeller, R. J. Nichols, D. McCutcheon, A. Shalizi, M. Smith, N. A. Armstrong, J. P. Annes, CC-401 Promotes beta-Cell Replication via Pleiotropic Consequences of DYRK1A/B Inhibition. Endocrinology 159, 3143–3157 (2018).

12. J. P. Annes, J. H. Ryu, K. Lam, P. J. Carolan, K. Utz, J. Hollister-Lock, A. C. Arvanites, L. L. Rubin, G. Weir, D. A. Melton, Adenosine kinase inhibition selectively promotes rodent and porcine islet beta-cell replication. Proc Natl Acad Sci U S A 109, 3915–3920 (2012).

13. C. Dai, Y. Hang, A. Shostak, G. Poffenberger, N. Hart, N. Prasad, N. Phillips, S. E. Levy, D. L. Greiner, L. D. Shultz, R. Bottino, S. K. Kim, A. C. Powers, Age-dependent human beta cell proliferation induced by glucagon-like peptide 1 and calcineurin signaling. J Clin Invest 127, 3835–3844 (2017).

14. A. C. Nica, H. Ongen, J. C. Irminger, D. Bosco, T. Berney, S. E. Antonarakis, P. A. Halban, E. T. Dermitzakis, Cell-type, allelic, and genetic signatures in the human pancreatic beta cell transcriptome. Genome Res 23, 1554–1562 (2013).

15. H. Mi, A. Muruganujan, P. D. Thomas, PANTHER in 2013: modeling the evolution of gene function, and other gene attributes, in the context of phylogenetic trees. Nucleic Acids Res 41, D377–386 (2013).

16. J. Stein, W. M. Milewski, M. Hara, D. F. Steiner, A. Dey, GSK-3 inactivation or depletion promotes beta-cell replication via down regulation of the CDK inhibitor, p27 (Kip1). Islets 3, 21–34 (2011).

17. T. Uchida, T. Nakamura, N. Hashimoto, T. Matsuda, K. Kotani, H. Sakaue, Y. Kido, Y. Hayashi, K. I. Nakayama, M. F. White, M. Kasuga, Deletion of Cdkn1b ameliorates hyperglycemia by maintaining compensatory hyperinsulinemia in diabetic mice. Nat Med 11, 175–182 (2005).

18. N. Adham, H. T. Kao, L. E. Schecter, J. Bard, M. Olsen, D. Urquhart, M. Durkin, P. R. Hartig, R. L. Weinshank, T. A. Branchek, Cloning of another human serotonin receptor (5-HT1F): a fifth 5-HT1 receptor subtype coupled to the inhibition of adenylate cyclase. Proc Natl Acad Sci U S A 90, 408–412 (1993).

19. N. Amlaiky, S. Ramboz, U. Boschert, J. L. Plassat, R. Hen, Isolation of a mouse “5HT1E-like” serotonin receptor expressed predominantly in hippocampus. J Biol Chem 267, 19761–19764 (1992).

20. E. Gylfe, Association between 5-hydroxytryptamine release and insulin secretion. J Endocrinol 78, 239–248 (1978).

21. H. Kim, Y. Toyofuku, F. C. Lynn, E. Chak, T. Uchida, H. Mizukami, Y. Fujitani, R. Kawamori, T. Miyatsuka, Y. Kosaka, K. Yang, G. Honig, M. van der Hart, N. Kishimoto, J. Wang, S. Yagihashi, L. H. Tecott, H. Watada, M. S. German, Serotonin regulates pancreatic beta cell mass during pregnancy. Nat Med 16, 804–808 (2010).

22. J. H. Moon, H. Kim, H. Kim, J. Park, W. Choi, W. Choi, H. J. Hong, H. J. Ro, S. Jun, S. H. Choi, R. R. Banerjee, M. Shong, N. H. Cho, S. K. Kim, M. S. German, H. C. Jang, H. Kim, Lactation improves pancreatic beta cell mass and function through serotonin production. Sci Transl Med 12, (2020).

23. J. H. Moon, Y. G. Kim, K. Kim, S. Osonoi, S. Wang, D. C. Saunders, J. Wang, K. Yang, H. Kim, J. Lee, J. S. Jeong, R. R. Banerjee, S. K. Kim, Y. Wu, H. Mizukami, A. C. Powers, M. S. German, H. Kim, Serotonin Regulates Adult beta-Cell Mass by Stimulating Perinatal beta-Cell Proliferation. Diabetes 69, 205–214 (2020).

24. H. Bennet, I. G. Mollet, A. Balhuizen, A. Medina, C. Nagorny, A. Bagge, J. Fadista, E. Ottosson-Laakso, P. Vikman, M. Dekker-Nitert, L. Eliasson, N. Wierup, I. Artner, M. Fex, Serotonin (5-HT) receptor 2b activation augments glucose-stimulated insulin secretion in human and mouse islets of Langerhans. Diabetologia 59, 744–754 (2016).

25. M. Ohara-Imaizumi, H. Kim, M. Yoshida, T. Fujiwara, K. Aoyagi, Y. Toyofuku, Y. Nakamichi, C. Nishiwaki, T. Okamura, T. Uchida, Y. Fujitani, K. Akagawa, M. Kakei, H. Watada, M. S. German, S. Nagamatsu, Serotonin regulates glucose-stimulated insulin secretion from pancreatic beta cells during pregnancy. Proc Natl Acad Sci U S A 110, 19420–19425 (2013).

26. J. Almaca, J. Molina, D. Menegaz, A. N. Pronin, A. Tamayo, V. Slepak, P. O. Berggren, A. Caicedo, Human Beta Cells Produce and Release Serotonin to Inhibit Glucagon Secretion from Alpha Cells. Cell Rep 17, 3281–3291 (2016).

27. J. Fadista, P. Vikman, E. O. Laakso, I. G. Mollet, J. L. Esguerra, J. Taneera, P. Storm, P. Osmark, C. Ladenvall, R. B. Prasad, K. B. Hansson, F. Finotello, K. Uvebrant, J. K. Ofori, B. Di Camillo, U. Krus, C. M. Cilio, O. Hansson, L. Eliasson, A. H. Rosengren, E. Renstrom, C. B. Wollheim, L. Groop, Global genomic and transcriptomic analysis of human pancreatic islets reveals novel genes influencing glucose metabolism. Proc Natl Acad Sci U S A 111, 13924–13929 (2014).

28. N. Lawlor, J. George, M. Bolisetty, R. Kursawe, L. Sun, V. Sivakamasundari, I. Kycia, P. Robson, M. L. Stitzel, Single-cell transcriptomes identify human islet cell signatures and reveal cell-type-specific expression changes in type 2 diabetes. Genome Res 27, 208–222 (2017).

29. S. Korchynska, M. Krassnitzer, K. Malenczyk, R. B. Prasad, E. O. Tretiakov, S. Rehman, V. Cinquina, V. Gernedl, M. Farlik, J. Petersen, S. Hannes, J. Schachenhofer, S. N. Reisinger, A. Zambon, O. Asplund, I. Artner, E. Keimpema, G. Lubec, J. Mulder, C. Bock, D. D. Pollak, R. A. Romanov, C. Pifl, L. Groop, T. G. Hokfelt, T. Harkany, Life-long impairment of glucose homeostasis upon prenatal exposure to psychostimulants. EMBO J 39, e100882 (2020).

30. S. Amisten, P. Atanes, R. Hawkes, I. Ruz-Maldonado, B. Liu, F. Parandeh, M. Zhao, G. C. Huang, A. Salehi, S. J. Persaud, A comparative analysis of human and mouse islet G-protein coupled receptor expression. Sci Rep 7, 46600 (2017).

31. C. S. Rashid, Y. C. Lien, A. Bansal, L. J. Jaeckle-Santos, C. Li, K. J. Won, R. A. Simmons, Transcriptomic Analysis Reveals Novel Mechanisms Mediating Islet Dysfunction in the Intrauterine Growth-Restricted Rat. Endocrinology 159, 1035–1049 (2018).

32. C. Ackeifi, P. Wang, E. Karakose, J. E. Manning Fox, B. J. González, H. Liu, J. Wilson, E. Swartz, C. Berrouet, Y. Li, K. Kumar, P. E. MacDonald, R. Sanchez, B. Thorens, R. DeVita, D. Homann, D. Egli, D. K. Scott, A. Garcia-Ocaña, A. F. Stewart, GLP-1 receptor agonists synergize with DYRK1A inhibitors to potentiate functional human &#x3b2; cell regeneration. Science Translational Medicine 12, eaaw9996 (2020).

33. D. J. Koch, Phebus, L.A., Rocc, V.P., Sajdyk, T.J., in Google Patents, E. P. Office, Ed. (2000).

34. T. W. Lovenberg, M. G. Erlander, B. M. Baron, M. Racke, A. L. Slone, B. W. Siegel, C. M. Craft, J. E. Burns, P. E. Danielson, J. G. Sutcliffe, Molecular cloning and functional expression of 5-HT1E-like rat and human 5-hydroxytryptamine receptor genes. Proc Natl Acad Sci U S A 90, 2184–2188 (1993).

35. K. Shariati, Z. Pappalardo, D. G. Chopra, N. Yiv, R. Sheen, G. Ku, Selective monitoring of insulin secretion after CRISPR interference in intact pancreatic islets despite submaximal infection. Islets 12, 59–69 (2020).

36. T. Nagy, M. Kampmann, CRISPulator: a discrete simulation tool for pooled genetic screens. BMC Bioinformatics 18, 347 (2017).

37. M. E. Capozzi, J. B. Wait, J. Koech, A. N. Gordon, R. W. Coch, B. Svendsen, B. Finan, D. A. D’Alessio, J. E. Campbell, Glucagon lowers glycemia when beta-cells are active. JCI Insight 5, (2019).

38. M. C. Bassik, M. Kampmann, R. J. Lebbink, S. Wang, M. Y. Hein, I. Poser, J. Weibezahn, M. A. Horlbeck, S. Chen, M. Mann, A. A. Hyman, E. M. Leproust, M. T. McManus, J. S. Weissman, A systematic mammalian genetic interaction map reveals pathways underlying ricin susceptibility. Cell 152, 909–922 (2013).

39. C. Fellmann, T. Hoffmann, V. Sridhar, B. Hopfgartner, M. Muhar, M. Roth, D. Y. Lai, I. A. Barbosa, J. S. Kwon, Y. Guan, N. Sinha, J. Zuber, An optimized microRNA backbone for effective single-copy RNAi. Cell Rep 5, 1704–1713 (2013).

40. M. Pertea, D. Kim, G. M. Pertea, J. T. Leek, S. L. Salzberg, Transcript-level expression analysis of RNA-seq experiments with HISAT, StringTie and Ballgown. Nat Protoc 11, 1650–1667 (2016).

41. J. Besnard, G. F. Ruda, V. Setola, K. Abecassis, R. M. Rodriguiz, X. P. Huang, S. Norval, M. F. Sassano, A. I. Shin, L. A. Webster, F. R. Simeons, L. Stojanovski, A. Prat, N. G. Seidah, D. B. Constam, G. R. Bickerton, K. D. Read, W. C. Wetsel, I. H. Gilbert, B. L. Roth, A. L. Hopkins, Automated design of ligands to polypharmacological profiles. Nature 492, 215–220 (2012).

42. L. Salas-Estrada, D. Provasi, X. Qiu, H. U. Kaniskan, X. P. Huang, J. F. DiBerto, J. M. Lamim Ribeiro, J. Jin, B. L. Roth, M. Filizola, De Novo Design of kappa-Opioid Receptor Antagonists Using a Generative Deep-Learning Framework. J Chem Inf Model 63, 5056–5065 (2023).

43. S. L. Lewandowski, R. L. Cardone, H. R. Foster, T. Ho, E. Potapenko, C. Poudel, H. R. VanDeusen, S. M. Sdao, T. C. Alves, X. Zhao, M. E. Capozzi, A. H. de Souza, I. Jahan, C. J. Thomas, C. S. Nunemaker, D. B. Davis, J. E. Campbell, R. G. Kibbey, M. J. Merrins, Pyruvate Kinase Controls Signal Strength in the Insulin Secretory Pathway. Cell Metab 32, 736–750 e735 (2020).

44. D. G. Chopra, N. Yiv, T. G. Hennings, Y. Zhang, G. M. Ku, Deletion of Gpr27 in vivo reduces insulin mRNA but does not result in diabetes. Sci Rep 10, 5629 (2020).

45. J. Luo, Z. L. Deng, X. Luo, N. Tang, W. X. Song, J. Chen, K. A. Sharff, H. H. Luu, R. C. Haydon, K. W. Kinzler, B. Vogelstein, T. C. He, A protocol for rapid generation of recombinant adenoviruses using the AdEasy system. Nat Protoc 2, 1236–1247 (2007).

