## Supplemental Figures for "An shRNA screen in primary human beta cells identifies the serotonin 1F receptor as a negative regulator of survival during transplant"

**List of Supplementary Materials**

Fig. S1

Fig. S2

Fig. S3

Fig. S4

Fig. S5

Fig. S6

Fig. S7

Fig. S8

Data file S1: Donor information

Data file S2: p-values for screen

**Supplementary Materials**

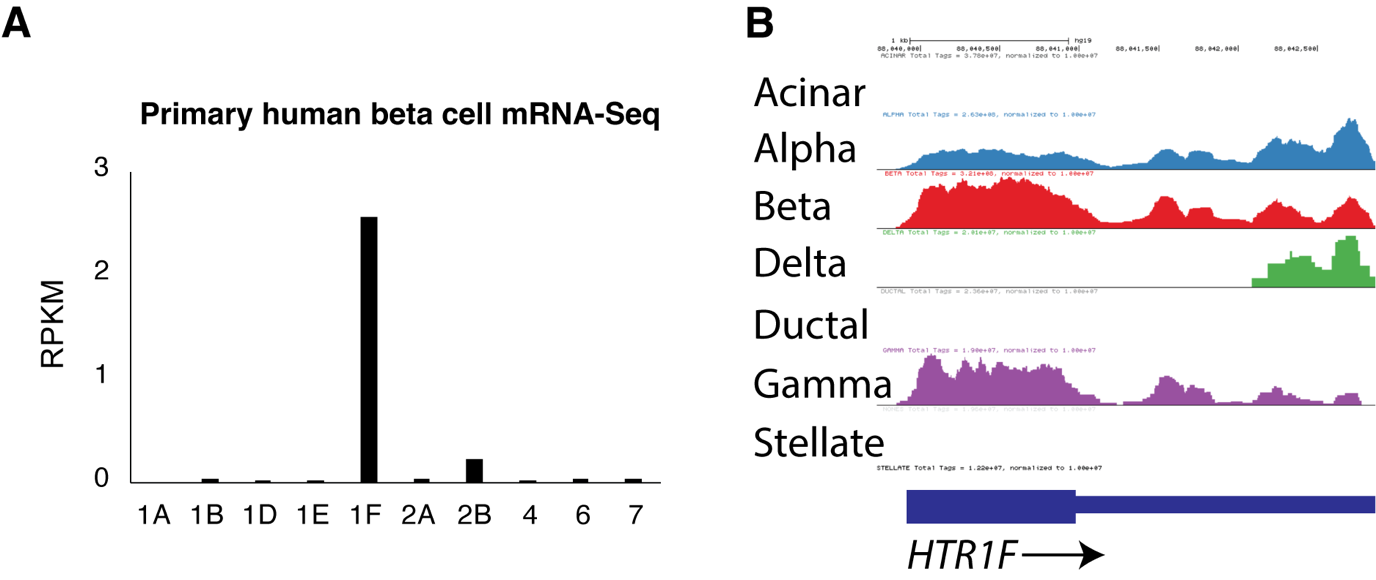

**Fig. S1. *5-HT_1F_* is the most highly expressed serotonin receptor in adult primary human beta cells. (A)** Expression of the indicated serotonin receptor gene from conventional mRNA-seq data from sorted primary human beta cells. Expression is reported in reads per kilobase per million (RPKM). Data replotted from(*14*). **(B)** Expression of 5-HT_1F_ in alpha, in human beta, delta, and gamma cells by aggregated single cell mRNA-seq. Replotted from (*28*).

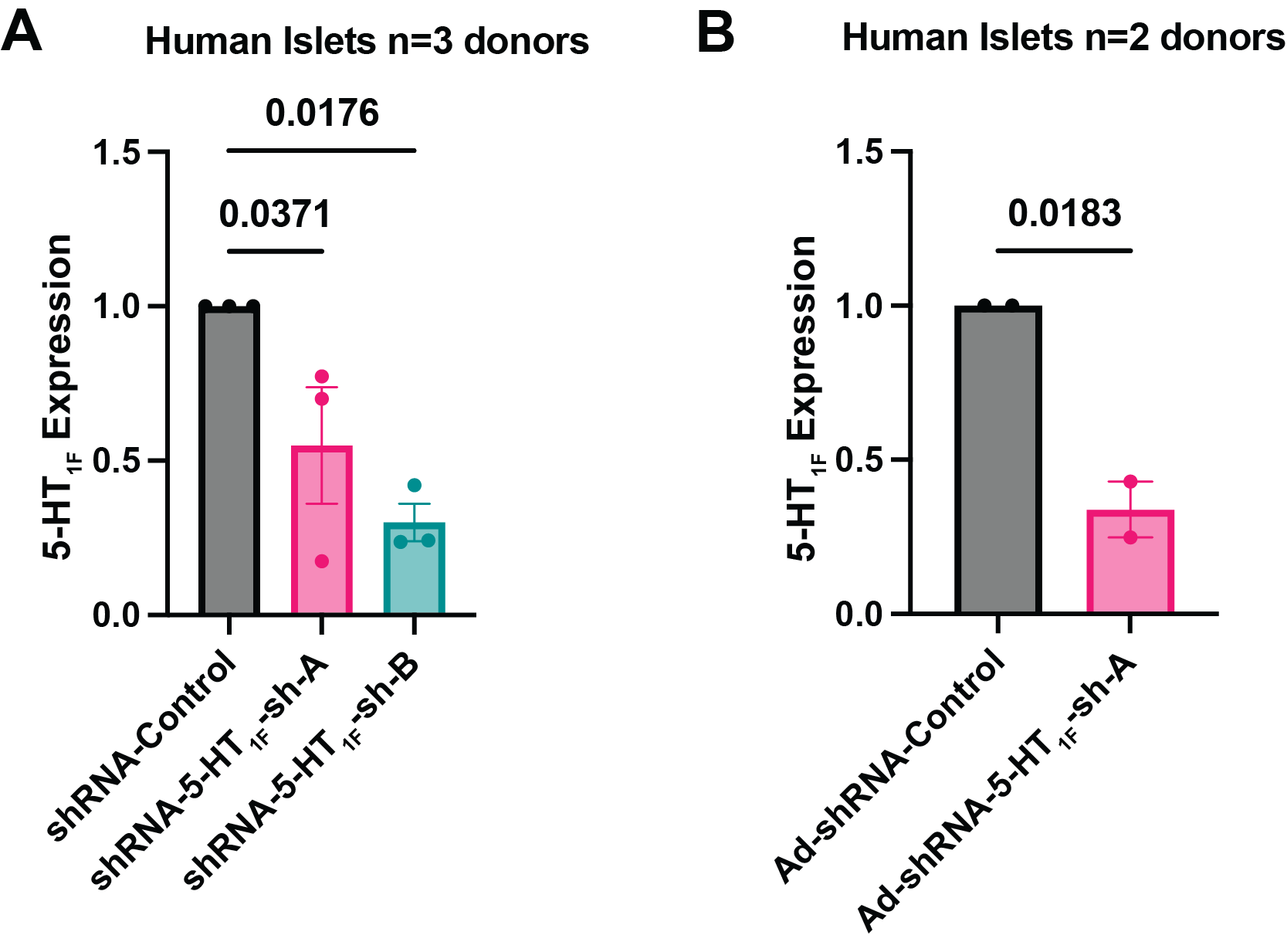

**Fig. S2. *5-HT_1F_* knockdown in human islets. (A)** 5-HT_1F_ mRNA normalized to TBP in human islets infected with lentivirus expressing shRNA a control, or two separate shRNAs targeting 5-HT_1F_: 5-HT_1F_ -sh-A or 5-HT_1F_ -sh-B from 3 independent donors with 3-4 replicates from each donor. P-values from a One-way ANOVA with Benjamini-Hochberg correction. (**B)** 5-HT_1F_ mRNA normalized to TBP in human islets infected with adenovirus expressing shRNA for control or 5-HT_1F_-sh-A from two independent donors. P-values from non-paired Student’s t-test. Error bars show standard error.

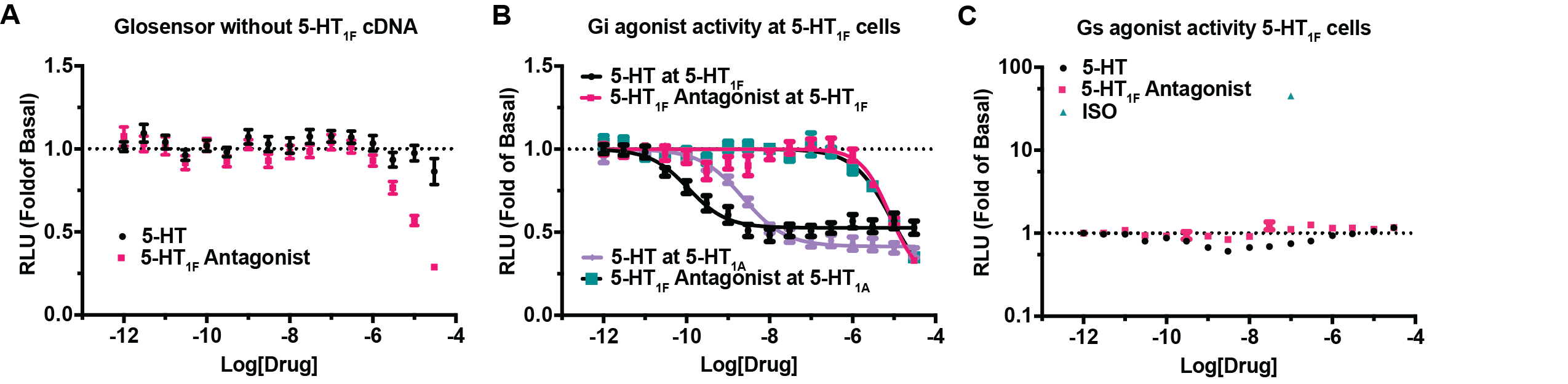

**Fig. S3. 1-(2-hydroxy-3-(naphthalen-2-yloxy)propyl)-4-(quinolin-3-yl)piperidin-4-ol is an 5-HT_1F_ antagonist. (A)** 293T cells transfected with the cAMP GloSensor plasmid (Promega) alone and treated with the indicated drug at the indicated concentration and then with 0.1μM isoproterenol prior to luminescence measurement, Error bars show standard error from n=4 independent assays, each in quadruplicate. **(B)** 293T cells were transiently transfected with the GloSensor plasmid (Promega) and the cDNA for 5-HT_1A_ or 5-HT_1F_ and were treated the indicated drugs at the indicated concentrations. Prior to luminescence measurement, the cells were treated with 0.1 μM isoproterenol. Error bars show standard error from n=4 independent assays, each in quadruplicate. **(C)** 293T cells transfected and treated as above without isoproterenol. Error bars show standard error from n=4 independent assays, each in quadruplicate.

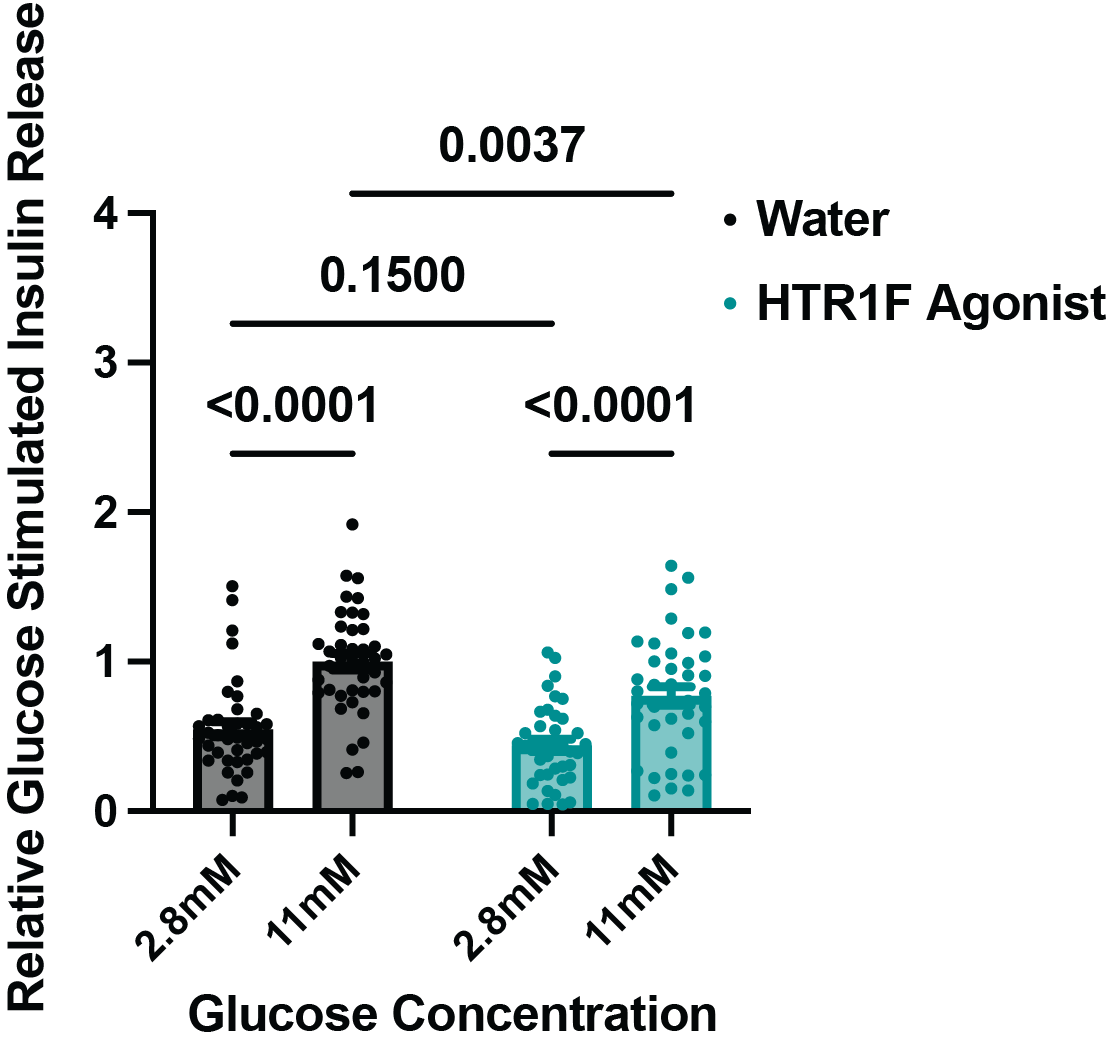

**Fig. S4. 5-HT_1F_ agonist inhibits glucose stimulated insulin secretion from human islets.** Human islets were pre-incubated with vehicle (water) or 150 nM 5-HT_1F_ agonist (LY344864) for 1 hour and during each hour of stimulation at 2.8 mM glucose and 11 mM glucose. n=8 independent donors, 5 replicates for each donor. Two-way ANOVA: drug effect p=0.0017, glucose effect p<0.0001. Interaction p=0.25. Post-hoc tests with Benjamini-Hochberg correction. Error bars show standard error.

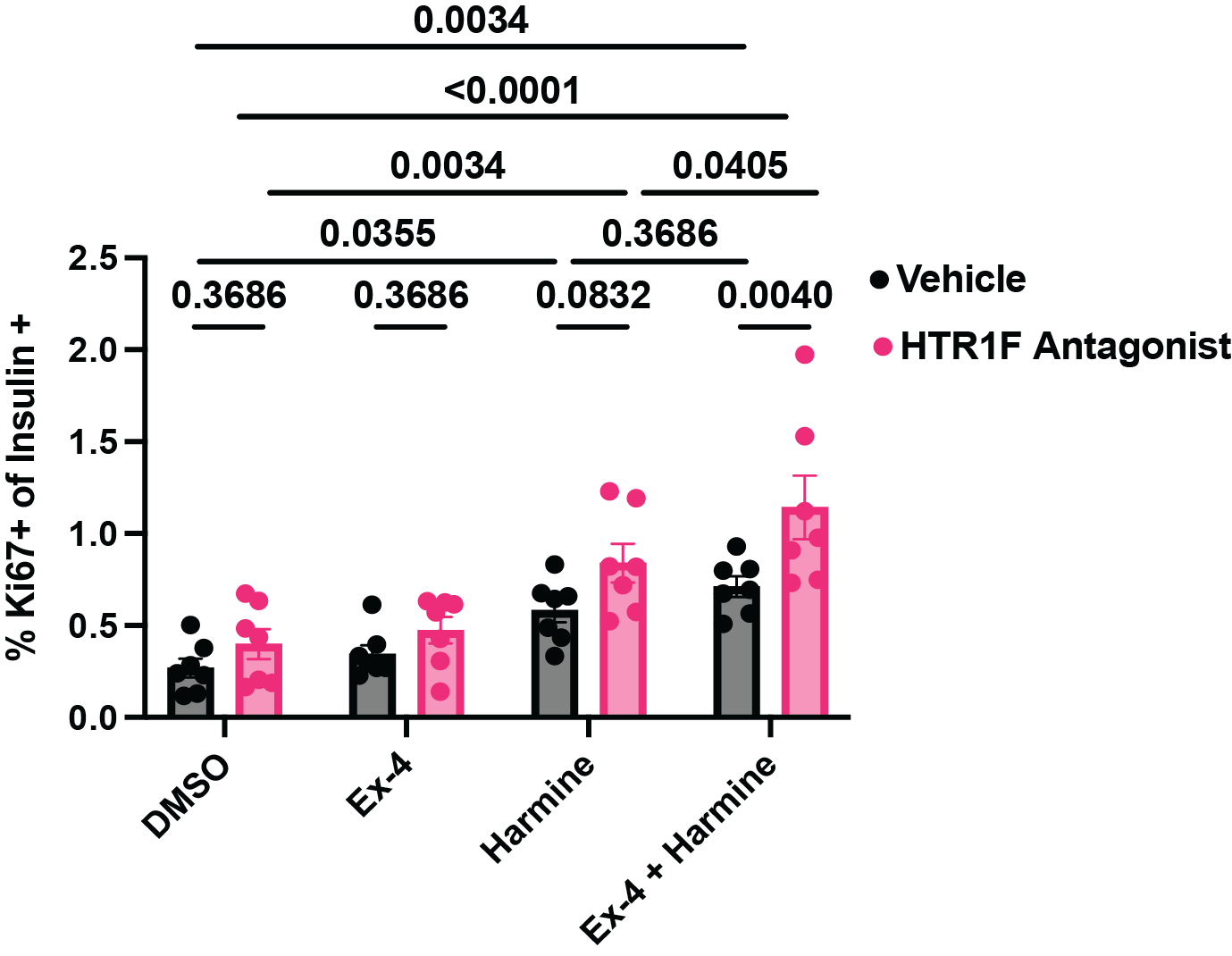

**Fig. S5. 5-HT_1F_ antagonist increases human beta cell proliferation but only in the presence of harmine and exendin-4.** Dissociated human islets were pretreated with 200nM 1-(2-hydroxy-3-(naphthalen-2-yloxy)propyl)-4-(quinolin-3-yl)piperidin-4-ol or vehicle for 24 hours and then treated with the indicated drugs for an additional 48-72 hours. n=7 donors. 2-way ANOVA: p=0.0005 for 1-(2-hydroxy-3-(naphthalen-2-yloxy)propyl)-4-(quinolin-3-yl)piperidin-4-ol effect, p<0.0001 for drug effect, p=0.3002 for interaction. Post-hoc testing with Benjamini-Hochberg correction. Only p-values <0.05 are shown for clarity (out of 28 tests). Error bars show standard error.

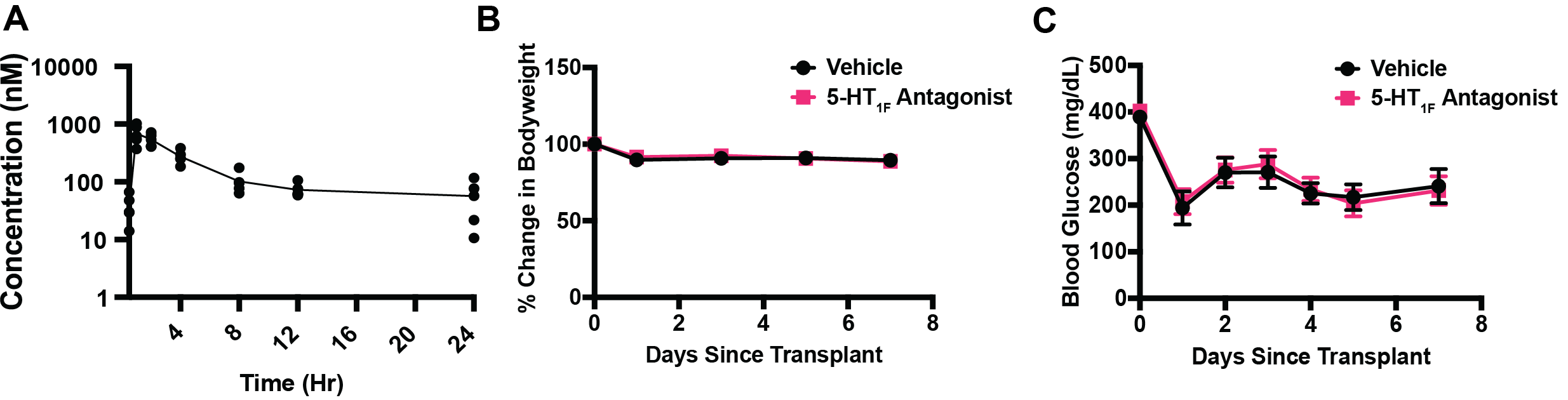

**Fig. S6.** **5-HT_1F_ antagonist in NSG mice. (A)** Plasma levels of 1-(2-hydroxy-3-(naphthalen-2-yloxy)propyl)-4-(quinolin-3-yl)piperidin-4-ol in mice after a single 20 mg/kg IP injection. n=5 mice. **(B)** Bodyweights of diabetic NSG mice who received human islet grafts treated with vehicle n=16 or 5-HT_1F_ antagonist n=15 from 3 independent donors. Two-way ANOVA: drug effect p=0.9631, time p<0.0001, interaction p=0.9414. **(C)** Random blood glucose levels. of diabetic NSG mice who received human islet grafts treated with vehicle n=16 or 5-HT_1F_ antagonist n=15 from 3 independent donors. Two-way ANOVA: drug effect p=0.7722, time p<0.0001, interaction p=0.9252. Error bars show standard error.

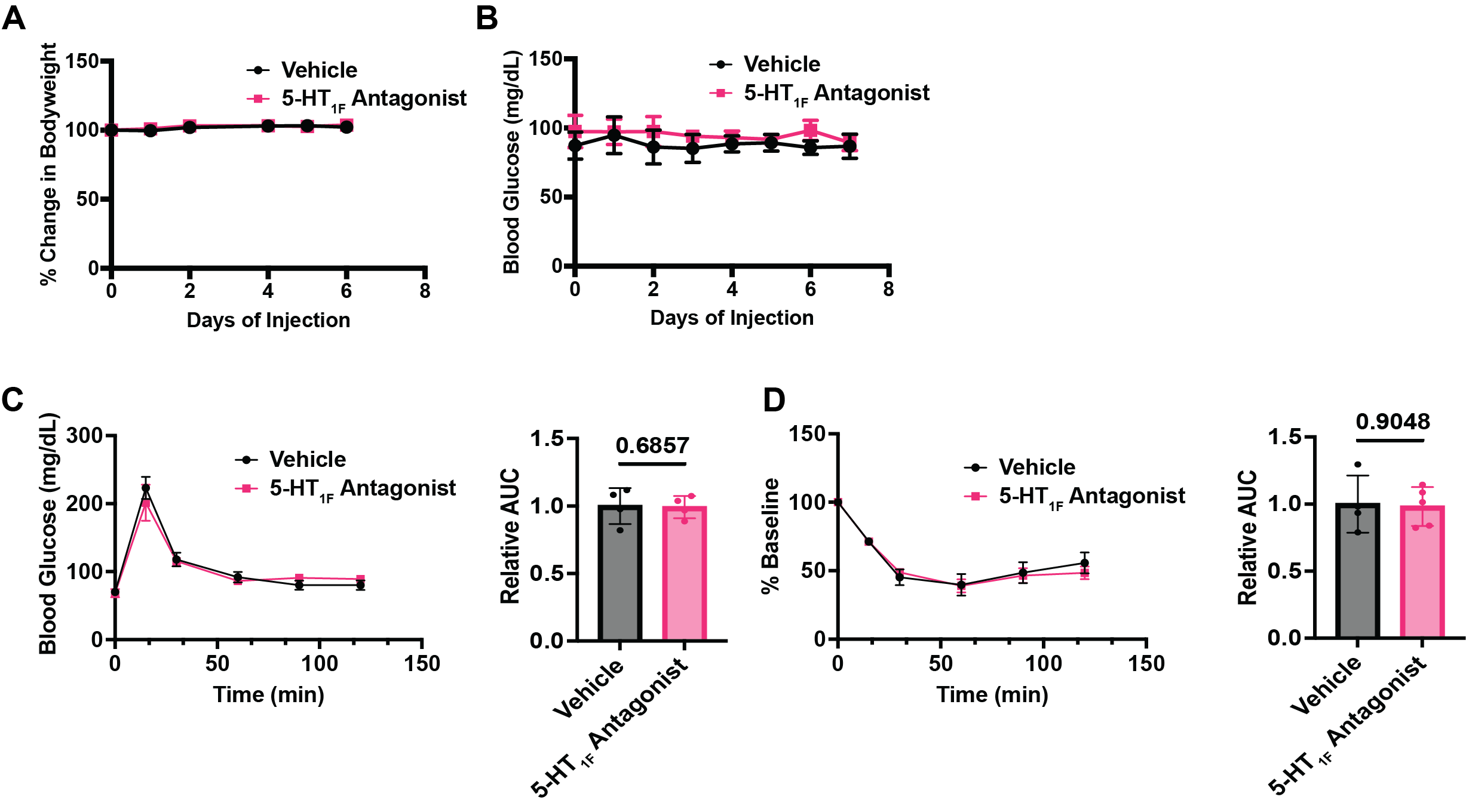

**Fig. S7. 5-HT_1F_ antagonist in NSG mice without human islet transplant. (A)** Bodyweight of non-transplanted NSG mice during a 7-day course of treatment with vehicle or HT_1F_ antagonist (20 mg/kg injected twice a day). Vehicle n=4, 5-HT_1F_ antagonist n=5. Two-way ANOVA: drug effect p=0.9156, time p=0.0109, interaction p=0.9156. **(B)** As in B, but non-fasting blood glucose is shown. Vehicle n=4, 5-HT_1F_ antagonist n=5. Two-way ANOVA: drug effect p=0.3075, time p=0.9879, interaction p=0.9931. **(C)** IPGTT on the mice from (**A**) at day 7. Vehicle n=4, 5-HT_1F_ antagonist n=4. Two-way ANOVA: drug effect p=0.8224, time p<0.0001, interaction p=0.6099. Relative area under the curve was calculated, p=0.6857 Mann-Whitney test. (**D**) Intraperitoneal insulin tolerance test on the mice from (B) at day 8. Vehicle n=4, 5-HT_1F_ antagonist n=5. Two-way ANOVA: drug effect p=0.8367, time p<0.0001, interaction p=0.7264. Relative area under the curve was calculated, p=0.9048 Mann-Whitney test. Error bars show standard error.

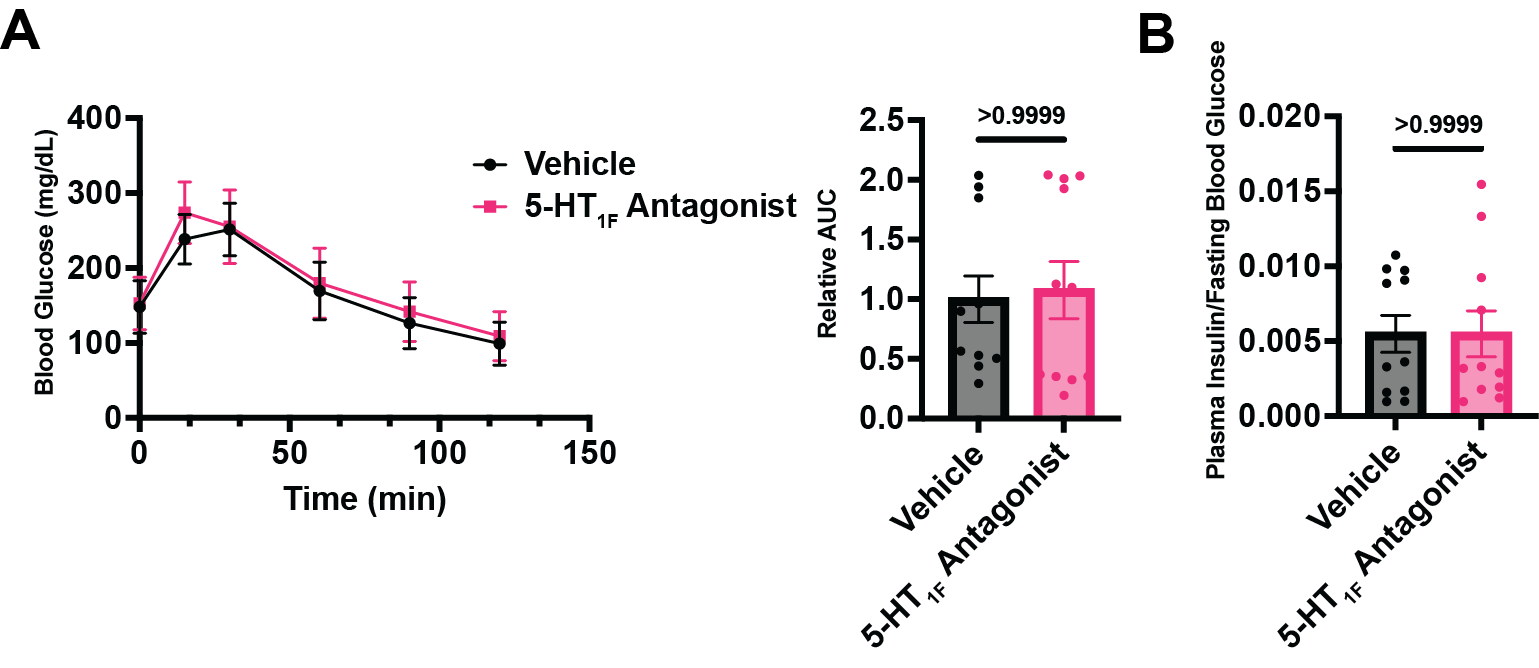

**Fig. S8. No effect of 5-HT_1F_ antagonist on glycemia or insulin in mice with stable human islet engraftment. (A)** IPGTT in diabetic NSG mice with human islets stably engrafted IP injected with vehicle or 1-(2-hydroxy-3-(naphthalen-2-yloxy)propyl)-4-(quinolin-3-yl)piperidin-4-ol 2 hours before the GTT. Two-way ANOVA: drug effect p=0.7998, time p<0.0001, interaction p=0.7590. Relative area under the curve was calculated, p>0.9999 Mann-Whitney test. **(B)** Ratio of fasting plasma insulin/fasting blood glucose. Vehicle n=11, 5-HT_1F_ antagonist n=11 from two independent donors, p>0.9999 Mann-Whitney test. Error bars show standard error.
